# Mathematical Modeling of Late-Stage LC Aggregation and Cardiac Injury Following Establishment of a Pathogenic Plasma-Cell Clone

**DOI:** 10.1101/2025.10.30.685705

**Authors:** Andrey V. Kuznetsov

## Abstract

AL amyloidosis is a rapidly progressive disorder characterized by clonal plasma cell expansion, excessive production of light chains (LCs), and their misfolding into aggregation-prone monomers. These monomers assemble into oligomers and ultimately deposit as amyloid fibrils, particularly within cardiac tissue, where they contribute to myocardial stiffening and direct cardiotoxicity. A reduced-order mechanistic model is developed to describe LC secretion by a pathogenic plasma-cell clone, LC unfolding and aggregation, cardiac deposition, and the resulting myocardial injury during advanced cardiac AL amyloidosis. Simulations reveal pronounced nonlinear LC aggregation kinetics: oligomer concentrations remain low during the early part of the modeled terminal cardiac-progression interval and subsequently increase rapidly as autocatalytic conversion becomes dominant. When aggregation is assumed to occur within cardiac tissue, fibril deposition is approximately 30 times greater, and oligomer-induced cardiotoxicity is about five times higher, compared with aggregation occurring in the blood plasma. These differences stem from the smaller cardiac volume, which accelerates autocatalytic oligomer formation. A combined cardiac damage criterion, integrating both oligomer-induced cardiotoxicity and fibril-associated myocardial stiffening, was introduced and found to reach values approximately tenfold higher when LC aggregation occurs within cardiac tissue compared with aggregation in the blood plasma. This parameter may provide a candidate model-based measure of cardiac aging or disease severity. The model also predicts that therapeutic intervention markedly reduces, but does not eliminate cardiac injury, highlighting the importance of early treatment initiation in AL amyloidosis.

## 1. Introduction

Immunoglobulin (Ig) light chain (AL) amyloidosis is the most prevalent form of systemic amyloidosis in Western countries, accounting for approximately 75% of cases (Rognoni et al. 2021; Kazman et al. 2021; Martinez-Rivas et al. 2022; Palladini and Merlini 2016). AL amyloidosis is characterized by misfolding of monoclonal Ig light chains (LCs) secreted by clonal plasma cells. These misfolded LCs demonstrate structural instability and a marked propensity to self-aggregate, forming soluble oligomers and insoluble fibrils (Eisele et al. 2015; Blancas-Mejía and Ramirez-Alvarado 2013). Progressive interstitial infiltration by amyloid fibrils compromises organ architecture and function, most prominently in the heart and kidneys, leading to restrictive cardiomyopathy, nephrotic syndrome, and ultimately multi-organ failure (Rognoni et al. 2021).

Cardiac involvement in AL amyloidosis is especially detrimental, presenting with restrictive cardiomyopathy, diastolic dysfunction, and arrhythmias, and remains the principal determinant of prognosis (Dispenzieri et al. 2003; Kumar et al. 2012). Once heart failure develops, median survival is limited to only a few months, underscoring the aggressive course of the disease (Dubrey et al. 1998).

Current therapeutic strategies in AL amyloidosis are largely adapted from multiple myeloma and focus on targeting the underlying plasma cell clone to suppress the production of pathogenic LCs (Palladini and Merlini 2016). Despite advances with chemotherapy, stem cell transplantation, proteasome inhibitors, immunomodulatory agents, and monoclonal antibodies, outcomes remain poor in patients with advanced cardiac involvement, highlighting a critical unmet need (Palladini et al. 2020; Kastritis Efstathios et al. 2021). This partly stems from a lack of detailed quantitative understanding of the processes that govern LC production, clearance, misfolding, and deposition within tissues. Current clinical approaches rely on indirect biomarkers, such as serum free LC assays and cardiac biomarkers (NT-proBNP, troponins), which provide prognostic information but do not capture the underlying dynamic interactions that drive disease progression (Dispenzieri et al. 2004; Katzmann et al. 2002).

Experimental studies have provided substantial insight into the molecular determinants of LC amyloidogenesis (Blancas-Mejia et al. 2018; Kazman et al. 2021; Pradhan et al. 2023). Amyloidogenic LCs are characterized by sequence-dependent differences in conformational stability, unfolding kinetics, and aggregation propensity (Blancas-Mejía, Hammernik, et al. 2015; Blancas-Mejia et al. 2018; Morgan et al. 2019). In particular, studies of patient-derived LC proteins have demonstrated that destabilization of the native state and subsequent formation of aggregation-prone conformations are important determinants of amyloid formation (Morgan et al. 2019; Kazman et al. 2021; Pradhan et al. 2023). Experimental investigations have further demonstrated that LC fibril formation is strongly influenced by the conformational state of the precursor and can be accelerated by pre-existing fibrillar aggregates, or seeds, supporting an important role for autocatalytic processes in LC amyloidogenesis (Blancas-Mejía and Ramirez-Alvarado 2016; Kazman et al. 2021).

Mathematical descriptions of protein aggregation have developed along several complementary lines (Morris et al. 2008). Early kinetic models represented amyloid formation as nucleation followed by fibril elongation, whereas subsequent models incorporated additional processes such as secondary nucleation and fibril fragmentation (Knowles et al. 2009; Cohen et al. 2013). The Finke–Watzky (F–W) two-step model provides a particularly compact representation of aggregation through an initial nucleation step followed by autocatalytic conversion of monomeric species into aggregates (Watzky and Finke 1997; Morris et al. 2008). Its minimal structure has allowed it to be applied to a range of protein and peptide aggregation systems while retaining a small number of kinetic parameters (Morris et al. 2008; Watzky et al. 2008).

Mechanistic kinetic frameworks have been developed to describe amyloid-β and other amyloidogenic proteins (Knowles et al. 2009; Meisl et al. 2016; Michaels et al. 2018). These frameworks can distinguish among microscopic processes including primary nucleation, fibril elongation, fibril fragmentation, and fibril-surface-catalyzed secondary nucleation, with the relative importance of these pathways depending on the protein and experimental conditions (Cohen et al. 2013; Meisl et al. 2014). Extensions of these approaches have also incorporated interactions with other molecular species, including molecular chaperones (Arosio et al. 2016). Such kinetic frameworks have been particularly successful in explaining the concentration dependence and strongly nonlinear aggregation kinetics of amyloid-β peptides and in identifying the dominant microscopic mechanisms responsible for aggregate proliferation (Cohen et al. 2013; Meisl et al. 2014).

Mathematical modeling has provided important insights into several processes relevant to AL amyloidosis. Models of hematologic malignancies have been used to investigate clonal plasma-cell kinetics, disease progression, and therapeutic resistance (Sullivan and Salmon 1972; Ayati et al. 2010). In parallel, quantitative models of amyloid fibrillogenesis have helped elucidate the kinetics of nucleation, fibril elongation, and secondary aggregation processes that govern the accumulation of amyloid burden (Watzky and Finke 1997; Knowles et al. 2009; Cohen et al. 2013). However, these two areas have largely been modeled separately. Although individual aspects of LC production, misfolding, aggregation, tissue deposition, and organ injury have been investigated (Takahashi et al. 2002; Hall and Edskes 2012; Sidana et al. 2020), considerably less attention has been devoted to mathematical frameworks that connect these processes within a unified representation of AL amyloidosis. To the best of the author’s knowledge, no existing model has integrated plasma cell–derived LC production with subsequent LC misfolding, oligomer formation, fibril deposition, and cardiac injury within a single kinetic framework.

The present study addresses this gap by extending a previously developed model of transthyretin amyloidosis (Kuznetsov 2025a) to AL amyloidosis. The resulting reduced-order framework incorporates clonal plasma-cell dynamics, LC secretion and clearance, LC unfolding, oligomer formation, fibril deposition, and cardiac injury. Two alternative sites of LC aggregation are considered: the blood plasma and cardiac tissue. The model is not intended to reproduce the full molecular and cellular complexity of AL amyloidosis; rather, it is designed to provide a tractable framework for examining how LC production, aggregation kinetics, compartmental volume, intercompartmental transport, and cardiac deposition interact to govern the accumulation of toxic oligomers, fibril burden, and cardiac damage.

The simulations are designed to: (1) quantify the relative contributions of clonal plasma-cell expansion and LC aggregation to the development of cardiac amyloid burden; (2) identify the model parameters that most strongly influence the timing and extent of LC aggregation and fibril deposition within cardiac tissue; and (3) establish a theoretical framework for evaluating the relative contributions of oligomer-induced cardiotoxicity and fibril-associated myocardial stiffening to overall cardiac injury.

## 2. Materials and models

### 2.1. Governing equations

#### 2.1.1. Equations expressing conservation of abnormal clonal plasma cells in the bone marrow and folded LCs secreted by clonal plasma cells

The mathematical model is formulated using balance equations for clonal plasma cells and LC species, including monomers, free oligomers, and fibrils (Fig. 1). A lumped capacitance model is employed, assuming spatial homogeneity of clonal plasma cells within the bone marrow and uniform distribution of all LC species across blood plasma and cardiac tissue, with time as the sole independent variable. In this formulation, model time can be mapped to calendar age by adding the assigned starting age of the modeled interval. The dependent variables are defined in Table 1, and model parameters are summarized in Table 2.

**Table 1.** Summary of the model’s dependent variables.

| Symbol | Definition | Unit |
| --- | --- | --- |
| $[B]_{\text{plasma}}$ | Molar concentration of free LC oligomers in blood plasma | $\mu\text{M}$ |
| $[B]_{\text{tissue}}$ | Molar concentration of free LC oligomers in the cardiac tissue | $\mu\text{M}$ |
| $[F]$ | Plasma molar concentration of natively folded LC monomers secreted by clonal plasma cells | $\mu\text{M}$ |
| $[I]_{\text{plasma}}$ | Molar concentration of LC oligomers deposited into amyloid protofibrils circulating in blood plasma | $\mu\text{M}$ |
| $[I]_{\text{tissue}}$ | Molar concentration of LC oligomers deposited into amyloid fibrils that infiltrated the cardiac tissue | $\mu\text{M}$ |
| $[U]_{\text{plasma}}$ | Plasma molar concentration of unfolded LC monomers susceptible to aggregation | $\mu\text{M}$ |
| $[U]_{\text{tissue}}$ | Cardiac tissue molar concentration of unfolded LC monomers susceptible to aggregation | $\mu\text{M}$ |
| $N$ | Number of clonal plasma cells in bone marrow | |

**Table 2.**
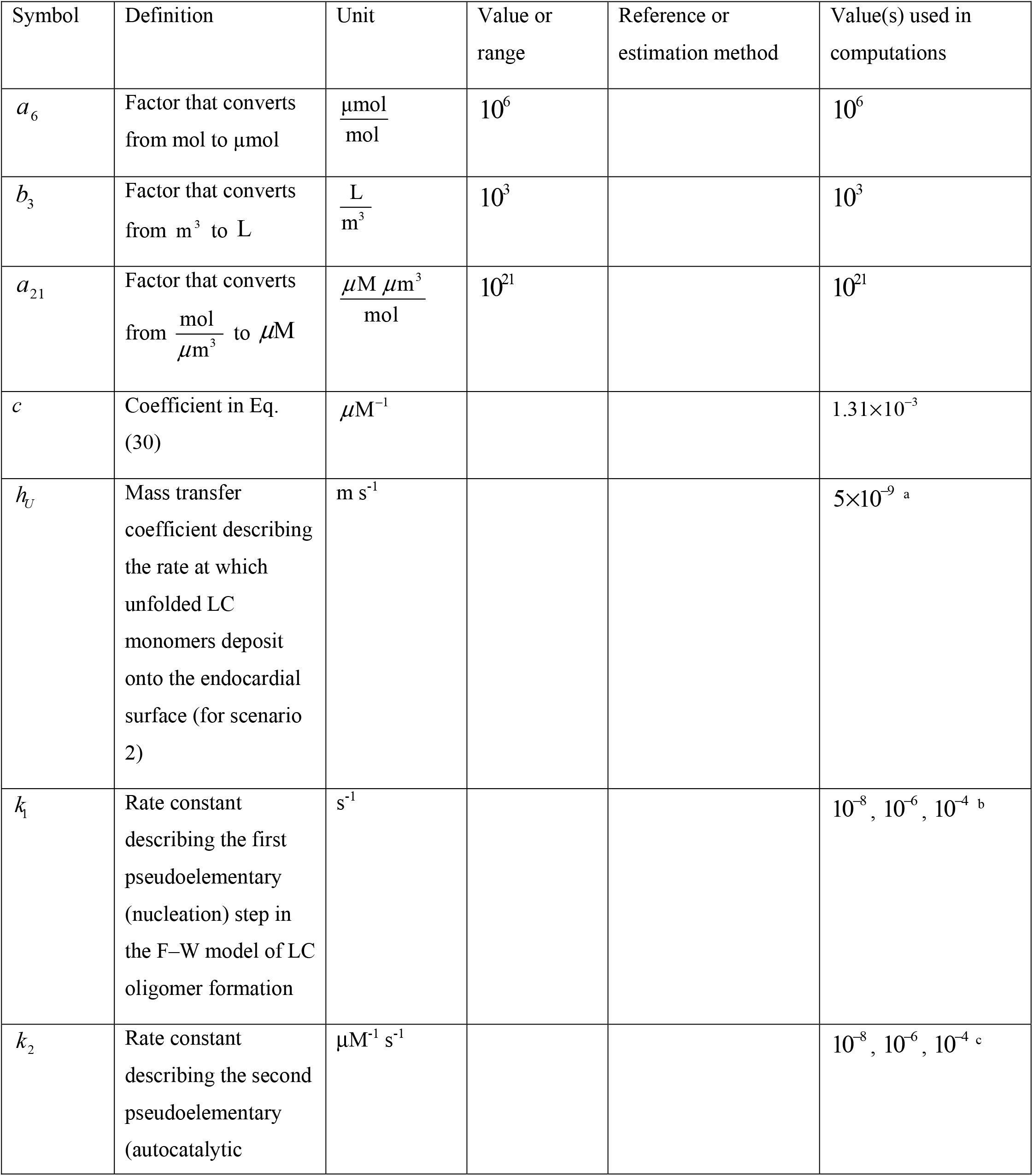
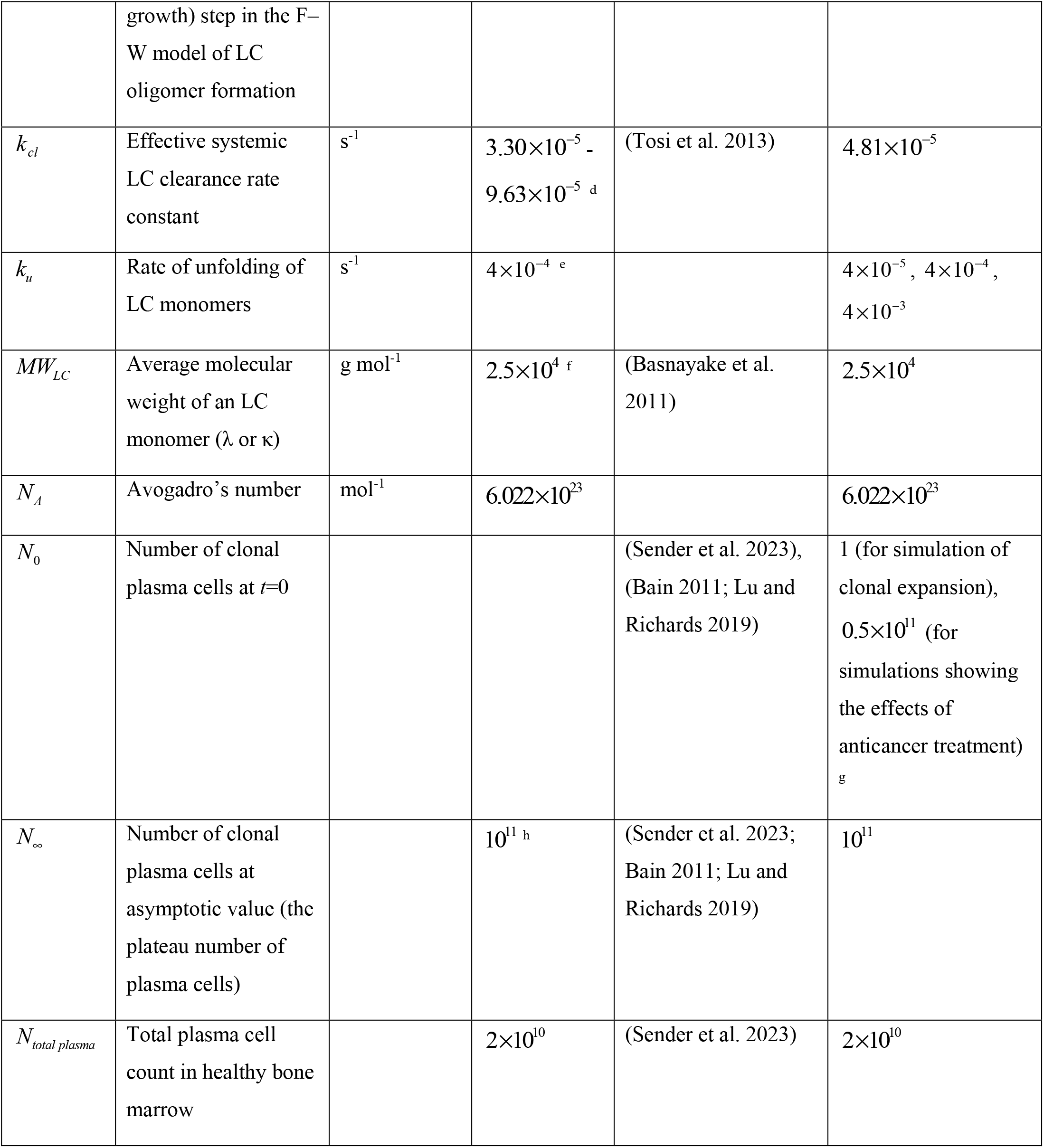
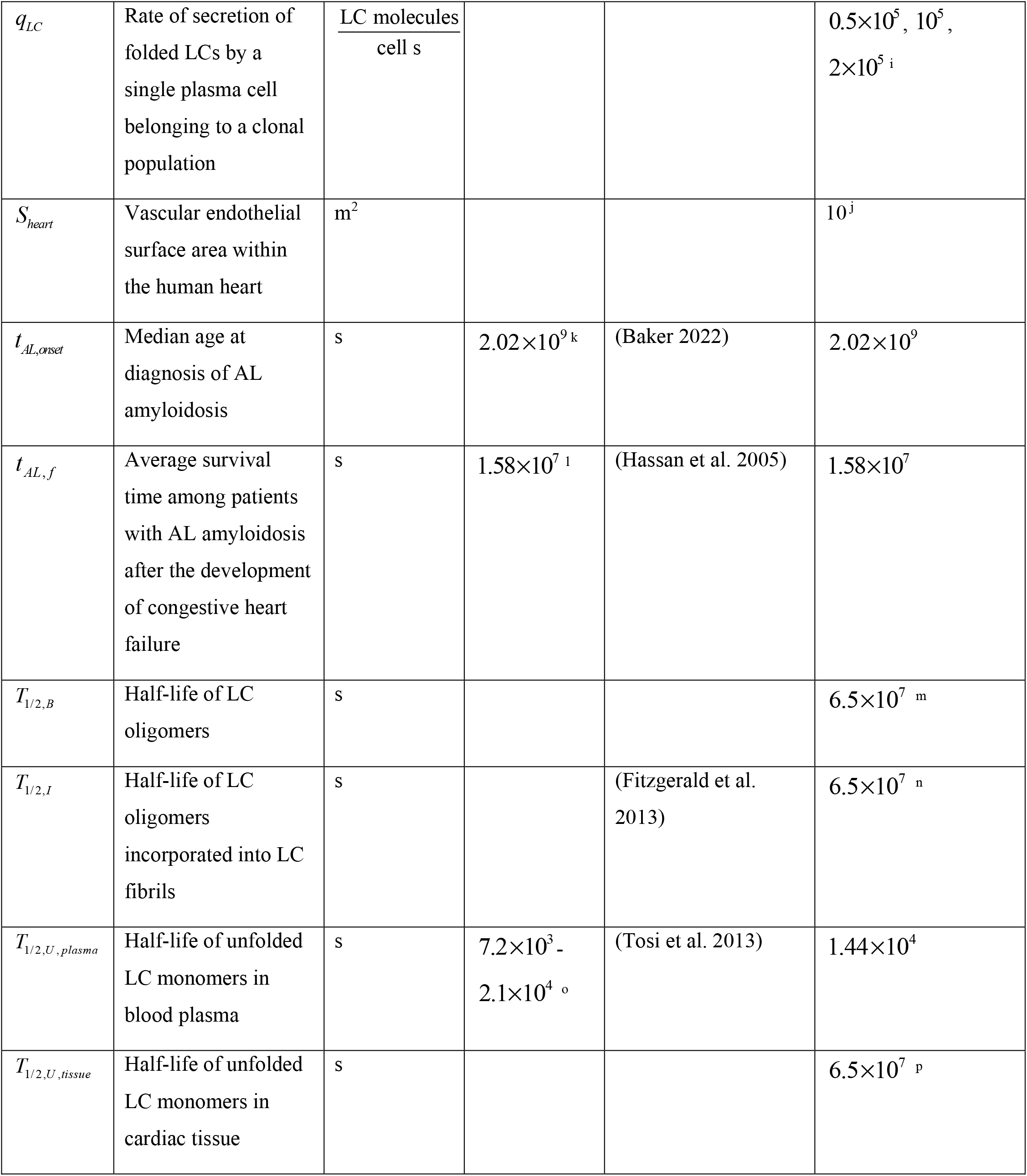
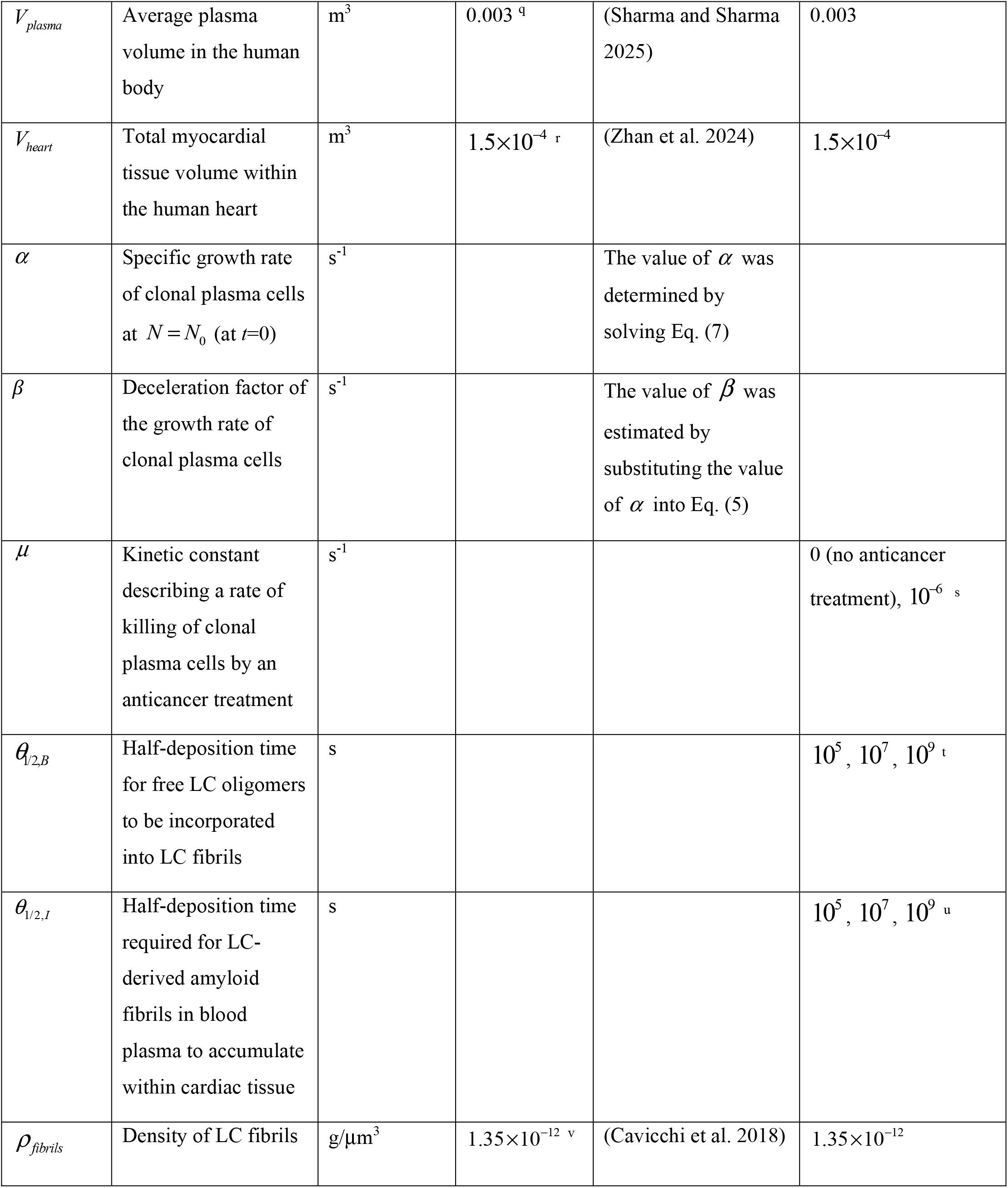
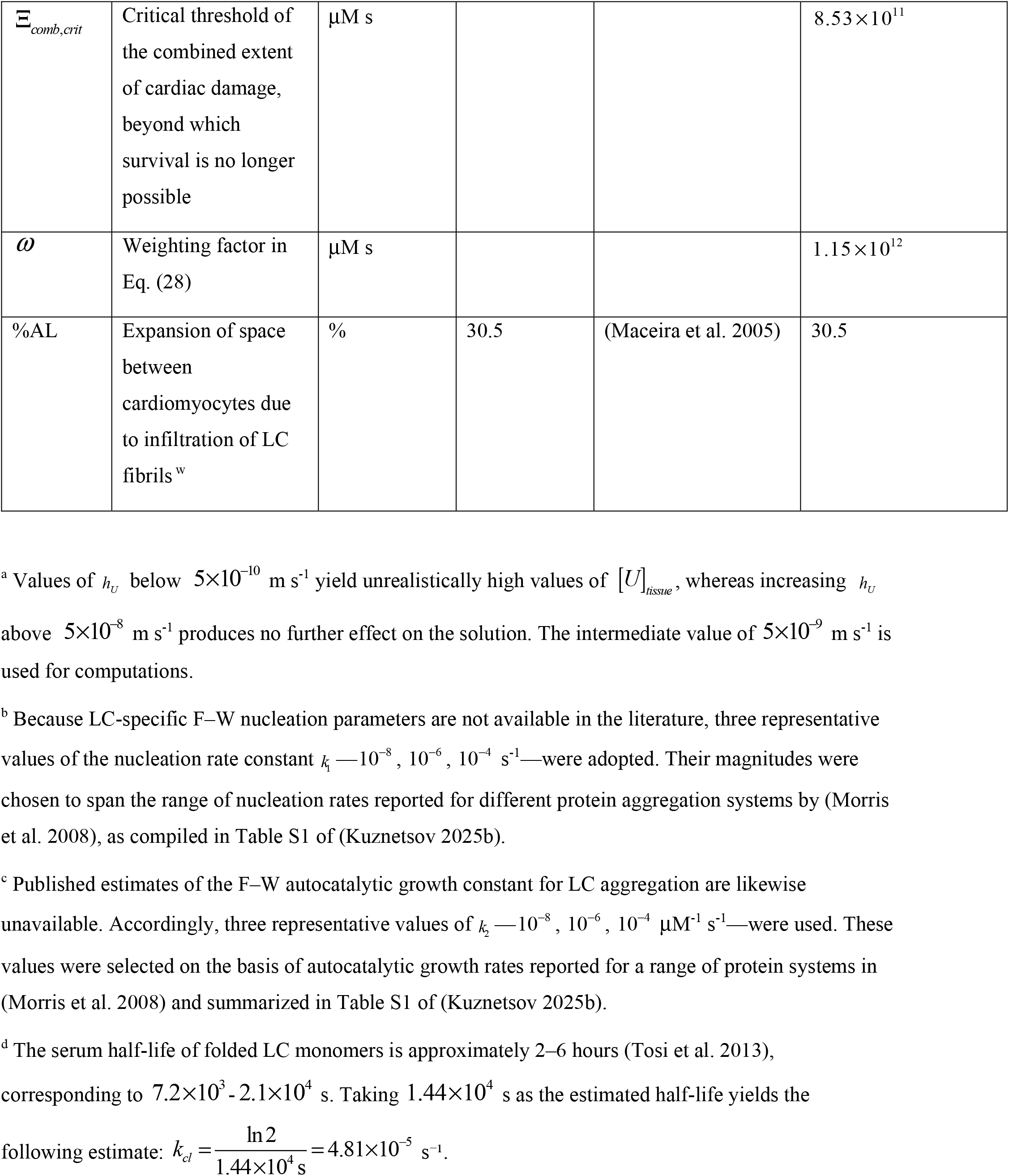
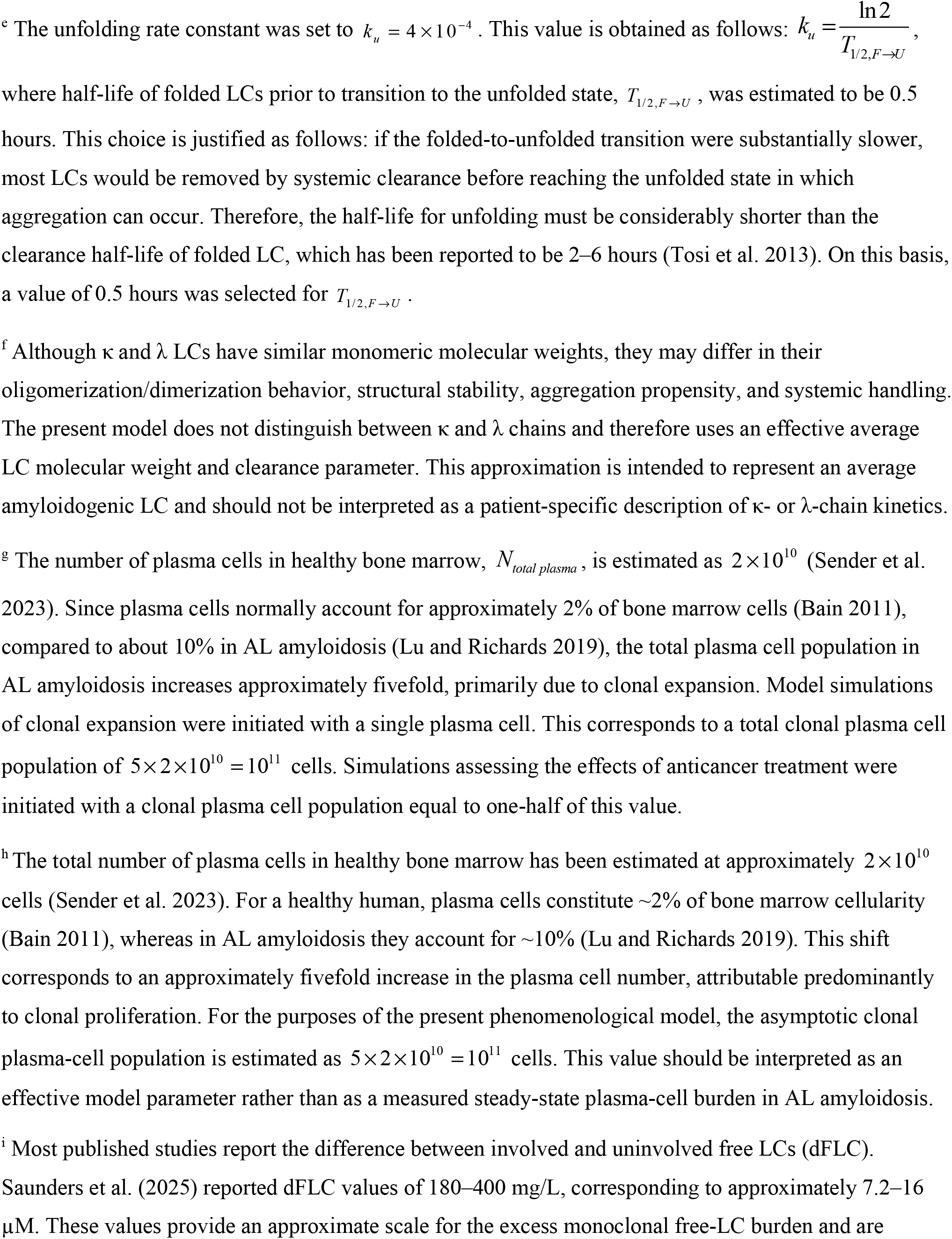
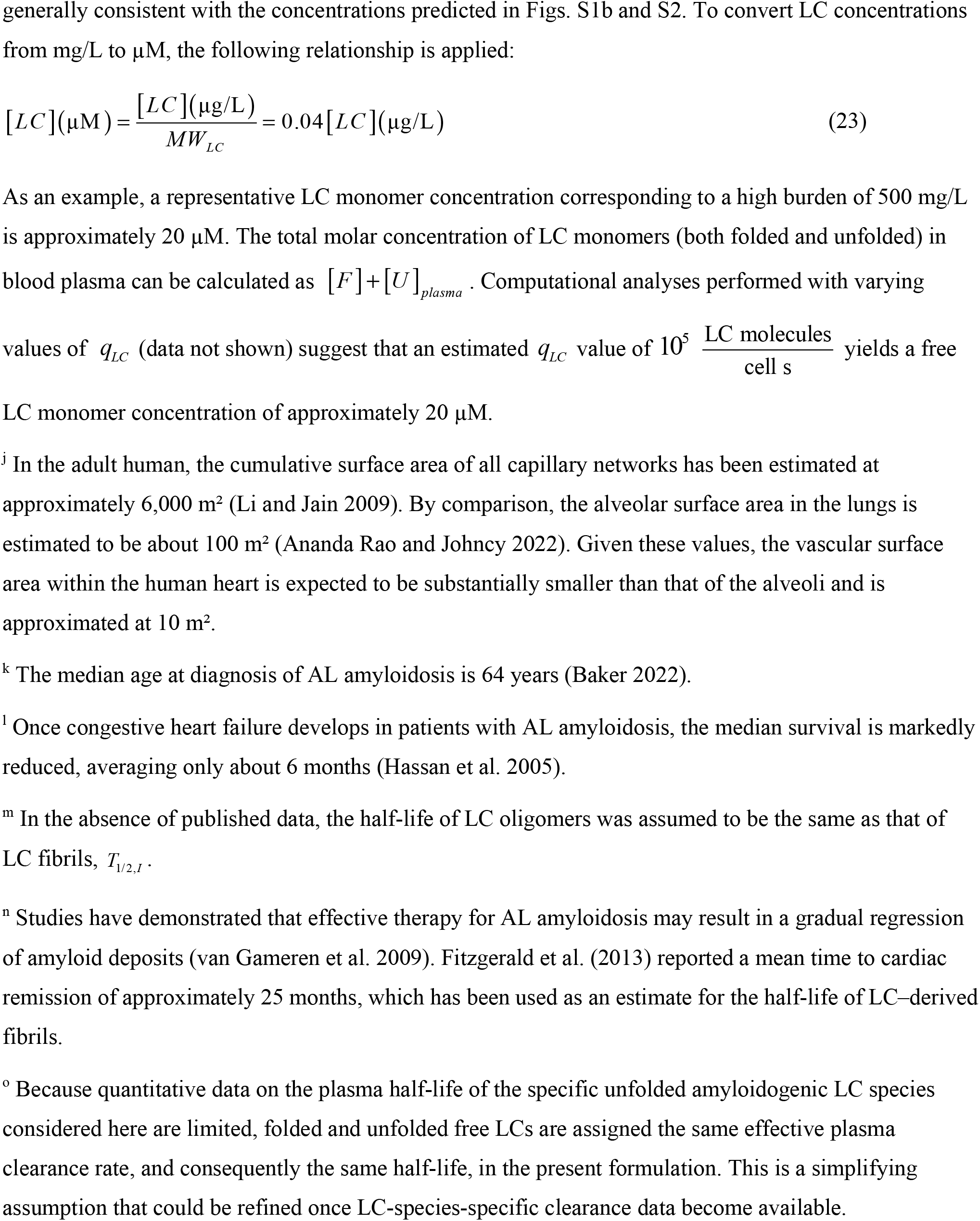
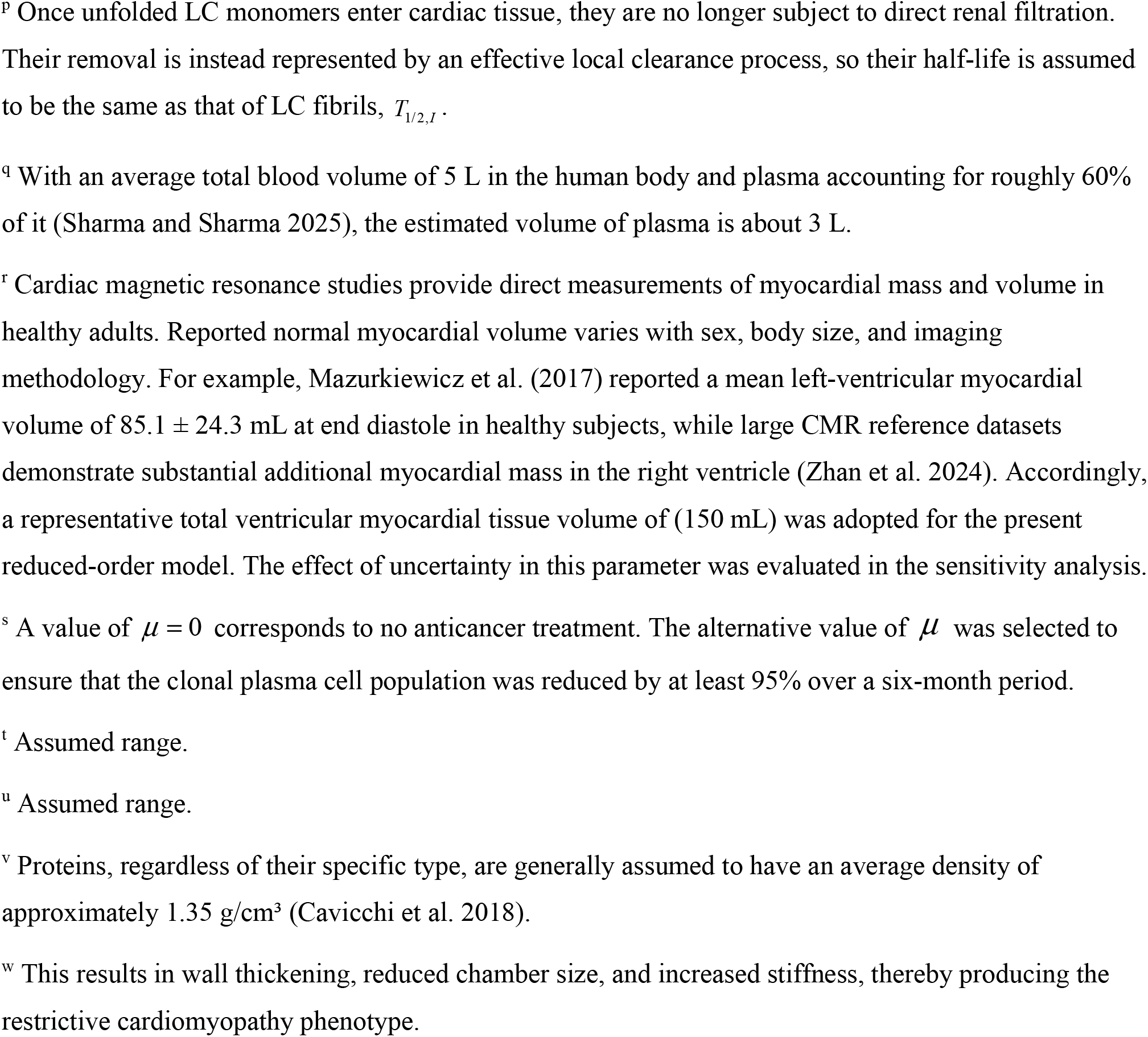
Summary of the model’s parameters and their estimated values.

**Fig. 1.**
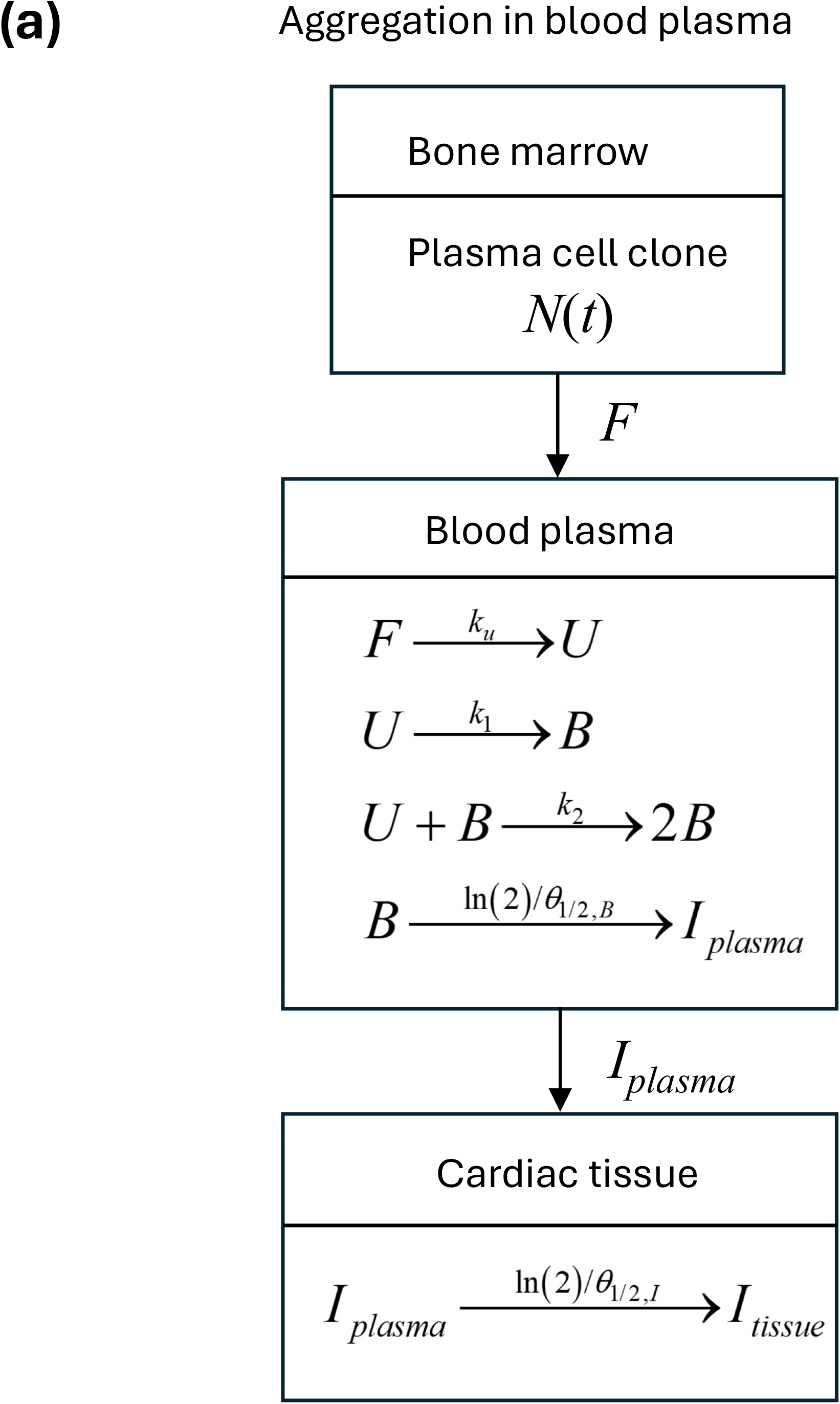
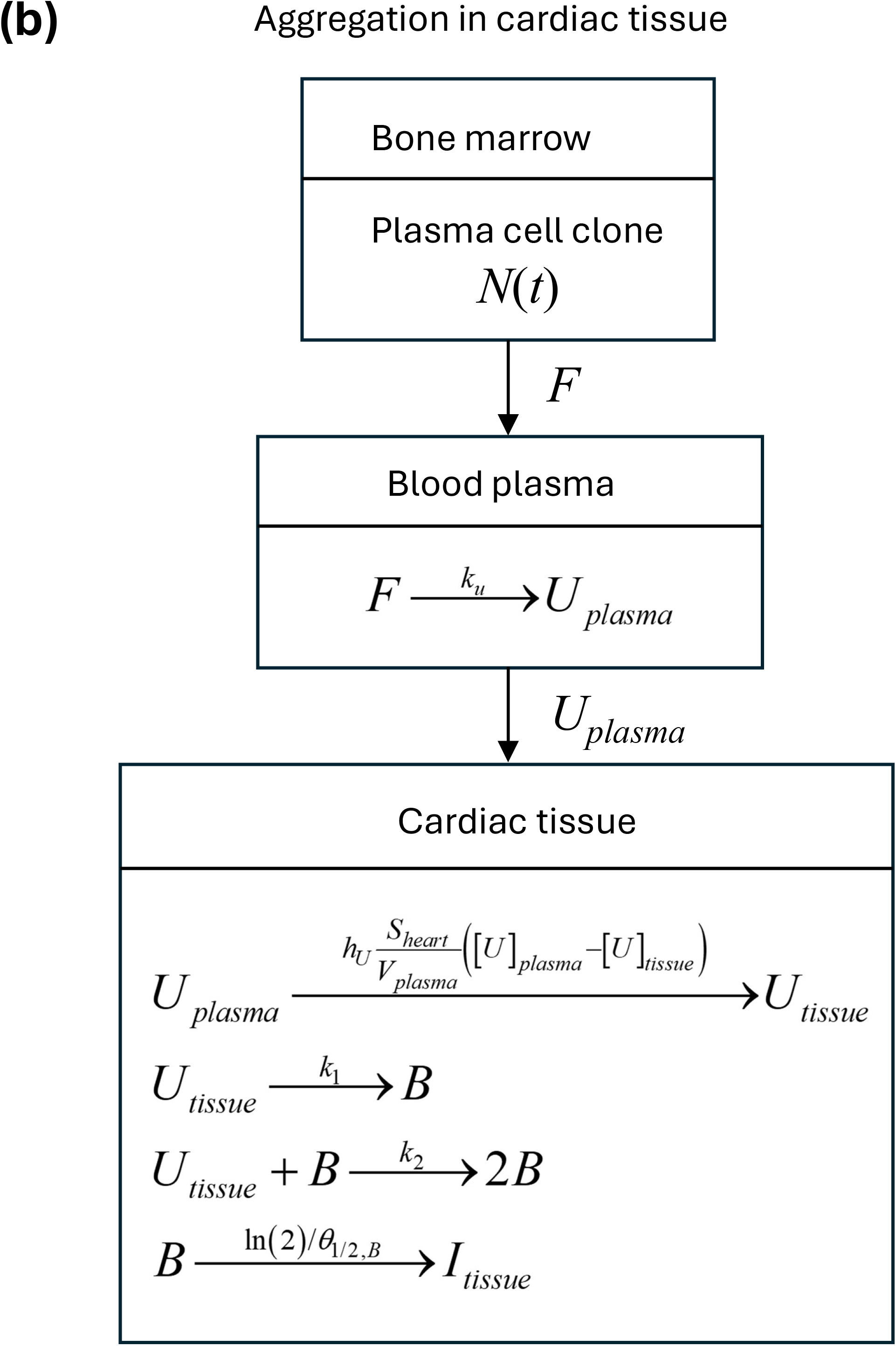
Block diagrams illustrating pathological processes occurring in the bone marrow, blood plasma, and cardiac tissue. (a) Scenario in which aggregation of Ig LCs takes place in the blood plasma. In this case, LC unfolding, oligomer formation through nucleation and autocatalytic mechanisms, and subsequent deposition of oligomers into amyloid protofibrils all occur within the plasma. Circulating fibrils are then deposited in the cardiac tissue. (b) Scenario in which aggregation of Ig LCs occurs within the cardiac tissue. In this case, LCs unfold in the blood plasma, and the unfolded LCs subsequently deposit into the cardiac tissue. Within the tissue, the unfolded LCs form oligomers via nucleation and autocatalytic growth, which then assemble into amyloid fibrils.

AL amyloidosis is a clonal plasma cell dyscrasia that shares biological features with multiple myeloma but represents a distinct clinical entity (Falk et al. 2016). The first equation of the model characterizes the population dynamics of clonal plasma cells, derived from the balance governing the number of these cells. The Gompertz equation is a widely used mathematical model for describing tumor growth, characterized by an initial phase of exponential expansion that gradually decelerates as tumor size approaches a biological maximum or carrying capacity (Vaghi et al. 2020).

In the present study, the Gompertz equation is used as a phenomenological representation of the temporal evolution of the pathogenic plasma-cell clone rather than as a mechanistic model of plasma-cell proliferation. Gompertzian kinetics have previously been used to describe tumor growth in multiple myeloma and are widely employed as phenomenological representations of tumor growth dynamics (Sullivan and Salmon 1972; Ribba et al. 2014). However, AL amyloidosis differs substantially from multiple myeloma and is generally characterized by a relatively small, often indolent plasma-cell clone that produces an amyloidogenic LC (Palladini and Merlini 2016; Palladini et al. 2020). Therefore, the use of the Gompertz formulation in the present model is not intended to imply myeloma-like tumor growth, but rather to provide a smooth time-dependent representation of the source population responsible for monoclonal LC secretion.

Importantly, disease severity in AL amyloidosis is not determined solely by the size of the plasma-cell clone but also by the amount and intrinsic pathogenic properties of the secreted LC, including its conformational stability, misfolding propensity, and capacity to form toxic and amyloidogenic species (Blancas-Mejia et al. 2018; Palladini et al. 2020; Palladini and Merlini 2022). Accordingly, the LC secretion rate per clonal plasma cell is treated in the present model as an independent parameter, allowing the effects of clone size and LC production to be examined separately.

The first equation of the model describes the temporal evolution of the clonal plasma-cell population responsible for LC secretion. Because direct longitudinal measurements of clonal plasma-cell kinetics over the natural history of AL amyloidosis are limited, a phenomenological growth function is adopted. The Gompertz formulation provides a convenient representation of a slowly changing clonal population with an asymptotic plateau, while avoiding the assumption of indefinitely exponential growth.

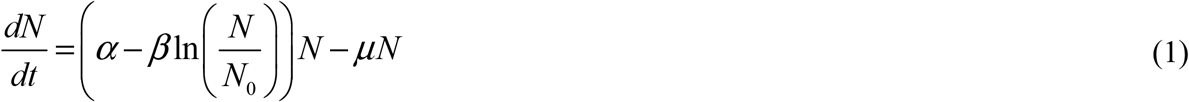

The first term on the right-hand side represents proliferation of the pathogenic plasma-cell clone, whereas the second term represents removal of clonal plasma cells due to anticancer treatment (Murphy et al. 2016; Usher 1994). The Gompertz formulation is used here phenomenologically to describe the time-dependent LC-producing plasma-cell population. The model does not assume that AL amyloidosis exhibits the same clonal kinetics as multiple myeloma. In Eq. (1), *α* denotes the specific growth rate at *N* = *N*_0_, *β* is the deceleration factor of the growth rate, and *μ* is the kinetic constant describing the rate of clonal plasma cell elimination by anticancer treatment.

Eq. (1) is solved subject to the following initial condition:

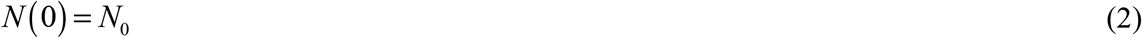

where *N*_0_ denotes the number of clonal plasma cells at *t*=0. In AL amyloidosis, the abnormal plasma cells are clonal, arising from a single progenitor cell (*N*_0_ =1) and therefore genetically identical. Because this study focuses on the late stage of LC aggregation, the use *N*_0_ =1 should be regarded as a of modeling idealization.

The solution to Eqs. (1) and (2) is given by the following function:

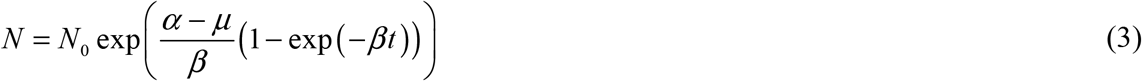

For the case of *μ* = 0, the expression in Eq. (3) reduces to the Gompertzian function, which is defined in Eq. (S1). Further details of the process described by the Gompertzian function are provided in Section S1 of the Supplemental Materials.

If *t* →∞, in the absence of anticancer treatment (*μ* = 0), Eq. (3) reduces to:

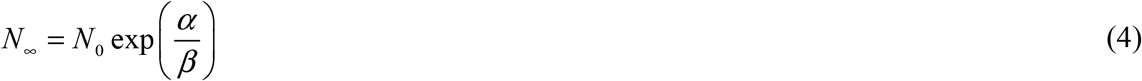

where *N*_∞_ is the plateau number of plasma cells.

From Eq. (4) the following is obtained:

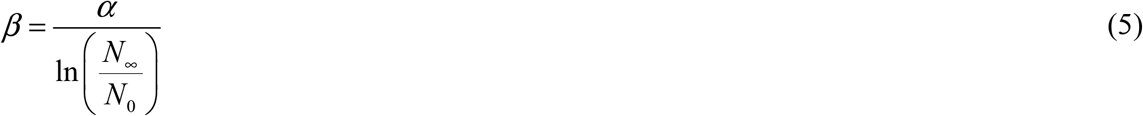

Clinical studies have reported a median survival of approximately six months after the development of congestive heart failure in patients with advanced AL amyloidosis (Hassan et al. 2005). In the present model, this six-month interval is used only to define the characteristic duration of the terminal cardiac-progression phase and is not interpreted as the duration of the underlying plasma-cell disorder or amyloidogenesis. For purposes of parameter estimation, the clonal plasma-cell population is assumed to reach 90% of its asymptotic value (*N*_∞_) at the end of this modeled six-month interval. Accordingly, at *t*_*AL, f*_ = 6 months (1.58×10^7^ s), *N* = 0.9*N*_∞_ . Stating this condition for *μ* = 0, the following equation is obtained:

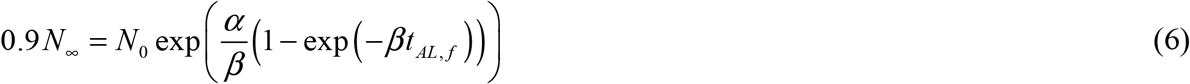

Substituting Eq. (5) into Eq. (6), the following equation for *α* is obtained:

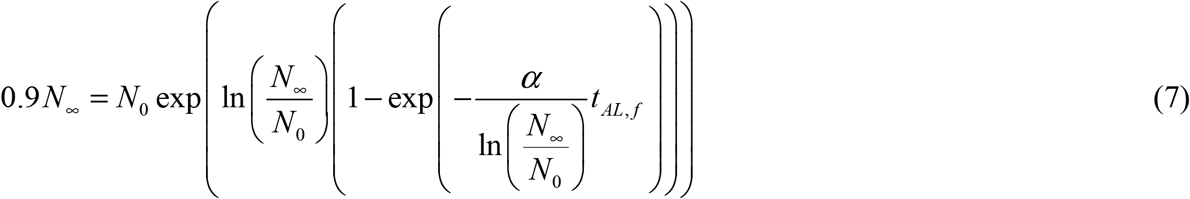

The parameter value for *α* is found by solving Eq. (7), and the parameter value for *β* is estimated by substituting the result in Eq. (5).

Although AL amyloidosis arises from a pathogenic plasma-cell clone, prognosis is determined primarily by the extent of cardiac LC deposition, making reduction of circulating pathogenic LCs a central therapeutic objective (Witteles and Liedtke 2019). Accordingly, the model incorporates conservation equations for LCs at distinct stages of aggregation, including monomeric LCs secreted by clonal plasma cells, soluble oligomers, and insoluble fibrils.

The excess monoclonal LCs fail to assemble into intact Igs and instead exhibit conformational instability, leading to misfolding and aggregation into amyloid fibrils. The progressive deposition of these fibrils in vital organs, particularly the heart, disrupts tissue architecture and function, ultimately leading to organ dysfunction and clinical disease manifestations. In AL amyloidosis, either κ or λ LCs may be overproduced, with λ chains more commonly implicated (Falk et al. 2016). Plasma cells secrete λ LCs primarily as monomers; however, once in circulation, these proteins exhibit a strong propensity to self-associate into noncovalent or disulfide-linked dimers (Goldis et al. 2024). In a simplified model, the dimerization of λ LCs may be considered an integral step in the aggregation process of λ LCs into oligomers. Therefore, at the resolution of the present model, the specific LC type (κ or λ) is not differentiated.

In contrast to intrinsically disordered proteins, Ig LCs are natively folded and require partial unfolding as a prerequisite for aggregation (Blancas-Mejía and Ramirez-Alvarado 2013; Morgan et al. 2019; Kazman et al. 2021). Although LC unfolding is reversible, in AL amyloidosis the equilibrium is likely shifted toward irreversible misfolding and aggregation (Blancas-Mejía, Horn, et al. 2015). Once partially unfolded, LCs preferentially misfold and self-associate rather than refolding into their native conformation. For this reason, the transition of folded LCs to the unfolded state is modeled as an irreversible process:

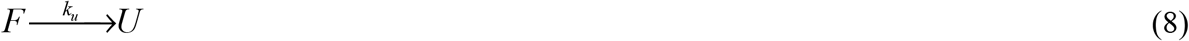

Formulating the conservation of the number of natively folded LC monomers secreted by clonal plasma cells and dividing by *V*_*plasma*_ yields the following equation:

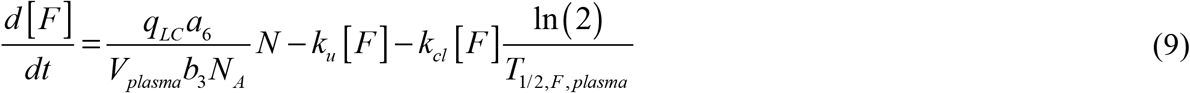

In Eq. (9), the terms on the right-hand side correspond to distinct biological processes: (i) secretion of properly folded LCs by clonal plasma cells; (ii) loss due to conversion into misfolded species prone to aggregation; and (iii) an effective clearance term representing the net removal of circulating properly folded free LCs from plasma; *k*_*cl*_ in this term is the effective systemic LC clearance rate constant. Renal filtration and subsequent proximal tubular catabolism constitute important components of this process, but the model does not resolve the individual physiological clearance pathways (Hutchison et al. 2008; Sanders 2012).

Eq. (9) is solved subject to the following initial condition:

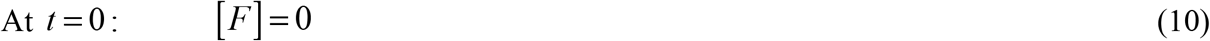

The initial concentration of natively folded LC monomers in Eq. (10) is set to zero, as for sufficiently long simulation times—corresponding to the timescale of cardiac fibril deposition (months) (Hassan et al. 2005)—the influence of the initial condition becomes negligible. Because the present study focuses on the late stage of LC aggregation, setting the initial concentration of [*F*] to zero should be regarded as a modeling idealization rather than a representation of the actual biological state.

The model incorporates assumptions regarding the site of LC aggregation, considering two potential scenarios: (1) aggregation occurs in blood plasma, generating protofibrils that subsequently deposit in organ tissues and become uniformly distributed by diffusion (Fig. 1a); or (2) unfolding of LCs occurs in blood plasma, after which the unfolded LCs deposit in organ tissues, where aggregation subsequently proceeds locally (Fig. 1b). In this study, LC deposition in cardiac tissue was simulated, as cardiac involvement is associated with the poorest prognosis (Banypersad et al. 2012; Tahir et al. 2019).

#### 2.1.2. Development of conservation equations describing amyloid fibril formation for the scenario in which LC aggregation occurs in blood plasma

The minimal model for this scenario tracks unfolded LC monomers susceptible to aggregation, free oligomers in the circulation, oligomers incorporated into amyloid protofibrils within the plasma, and oligomers deposited as amyloid fibrils infiltrating cardiac tissue (Fig. 1a).

In many protein misfolding disorders, aggregation is governed by two principal mechanisms: primary nucleation and secondary nucleation (Rinauro et al. 2024). Secondary nucleation is an autocatalytic process in which pre-existing aggregates catalyze their own formation. This phenomenon has also been demonstrated in LC aggregation (Kazman et al. 2021; Pradhan et al. 2023).

LC oligomerization was modeled using the F–W kinetic framework, a minimal two-step scheme that captures both nucleation and autocatalytic growth (Morris et al. 2008; Iashchishyn et al. 2017; Finke et al. 2020). This model has been extensively utilized to characterize aggregation dynamics in various amyloidogenic proteins (Morris et al. 2008). In this framework, the first pseudoelementary step corresponds to the primary nucleation of LC oligomers, while the second step represents their secondary, autocatalytic conversion into oligomers. The latter process is catalyzed by pre-existing oligomers that act as catalytic templates, enhancing the formation of new oligomers from LC monomers (Aragonès Pedrola et al. 2025).

The first pseudoelementary step corresponds to the primary nucleation of LC oligomers:

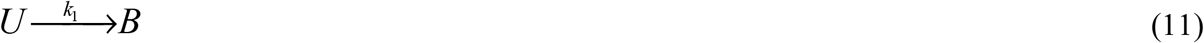

In the F–W model, oligomers are treated as activated monomers capable of autocatalysis. The model does not distinguish between the size or mass of a monomer and an oligomer. The second pseudoelementary step represents the autocatalytic conversion of monomeric LCs into oligomers:

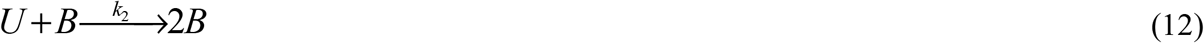

The first F–W step represents the formation of aggregation-competent LC oligomers from unfolded LC monomers and therefore introduces the nucleation timescale. The second step represents autocatalytic conversion of unfolded LC monomers into additional oligomers in the presence of pre-existing oligomers. Thus, the second step captures the positive feedback responsible for the sigmoidal and strongly nonlinear aggregation kinetics predicted by the model.

The F–W representation is intentionally minimal and does not resolve individual oligomer sizes, molecular conformations, fibril lengths, or distinct microscopic secondary-nucleation pathways. Accordingly, the kinetic constants should be interpreted as effective parameters describing the net aggregation behavior represented by the model.

Conservation of unfolded LC monomers in blood plasma gives:

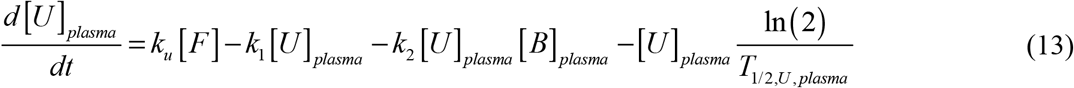

In Eq. (13), the terms on the right-hand side describe the following processes: (i) gain of unfolded LC monomers through conversion from natively folded monomers (corresponding to the second term in Eq. (9), but with opposite sign); (ii) loss of unfolded monomers via nucleation-driven oligomer formation; (iii) loss of unfolded monomers through autocatalytic oligomer production; and (iv) systemic clearance of unfolded monomers, represented by their effective plasma half-life. Renal filtration and proximal tubular catabolism constitute important components of this clearance process (Hutchison et al. 2008; Sanders 2012).

The transformation of free LC oligomers into protofibrils is simulated using an analogy to coagulation phenomena in colloidal suspensions (Boltachev and Ivanov 2020). In this framework, oligomeric LCs (*B*) are assigned a characteristic half-deposition time, *θ*_1/ 2,*B*_, defined as the interval required for half of the oligomer population to incorporate into protofibrils. Conservation of free LC oligomers in plasma, normalized by *V*_*plasma*_, yields:

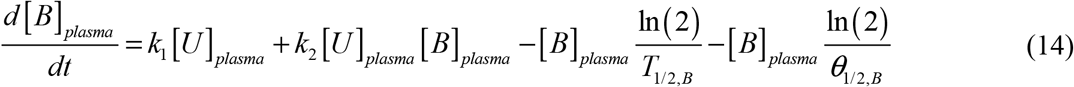

In Eq. (14), the right-hand side terms correspond to the following processes: (i) generation of oligomers through nucleation-driven formation; (ii) additional generation of oligomers via autocatalytic formation (these two terms are equivalent to the second and third terms in Eq. (13), but with opposite signs); (iii) degradation of oligomers due to their finite half-life; and (iv) loss of oligomers through deposition into protofibrils.

Conservation of LC oligomers incorporated into protofibrils circulating in blood plasma, normalized by *V*_*plasma*_, yields the following equation:

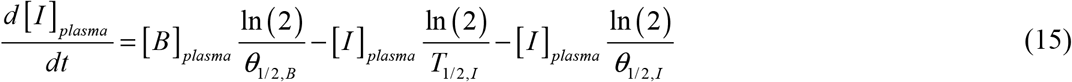

Here, the terms on the right-hand side represent: (i) an increase in the concentration of LC oligomers deposited into protofibrils as free oligomers incorporate into these structures; (ii) a reduction in protofibril concentration in the bloodstream due to their finite half-life; and (iii) a further reduction in concentration resulting from capture of protofibrils in the bloodstream by cardiac tissue.

In this model, protofibrils are assumed to form within the bloodstream, circulate, and deposit in tissues that provide compatible binding sites. The analysis specifically focuses on deposition within cardiac tissue. Conservation of LC oligomers incorporated into fibrils infiltrating the myocardium, after normalization by *V*_*heart*_, yields the following equation:

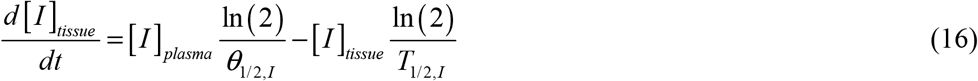

Here, the terms on the right-hand side represent: (i) an increase in cardiac fibril concentration driven by the capture of circulating protofibrils and their subsequent infiltration into myocardial tissue; and (ii) degradation of existing deposits over time (regression), a phenomenon observed when treatment (for example, high-dose chemotherapy combined with stem cell transplantation) induces hematologic remission (Fitzgerald et al. 2013).

The initial conditions for Eqs. (13)–(16) are given by:

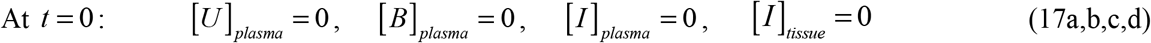

In Eq. (17), the concentrations of all LC species are initialized at zero as a modeling idealization that defines a reference state at the beginning of the simulated advanced cardiac-progression interval. These initial conditions should not be interpreted as implying that pathogenic LCs or cardiac amyloid deposits are absent at the onset of the terminal cardiac phase of AL amyloidosis. Rather, the underlying plasma-cell disorder and amyloidogenic process may have begun substantially earlier than the advanced cardiac stage considered here.

The zero initial conditions are justified by the focus of the present model on the subsequent evolution of LC production, aggregation, and fibril deposition over a timescale of months, corresponding to the terminal stage of cardiac involvement (Hassan et al. 2005). Over this interval, the influence of the assumed initial concentrations becomes progressively less important relative to the LC species generated and accumulated during the simulation. Accordingly, the model is intended to quantify the additional accumulation of LC species and fibrillar deposits during the modeled interval rather than to reconstruct the total amyloid burden accumulated over the full natural history of the disease. Using the same zero initial conditions for both aggregation scenarios also facilitates their direct comparison by isolating the effects of LC production, aggregation, transport, and deposition during the simulated period.

Details of the numerical solution methodology are provided in Section S2 of the Supplemental Materials.

#### 2.1.3 Development of conservation equations describing amyloid fibril formation for the scenario in which LC aggregation occurs in cardiac muscle tissue

In the second scenario, unfolded (and prone to aggregation) LC monomers are taken up by myocardial tissue, where they subsequently aggregate to form amyloid fibrils (Fig. 1b). This scenario is supported by evidence that prefibrillar LC precursors exert direct cardiotoxic effects (Ikura et al. 2022; Rinauro et al. 2024). If aggregation occurred exclusively in the bloodstream with only mature fibrils depositing in cardiac tissue, prefibrillar LCs would not be present within the myocardium.

This model assumes that unfolding of LC monomers occurs in blood plasma. Applying conservation of unfolded monomers in plasma, and normalizing both sides of the equation by *V*_*plasma*_ yields the following equation:

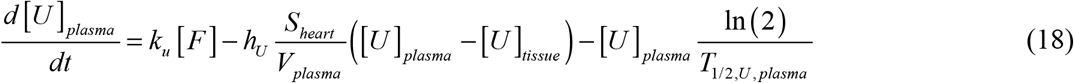

In Eq. (18), the terms on the right-hand side correspond to: (i) generation of unfolded LC monomers via conversion from natively folded species (analogous to the second term on the right-hand side of Eq. (9), but with opposite sign); (ii) deposition of unfolded monomers into cardiac tissue, with *h*_*U*_ *S*_*heart*_ ([*U*]_*plasma*_ − [*U*]_*tissue*_) representing the total deposition rate into the myocardium; and (iii) clearance of unfolded monomers, determined by their finite half-life due to clearance of unfolded monomers through the effective systemic clearance process, of which renal filtration and proximal tubular catabolism are important components (Hutchison et al. 2008; Sanders 2012). Note that as infiltration of LCs into the cardiac tissue progresses, the total LC concentration in the heart—including all aggregation states, such as monomers, oligomers, and fibrils—becomes greater than that in the plasma and is likely to exceed the total plasma LC concentration. However, the concentration of unfolded LC monomers is assumed to remain higher in the plasma than in the cardiac tissue, providing the driving force for their deposition into the heart. The concentration of unfolded monomeric LCs within the cardiac tissue remains low because they rapidly assemble into oligomers. Although receptor-mediated uptake of amyloidogenic LCs has been described in certain cell types, including renal mesangial cells, available evidence suggests that LC internalization by cardiomyocytes occurs predominantly through macropinocytosis rather than through a dedicated LC receptor-mediated pathway (Marin-Argany et al. 2016; Bézard et al. 2024).

Conservation of the number of unfolded LC monomers in cardiac tissue, normalized by *V*_*heart*_, gives:

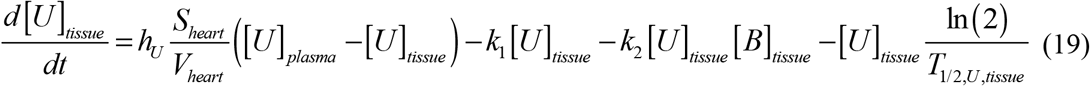

The terms on the right-hand side of Eq. (19) correspond to the following processes: (i) deposition of unfolded monomers into cardiac tissue (analogous to the second term in Eq. (18), but with opposite sign); (ii) and (iii) loss of unfolded monomers through conversion into oligomers via nucleation and autocatalytic mechanisms, respectively; and (iv) clearance of unfolded monomers, determined by their finite half-life within cardiac tissue.

Applying the conservation principle to LC oligomers in cardiac tissue and normalizing the resulting equation by *V*_*heart*_ gives the following:

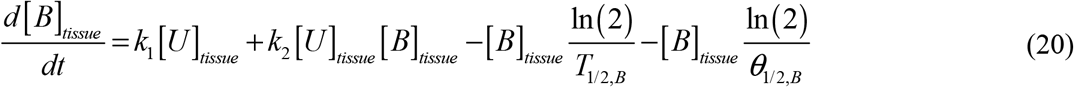

The terms on the right-hand side of Eq. (20) correspond to the following processes: (i) and (ii) represent the generation of free LC oligomers from unfolded monomers through nucleation and autocatalytic pathways, respectively (they are analogues to the second and third terms on the right-hand side of Eq. (19), but with opposite signs); (iii) denotes clearance of oligomers due to their finite half-life; and (iv) accounts for their loss through incorporation into LC fibrils.

By applying the conservation principle to LC oligomers sequestered into fibrillar structures within cardiac tissue, and subsequently normalizing the equation by *V*_*heart*_, the following equation is obtained:

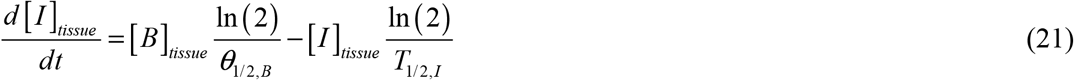

The terms on right-hand side of Eq. (21) simulate the following processes: (i) accumulation of LC fibrils through incorporation of free oligomers into existing fibrillar deposits within cardiac tissue (corresponding to the fourth term in Eq. (20), but with opposite sign); and (ii) degradation of established fibril deposits, a regression that has been observed following successful treatment with high-dose chemotherapy and subsequent stem cell transplantation (Fitzgerald et al. 2013).

Eqs. (18)–(21) are solved subject to the following initial conditions:

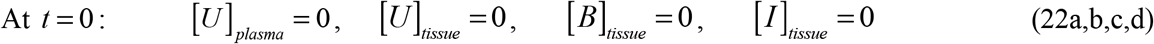

As in Eq. (17), the concentrations of all LC species in Eq. (22) are initialized at zero as a modeling idealization.

### 2.2. Calculation of the volume occupied by LC fibrils in cardiac tissue

Although AL amyloidosis arises from an underlying plasma cell dyscrasia, clinical outcomes are influenced more by the extent of cardiac involvement than by the characteristics of the hematologic malignancy itself (Witteles and Liedtke 2019). Deposition of LC fibrils in the interstitial space between cardiomyocytes increases myocardial stiffness. Accordingly, it is important for the model to predict the extent of cardiac volume expansion resulting from fibrillar infiltration.

This is done by calculating the total number of LC monomers incorporated into fibrils, *N*_*fibrils*_, over a given time *t*. In the F–W model, monomers and oligomers are assumed to have the same molecular weight (see Eq. (11)). Using the approach adapted from Watzky et al. (2008), the following equation is obtained:

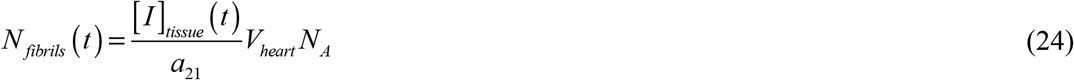

where *N* _*A*_ represents Avogadro’s number.

An alternative approach for determining *N* _*fibrils*_ (*t*) is based on the principle that the number of monomers incorporated into fibrils is proportional to the volume of fibrillar deposits within the myocardium. This relationship can be expressed using the following equation, as also reported in Watzky et al. (2008):

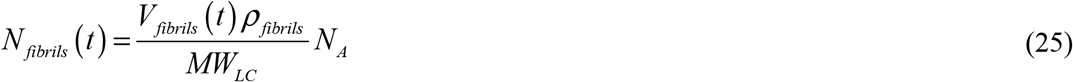

where *MW*_*LC*_ denotes the molecular weight of an LC monomer (λ or κ), *V*_*fibrils*_ (*t*) represents the volume of LC amyloid fibril deposits within the myocardium at time *t*, and *ρ* _*fibrils*_ specifies the density of the fibrillar material.

By equating the right-hand sides of Eqs. (24) and (25) and solving for the fraction of myocardial volume occupied by LC fibrils, the following equation is obtained:

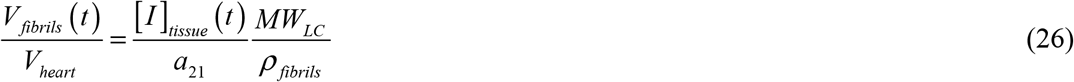

### 2.3. Criteria for accumulated cardiotoxicity of LC oligomers, the combined effect of cytotoxicity and myocardial stiffening, and the proposed biomarker of cardiac cellular biological age

Substantial evidence indicates that soluble oligomers are the main species responsible for cytotoxicity in amyloid diseases. Proposed mechanisms include disruption of membrane integrity through interactions of their hydrophobic surfaces with phospholipids, as well as the induction of apoptotic pathways (Absmeier et al. 2023; Blancas-Mejía and Ramirez-Alvarado 2013; Pradhan et al. 2023).

Analogous to criteria previously proposed to quantify accumulated neurotoxicity induced by Aβ oligomers in Alzheimer’s disease (Kuznetsov 2025c) and by α-synuclein oligomers in Parkinson’s disease (Kuznetsov 2025d), a parameter was defined to quantify accumulated cardiotoxicity attributable to LC oligomers as follows.

For the scenario in which amyloid fibril aggregation takes place in the blood plasma, this parameter is defined as follows:

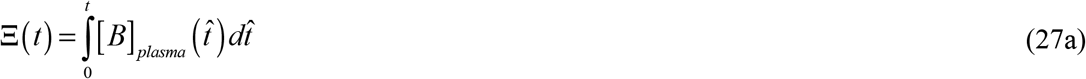

For the scenario in which amyloid fibril aggregation occurs in the cardiac muscle tissue, this parameter is defined as:

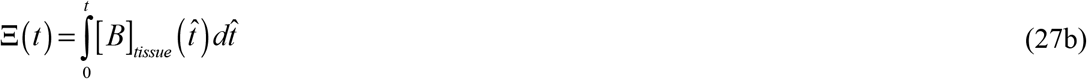

Because cardiac injury in AL amyloidosis arises from both the direct cytotoxicity of LC oligomers and the increased myocardial stiffness caused by fibrillar deposition within cardiac tissue (Falk et al. 2016), the combined extent of cardiac damage—reflecting contributions from both mechanisms—is quantified by the following parameter:

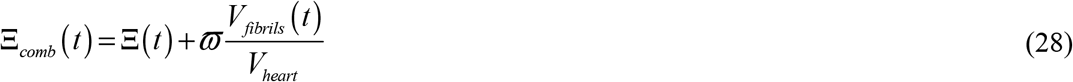

Because LC oligomers circulating in the blood plasma are likely less cytotoxic than those present within cardiac tissue, a scaling factor less than unity should, in principle, be applied to Ξ(*t*) in Eq. (28) when modeling the scenario in which LCs aggregate in the plasma. This adjustment was not implemented here due to insufficient experimental data to determine the appropriate value of this factor.

The value of *ϖ* is estimated as follows. It is assumed that cardiotoxicity and myocardial stiffening each contribute around half of overall cardiac injury in AL amyloidosis. From Fig. S8b for *q*_*LC*_ =10^5^ folded LC molecules per plasma cell per second, the maximum value of Ξ is determined to be 3.50 × 10^11^ µM·s. The proportion of cardiac tissue occupied by LC fibril deposits is estimated at 30.5% (%AL, Table 2). On this basis, the value of *ϖ* is calculated as:

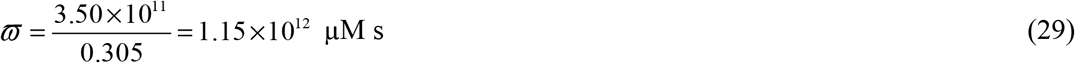

Dormann and Lemke (2024) suggested the development of a new type of aging clock that uses the aggregation dynamics of intrinsically disordered proteins (IDPs)—including amyloid-β (Aβ), tau, α- synuclein, and TDP-43—as a molecular indicator of biological aging. In the present study, this concept is further advanced and applied to Ig LCs, which are natively folded proteins that must undergo partial unfolding as a prerequisite for aggregation. The concept of biological age of the heart (hereafter referred to as biological age) is introduced and defined as follows:

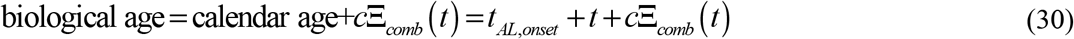

Simulations modeling amyloid fibril formation within cardiac muscle tissue (scenario 2), conducted using the following parameter values: *k* = 10^−6^ s^-1^, *k* = 10^−6^ *μ*M^-1^ s^-1^, *k* = 4×10^−4^ s^-1^, *N* =1 cell, 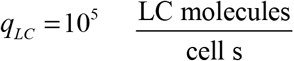, *θ* = 10^7^ s, *μ* = 0 cells^-1^, together with additional parameter values listed in Table 2, yielded a result of Ξ_*comb*_ (*t*_*AL, f*_) = 8.53×10^11^ µM s. When LC aggregation occurs within cardiac tissue, the biological-age criterion reaches 100 years at a calendar age of approximately 64.5 years in the modeled terminal progression scenario. Here, the calendar age of 64 years is used solely as the starting point of the modeled advanced cardiac phase. It does not represent the onset of the underlying plasma-cell clone or the initiation of amyloidogenesis. Thus, the calculated 0.5-year interval should be interpreted as the modeled duration of terminal cardiac deterioration rather than the total duration of AL amyloidosis. This yields the following estimate for the parameter c in Eq. (30):

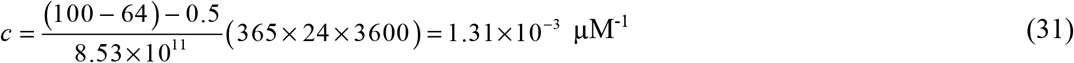

### 2.4. Sensitivity of the criterion characterizing the combined effects of cytotoxicity and myocardial stiffening to various model parameters

The sensitivity of the criterion quantifying combined cardiac damage—encompassing both cardiotoxicity and myocardial stiffening—to various model parameters was investigated. Sensitivity analysis was performed by calculating local sensitivity coefficients, defined as the first-order partial derivatives of the output variable (the combined cardiac damage criterion) with respect to individual parameters, such as the secretion rate of LCs by a single clonal plasma cell, *q*_*LC*_ . The methodology followed established approaches (Beck and Arnold 1977; Zadeh and Montas 2010; Zi 2011; Kuznetsov and Kuznetsov 2019).

For example, the sensitivity coefficient of the combined damage criterion with respect to *q*_*LC*_ was computed using the following equation:

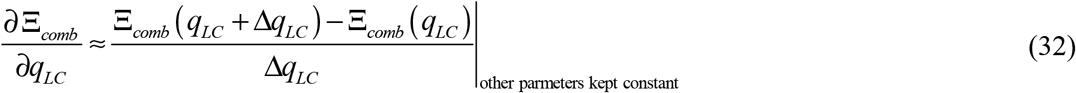

where Δ*q*_*LC*_ =10^−3^ *q*_*LC*_ denotes the step size employed for numerical differentiation. Different step sizes were tested to verify the independence of the sensitivity coefficients from the step size.

To enable direct comparison across parameters expressed in different units and magnitudes, dimensionless relative sensitivity coefficients were also calculated, following the procedures outlined in Zadeh and Montas (2010), Kuznetsov and Kuznetsov (2019). For example:

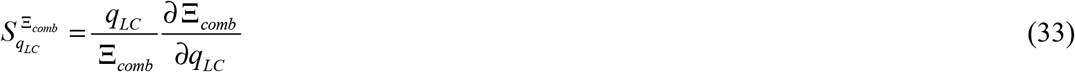

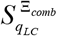 represents the fractional change in Ξ_*comb*_ resulting from a fractional change in *q*_*LC*_, i.e., 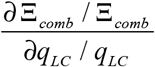.

## 3. Results

The number of clonal plasma cells initially increases exponentially; however, as proliferation slows, the growth curve assumes an S-shaped (sigmoidal) form and eventually reaches an asymptotic value of 10^11^ cells (see footnote *g* after Table 2), consistent with the Gompertz model of cancer cell growth (Fig. S1a). The concentration of folded Ig LC monomers secreted by clonal plasma cells into the blood plasma also exhibits an S-shaped growth profile. The plasma concentration of folded LCs increases with the secretion rate of LCs by an individual clonal plasma cell, and this increase is roughly proportional to the rate of secretion (Fig. S1b).

For the scenario in which LC aggregation occurs in the blood plasma, the molar concentration of unfolded Ig LC monomers initially increases during approximately the first half of the six-month simulation period and subsequently declines (Fig. S2a). The initial increase reflects the growing production of folded LC monomers by clonal plasma cells, followed by their conversion into unfolded, aggregation-prone species. During the latter half of the simulation, the concentration of unfolded LCs decreases as these species are progressively consumed through oligomer formation, particularly via autocatalytic conversion. In contrast, when LC aggregation occurs within cardiac tissue, the plasma concentration of unfolded LCs increases throughout the entire simulation period (Fig. S2b). In both scenarios, the concentration of unfolded LCs increases with increasing secretion rate of folded LCs (Fig. S2).

Saunders et al. (2025) reported dFLC values of 180–400 mg/L, corresponding to approximately 7.2–16 µM (Eq. (23)). These values provide an approximate scale for the excess monoclonal free-LC burden and are generally consistent with the concentrations of [*F*] + [*U*]_*plasma*_ predicted in Figs. S1b and S2, although, for the scenario in which LC aggregation occurs within cardiac tissue, the model may overpredict the concentration of involved LCs.

For the scenario in which LC aggregation occurs within cardiac tissue, the concentration of unfolded LC monomers in cardiac tissue (Fig. S3) is substantially lower than the corresponding concentration of unfolded LCs in the plasma (Fig. S2b). This difference can be attributed to the relatively low value of the mass-transfer coefficient, *h*_*U*_ (5×10^−9^ m·s^−1^, Table 2).

The concentration of free LC oligomers in the blood plasma for the scenario in which aggregation occurs in the plasma (Fig. S4a), and in cardiac tissue for the scenario in which aggregation occurs within the tissue (Fig. S4b), increases slowly during the first three months and much more rapidly during the subsequent three months. The initially slow increase reflects nucleation-driven oligomer formation from unfolded LC monomers, which is a relatively slow kinetic process and precedes the accumulation of a sufficiently large oligomer population to catalyze further oligomer production. During the later stage, oligomer formation accelerates as autocatalytic conversion becomes dominant.

The final oligomer concentration is approximately fivefold higher when aggregation occurs within cardiac tissue (compare Figs. S4b and S4a), even though the volume of the heart is roughly twenty times smaller than the plasma volume (see Table 2). This suggests that oligomer production within the heart is limited primarily by the supply of unfolded LC monomers to the cardiac tissue rather than by the intrinsic aggregation kinetics. This behavior can be attributed to the relatively small value of the mass-transfer coefficient governing the deposition of unfolded LC monomers onto the endocardial surface.

The plasma concentration of LC oligomers incorporated into amyloid protofibrils, [*I*]_*plasma*_, exhibits a pattern similar to that of free oligomers, [*B*]_*plasma*_, but is approximately 25-fold lower (compare Fig. S5 with Fig. S4a). This difference arises from the long half-deposition time of oligomers into fibrils, *θ*_1/2,*B*_ .

The curves displaying the time-dependent concentrations of LC oligomers incorporated into amyloid fibrils infiltrating the cardiac tissue exhibit similar profiles under both scenarios—LC aggregation occurring in the plasma and LC aggregation occurring within the cardiac tissue. However, the concentration is approximately thirty times higher when aggregation occurs in the cardiac tissue (compare Fig. S6b with Fig. S6a). This difference arises because the concentration of LC oligomers is approximately fivefold higher in the scenario where aggregation occurs within the cardiac tissue (compare Fig. S4b with Fig. S4a). An additional sixfold difference results from the fact that, in the plasma aggregation scenario, fibrils must deposit into the cardiac tissue, and the concentration of fibrils in the plasma is about sixfold higher than in the cardiac tissue (compare Fig. S6a with Fig. S5). The latter effect is due to the large half-deposition time of fibrils from the plasma to the cardiac tissue, *θ*_1/2,*I*_ .

The curves displaying the time-dependent fraction of myocardial volume occupied by LC fibrils are similar in shape for both aggregation scenarios—in the plasma and within the cardiac tissue—but are approximately 30 times higher when aggregation occurs in the cardiac tissue (compare Fig. S7a and Fig. S7b). When LCs aggregate within the cardiac tissue, a secretion rate of *q*_*LC*_ =10^5^ folded LC molecules per plasma cell per second results in approximately 44% of the myocardium being occupied by LC fibrils at the end of the six-month modeled terminal cardiac-progression interval (Fig. S7b). At a secretion rate of *q*_*LC*_ =2×10^5^ folded LC molecules per plasma cell per second, this fraction increases to about 89% at the end of the six-month modeled terminal cardiac-progression interval (Fig. S7b), which likely represents an unrealistically high value, as amyloid fibrils have been estimated to occupy approximately 30.5% of the myocardial volume in patients at the terminal stage of the disease (Maceira et al. 2005).

The criterion representing accumulated cardiotoxicity induced by LC oligomers attains values approximately fivefold higher in the scenario where LC aggregation occurs within the cardiac tissue (compare Fig. S8b with Fig. S8a). This difference is attributable to the roughly fivefold higher oligomer concentration observed in this scenario (compare Fig. S4b with Fig. S4a). Notably, most of the accumulated toxicity develops during the latter half of the modeled interval, when it increases rapidly.

The criterion representing the combined effects of LC oligomer-induced cardiotoxicity and myocardial stiffening, Ξ _*comb*_, reaches values approximately tenfold higher in the scenario where LC aggregation occurs within the cardiac tissue (compare Figs. 2b and 2a). In the cardiac tissue aggregation scenario, at an LC secretion rate of *q*_*LC*_ =10^5^ folded LC molecules per plasma cell per second, the combined criterion attains the assumed critical threshold at the end of the six-month modeled terminal cardiac-progression interval. When the secretion rate increases to *q*_*LC*_ =2×10^5^ folded LC molecules per plasma cell per second, this critical threshold is reached after 5 months (Fig. 2b). In contrast, under the scenario of LC aggregation in the blood plasma, even at an LC secretion rate of *q*_*LC*_ =2×10^5^ folded LC molecules per plasma cell per second, the combined criterion remains approximately fivefold below the critical value at the end of the six-month modeled terminal cardiac-progression interval. These findings indicate substantially faster accumulation of modeled cardiac damage when LC aggregation occurs within cardiac tissue than when it occurs in blood plasma.

**Fig. 2.**
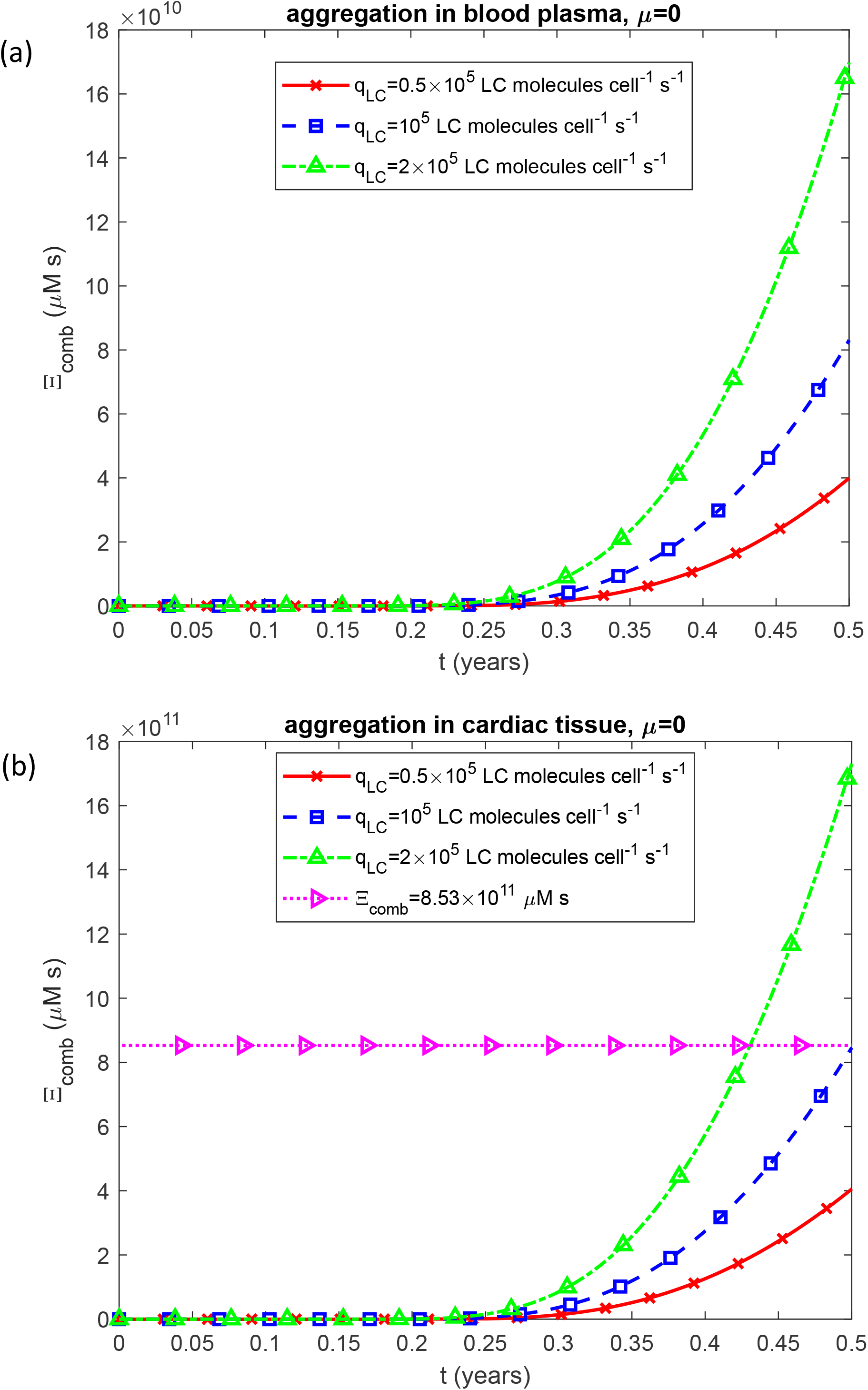
Time-dependent criterion characterizing the combined effect of cytotoxicity and myocardial stiffening, Ξ _*comb*_, vs time, *t*, for two scenarios: (a) aggregation occurring in the blood plasma and (b) aggregation occurring within the cardiac tissue. Results are presented for three different values of the LC secretion rate per clonal plasma cell, *q*_*LC*_ . *k*_1_ =10^−6^ s^-1^, *k*_2_ = 10^−6^ *μ*M^-1^ s^-1^, *k*_*u*_ = 4 ×10^−4^ s^-1^, *θ*_1/ 2,*B*_ = 10^7^ s, and *θ* _1/ 2,*I*_ = 10^7^ s. Results correspond to simulations without anticancer therapy.

Because the combined criterion shown in Fig. 2 is used as a biomarker of aging, Fig. S9, which depicts biological age as a function of chronological age, demonstrates a similar trend. When LC aggregation occurs within cardiac tissue, and the LC secretion rate is *q*_*LC*_ =10^5^ folded LC molecules per plasma cell per second, the biological age reaches 100 years (the assumed human lifespan) at a calendar age of 64.5 years (Fig. S9b). For the purposes of the terminal-stage simulation, the onset of the modeled progression phase is assigned to a calendar age of 64 years. This modeling time point should not be interpreted as the biological onset of AL amyloidosis, which may precede the development of clinically apparent cardiac involvement by a substantially longer period. In contrast, when LC aggregation takes place in the blood plasma, the increase in biological age is much slower. For the same secretion rate *q*_*LC*_, the biological age reaches only 68 years at the end of the six-month modeled terminal cardiac-progression interval (Fig. S9a).

Figs. S10–S18 and Fig. 3 illustrate the effect of the LC monomer unfolding rate constant, *k*_*u*_ . The unfolding rate constant does not affect the number of clonal plasma cells (Fig. S10a). An increase in the LC unfolding rate constant, *k*_*u*_, decreases the concentration of folded LC monomers by accelerating their conversion into unfolded, aggregation-prone species (Fig. S10b). The subsequent behavior of the unfolded LC concentration depends strongly on the assumed site of aggregation. When LC aggregation occurs in the blood plasma, the concentration of unfolded LC monomers initially increases, reaches a maximum, and then decreases (Fig. S11a). During the early stage, production of unfolded monomers through conversion of folded LCs exceeds their removal. As the concentration of LC oligomers increases, however, consumption of unfolded monomers through nucleation and, particularly, autocatalytic oligomer formation becomes progressively stronger. Eventually, these aggregation-related losses exceed the supply of newly unfolded LCs, causing the unfolded-monomer concentration to decline. This behavior is consistent with Eq. (13), in which unfolded LCs in plasma are depleted directly by both nucleation and autocatalytic conversion in addition to systemic clearance.

**Fig. 3.**
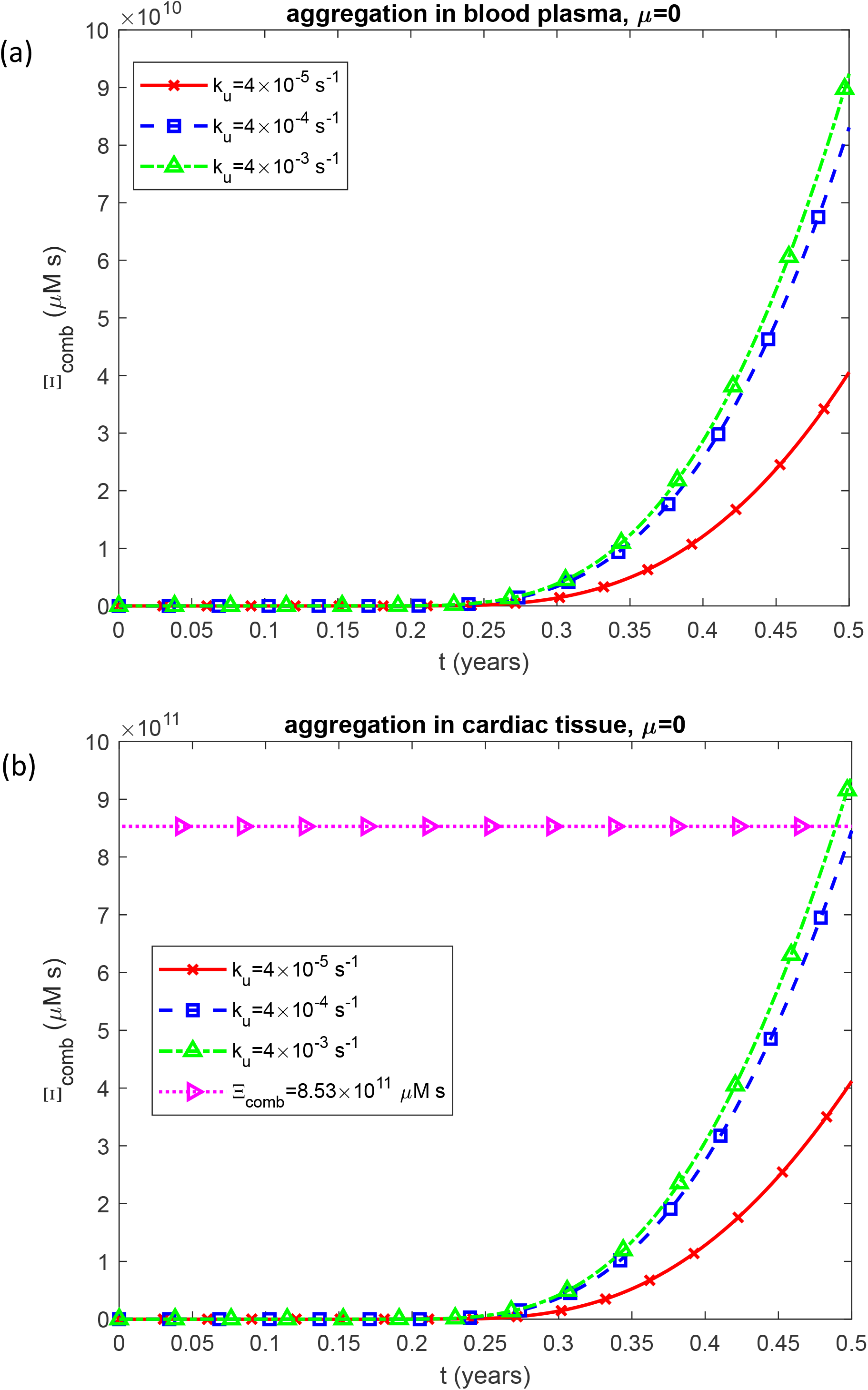
Time-dependent criterion characterizing the combined effect of cytotoxicity and myocardial stiffening, Ξ _*comb*_, vs time, *t*, for two scenarios: (a) aggregation occurring in the blood plasma and (b) aggregation occurring within the cardiac tissue. Results are shown for three different values of the rate of unfolding of LCs, *k*_*u*_ . *k*_1_ =10^−6^ s^-1^, *k*_2_ = 10^−6^ *μ*M^-1^ s^-1^, 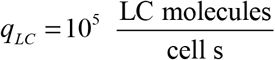, *θ* _1/ 2,*B*_ = 10^7^ s, and *θ*_1/ 2,*I*_ = 10^7^ s. Results correspond to simulations without anticancer therapy.

In contrast, when LC aggregation occurs within cardiac tissue, the concentration of unfolded LC monomers in the plasma increases throughout the entire modeled interval (Fig. S11b). In this scenario, oligomerization does not occur in the plasma; consequently, the plasma pool of unfolded LCs is not directly depleted by nucleation or autocatalytic conversion. Instead, unfolded monomers are removed from the plasma primarily through transport into cardiac tissue and systemic clearance. Because the mass-transfer coefficient governing the deposition of unfolded LC monomers onto the endocardial surface is relatively small (Table 2), their transfer into cardiac tissue is limited. As a result, the continuous production of unfolded LCs from folded monomers exceeds their removal from the plasma over the modeled interval, leading to their progressive accumulation. Increasing the unfolding rate constant, *k*_*u*_, therefore produces a particularly pronounced increase in the plasma concentration of unfolded LC monomers in the cardiac-tissue aggregation scenario (Fig. S11b).

For the scenario in which LC aggregation occurs within cardiac tissue, increasing *k*_*u*_ similarly elevates the concentration of unfolded LC monomers in the cardiac tissue (Fig. S12). The concentration of free (not incorporated into fibrils) LC oligomers also rises with increasing *k*_*u*_(Fig. S13). Likewise, the concentrations of oligomers incorporated into fibrils in the blood plasma (Fig. S14) and in the cardiac tissue (Fig. S15) both increase with increasing *k*_*u*_ . The same trend is observed for the fraction of myocardial volume occupied by LC fibrils (Fig. S16) and for the accumulated cardiotoxicity induced by LC oligomers (Fig. S17).

When LC aggregation occurs within the cardiac tissue, the criterion representing the combined effects of LC oligomer–induced cardiotoxicity and myocardial stiffening, Ξ _*comb*_, reaches approximately half of its critical value (corresponding to patient death) at the end of the six-month modeled terminal cardiac-progression interval for *k*_*u*_ = 4 ×10^−5^ s^−1^ (Fig. 3b). For the same aggregation scenario, Ξ _*comb*_ attains the critical value of 8 ×10^11^ µM·s at the end of the modeled interval for *k*_*u*_ = 4 ×10^−4^ s^−1^ and approximately one week earlier for *k*_*u*_ = 4 ×10^−3^ s^−1^ (Fig. 3b). These results suggest that at lower values, increases in *k*_*u*_ have a strong amplifying effect on Ξ _*comb*_, whereas this effect diminishes as *k*_*u*_ becomes larger. When LC aggregation occurs in the blood plasma, Ξ _*comb*_ remains approximately tenfold below its critical value even for the highest tested *k*_*u*_ value (4 ×10^−3^ s^−1^) (Fig. 3a).

Biological age also depends on *k*_*u*_ . In the cardiac tissue aggregation scenario, the biological age reaches approximately 100 years at the end of the six-month modeled terminal cardiac-progression interval for *k*_*u*_ = 4 ×10^−4^ s^−1^ and about 103 years for *k*_*u*_ = 4 ×10^−3^ s^−1^ (Fig. S18b). In contrast, when aggregation occurs in the blood plasma, the biological age at the end of the six-month modeled terminal cardiac-progression interval is approximately 68.0 and 68.3 years for *k*_*u*_ = 4 ×10^−4^ s^−1^ and 4 ×10^−3^ s^−1^, respectively (Fig. S18a).

An increase in the kinetic constant describing the nucleation rate of LC oligomers, *k*_1_, results in a slight increase of the criterion representing the combined effects of LC oligomer–induced cardiotoxicity and myocardial stiffening, Ξ _*comb*_ (Fig. S19), as well as a slight increase in biological age (Fig. S20). However, the influence of *k*_1_ is considerably smaller than that of *k*_*u*_ .

An increase in the kinetic constant describing the rate of LC oligomer formation by autocatalysis, *k*_2_, leads to a rise in Ξ _*comb*_, particularly at lower values of *k*_2_ (Fig. S21). The biological age shows a similar trend. When LC aggregation occurs in cardiac tissue, the biological age at the end of the six-month modeled terminal cardiac-progression interval is approximately 66 years for *k*_2_ = 10^−8^ µM^−1^s^−1^, 100 years for *k*_2_ = 10^−6^ µM^−1^s^−1^, and approximately 101 years for *k*_2_ = 10^−4^ µM^−1^s^−1^ (Fig. S22b).

The combined criterion characterizing cardiac damage in AL amyloidosis, Ξ _*comb*_, at the end of the six-month modeled terminal cardiac-progression interval, increases with higher rates of LC monomer production per plasma cell, *q* _*LC*_ (Fig. 4). When LC aggregation occurs within cardiac tissue, the curve displaying Ξ_*comb*_ (*q*_*LC*_) crosses the critical threshold of Ξ _*comb*_ corresponding to heart failure at a specific value of *q*_*LC,crit*_ . The critical value of *q*_*LC*_ increases as the LC monomer unfolding rate constant, *k*_*u*_, decreases (Fig. 4b). When LC aggregation occurs in the blood plasma, Ξ _*comb*_ remains below its critical threshold, reaching values approximately fourfold lower than the critical threshold even at the highest *k*_*u*_ value of 4 × 10^−3^ s^−1^ (Fig. 4a).

**Fig. 4.**
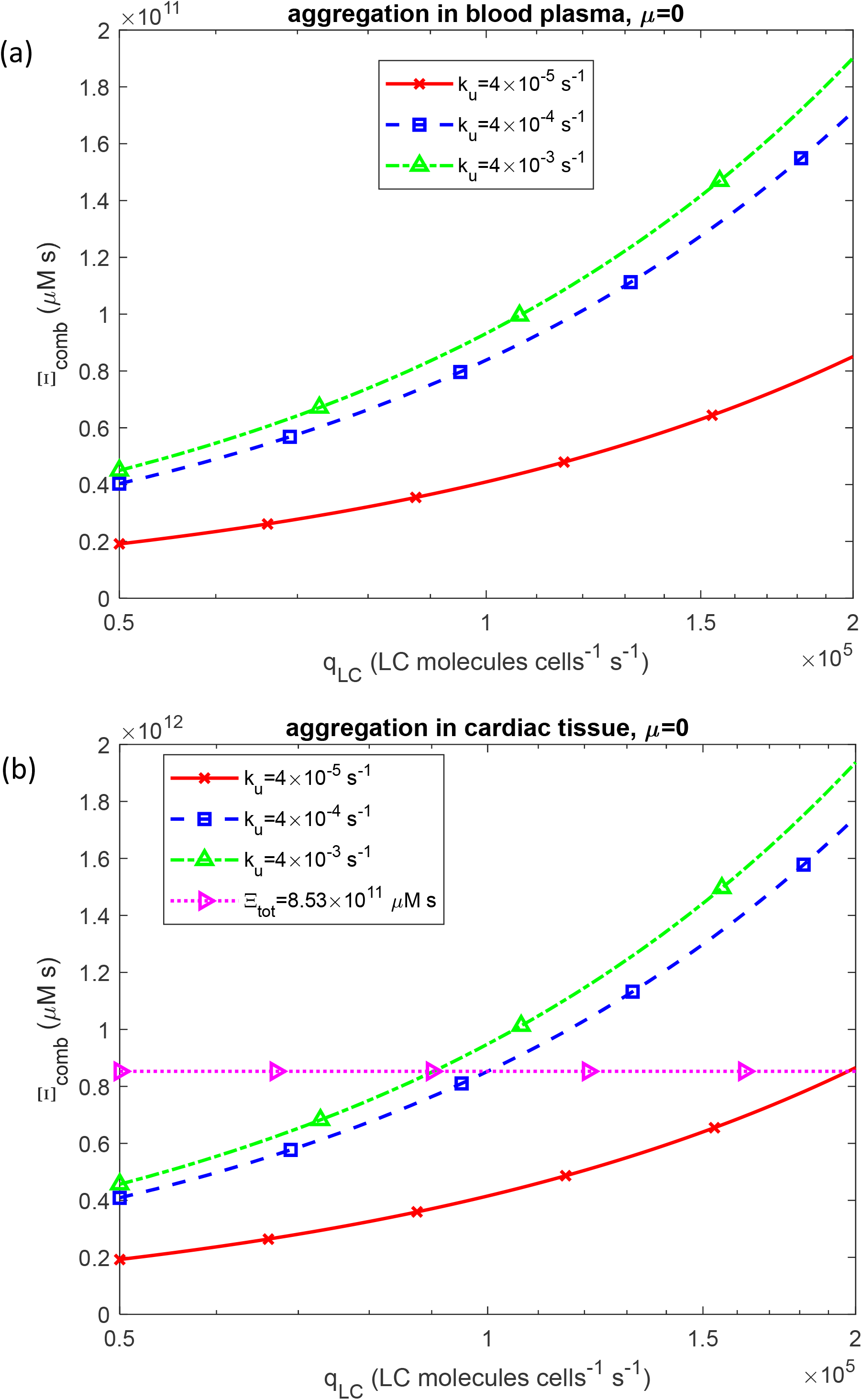
Criterion characterizing the combined effect of cytotoxicity and myocardial stiffening, Ξ _*comb*_, vs the LC secretion rate per clonal plasma cell, *q*_*LC*_, at the end of the six-month modeled terminal cardiac-progression interval for two scenarios: (a) aggregation occurring in the blood plasma and (b) aggregation occurring within the cardiac tissue. Results are shown for three different values of the rate of unfolding of LCs, *k*_*u*_ . *k*_1_ =10^−6^ s^-1^, *k*_2_ = 10^−6^ *μ*M^-1^ s^-1^, *θ*_1/ 2,*B*_ = 10^7^ s, and *θ*_1/ 2,*I*_ = 10^7^ s. Results correspond to simulations without anticancer therapy.

The sensitivity of the criterion characterizing cardiac damage, Ξ _*comb*_, to *q*_*LC*_ is positive (Fig. S23), indicating that Ξ _*comb*_ increases with rising *q*_*LC*_, consistent with the trend shown in Fig. 4. Similar to Ξ _*comb*_, biological age at the end of the six-month modeled terminal cardiac-progression interval increases with the LC monomer secretion rate, *q*_*LC*_ (Fig. S24).

An increase in the unfolding rate constant of LC monomers, *k*_*u*_, also leads to an increase in biological age (Fig. S24). For instance, at the highest secretion rate *q* _*LC*_ (2 ×10^5^ LC molecules per plasma cell per second), in the scenario of LC aggregation within cardiac tissue, the biological age reaches approximately 100 years for *k*_*u*_ = 4 ×10^−5^ s^−1^, 137 years for *k*_*u*_ = 4 ×10^−4^ s^−1^, and 145 years for *k*_*u*_ = 4 ×10^−3^ s^−1^ (Fig. S24b).

In contrast, for the scenario of LC aggregation in blood plasma, the corresponding biological ages are approximately 68, 71.5, and 72.5 years, respectively (Fig. S24a).

For the scenario of LC aggregation in cardiac tissue, an increase in the half-deposition time required for free LC oligomers to be incorporated into fibrils, *θ*_1/2,*B*_, decreases the combined criterion characterizing cardiac damage, Ξ _*comb*_ (Fig. 5b). Interestingly, in the scenario of LC aggregation in the blood plasma, Ξ _*comb*_ is nearly independent of *θ*_1/2,*B*_ (Fig. 5a). A similar pattern is observed in Fig. S25a, which demonstrates that biological age remains unaffected by *θ*_1/2,*B*_ when LC aggregation occurs in the plasma.

**Fig. 5.**
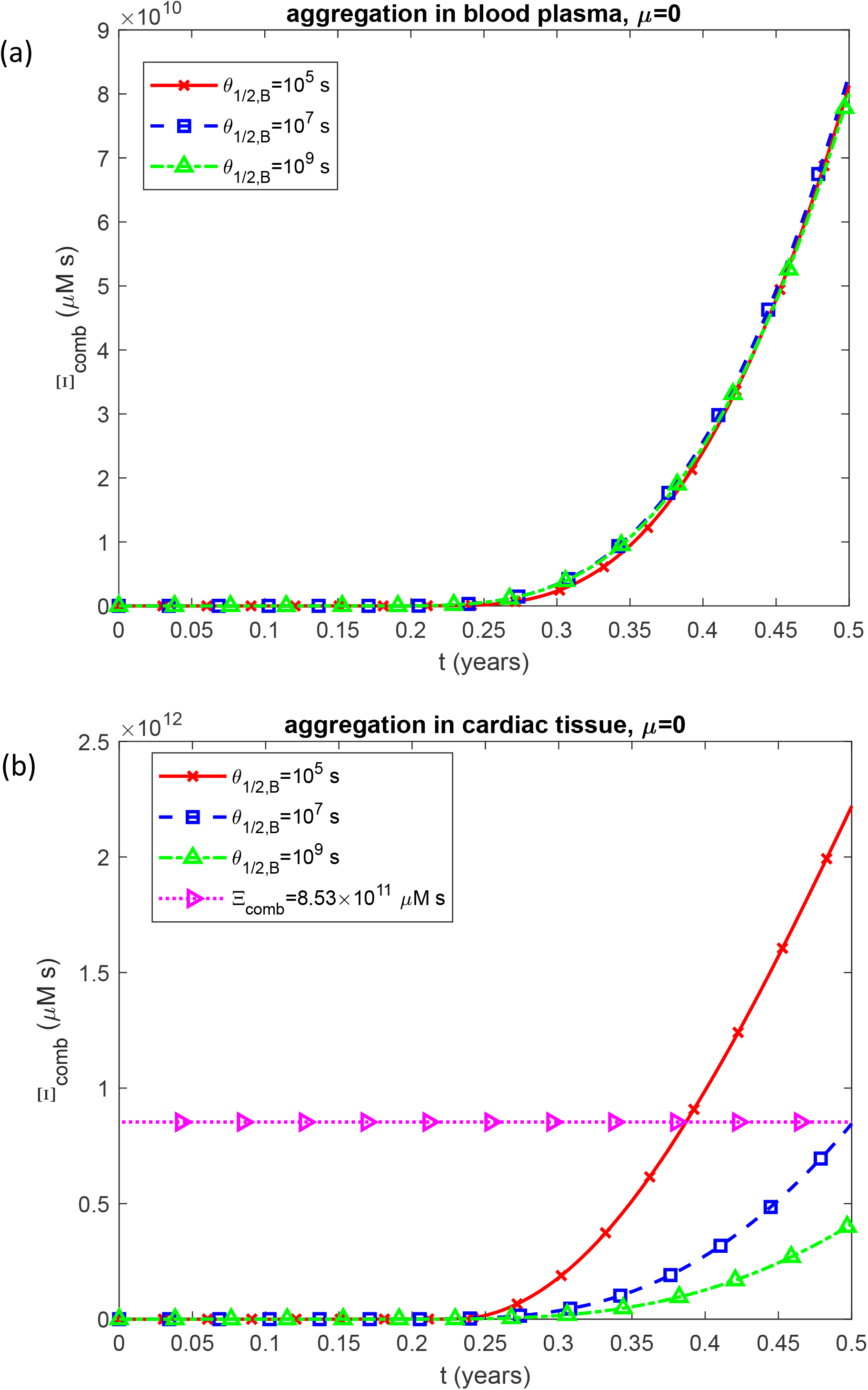
Criterion characterizing the combined effect of cytotoxicity and myocardial stiffening, Ξ _*comb*_, vs time, *t*, for two scenarios: (a) aggregation occurring in the blood plasma and (b) aggregation occurring within the cardiac tissue. Results are shown for three different values of the half-deposition time for free LC oligomers to be incorporated into LC fibrils, *θ*_1/ 2,*B*_ . Computations were performed for *q*_*LC*_ =10^5^ LC molecules per plasma cell per second. *k* =10^−6^ s^-1^, *k* = 10^−6^ *μ*M^-1^ s^-1^, *k* = 4 ×10^−4^ s^-1^, 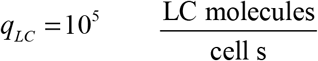, and *θ*_1/ 2,*I*_ = 10^7^ s. Results correspond to simulations without anticancer therapy.

To elucidate the reason for this unexpected phenomenon, the dependences of Ξ _*comb*_ vs *θ*_1/2,*B*_ were plotted in Fig. 6. For a secretion rate of *q*_*LC*_ =10^5^ LC molecules per plasma cell per second, under the scenario in which LC aggregation occurs in the blood plasma (blue dashed curve in Fig. 6a), Ξ _*comb*_ is nearly independent of *θ*_1/2,*B*_ . The blue dashed curve in Fig. 6a is almost horizontal, with only a minor deviation— a small peak in the range of 10^5^ s < *θ*_1/2,*B*_ <10^7^ s —whose magnitude (from minimum to maximum) is less than 10^10^ µM·s. In contrast, when aggregation occurs within the cardiac tissue, the variation along the blue dashed curve in Fig. 6b is approximately 2 ×10^12^ µM·s, about 200-fold greater than that observed in Fig. 6a.

**Fig. 6.**
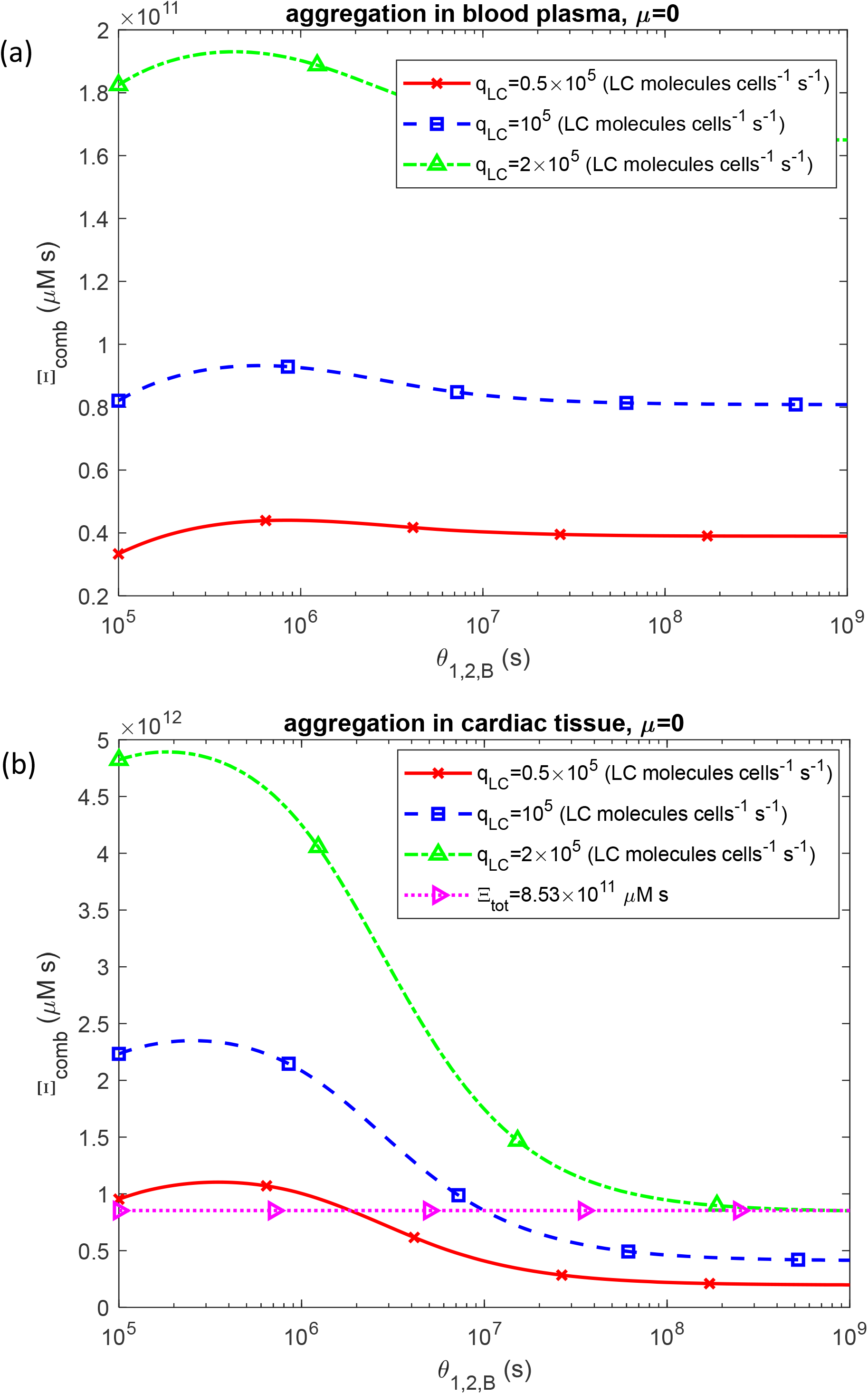
Criterion characterizing the combined effect of cytotoxicity and myocardial stiffening, Ξ _*comb*_, vs the half-deposition time for free LC oligomers to be incorporated into LC fibrils, *θ*_1/2,*B*_, for two scenarios: (a) aggregation occurring in the blood plasma and (b) aggregation occurring within the cardiac tissue. Results are presented at the end of the six-month modeled terminal cardiac-progression interval for three different values of the LC secretion rate per clonal plasma cell, *q*_*LC*_ . *k*_1_ =10^−6^ s^-1^, *k*_2_ = 10^−6^ *μ*M^-1^ s^-1^, *k*_*u*_ = 4 ×10^−4^ s^-1^, and *θ*_1/ 2,*I*_ = 10^7^ s. Results correspond to simulations without anticancer therapy.

A similar pattern is observed for biological age. For a secretion rate of *q*_*LC*_ =10^5^ folded LC molecules per plasma cell per second, in the case of LC aggregation occurring in the blood plasma, the variation in biological age is approximately 0.5 years (blue dashed line in Fig. S26a). For the same value of *q*_*LC*_, when aggregation occurs within the cardiac tissue, the variation in biological age is approximately 100 years (blue dashed line in Fig. S26b), representing the same 200-fold increase as observed in Fig. 6b. To elucidate the near independence of Ξ _*comb*_ from *θ*_1/2,*B*_, the behaviors of the two components comprising Ξ _*comb*_ (as defined in Eq. (28)) were analyzed. The first component, Ξ, represents cardiotoxicity attributable to LC oligomers. The accumulated cardiotoxicity, Ξ, increases with rising *θ*_1/ 2,*B*_ (Fig. 7), because higher values of *θ*_1/ 2,*B*_ extend the residence time of oligomers before their incorporation into fibrils. This leads to an increased oligomer concentration (Fig. 8) and consequently greater cardiotoxicity, as oligomers are considered the primary toxic species.

**Fig. 7.**
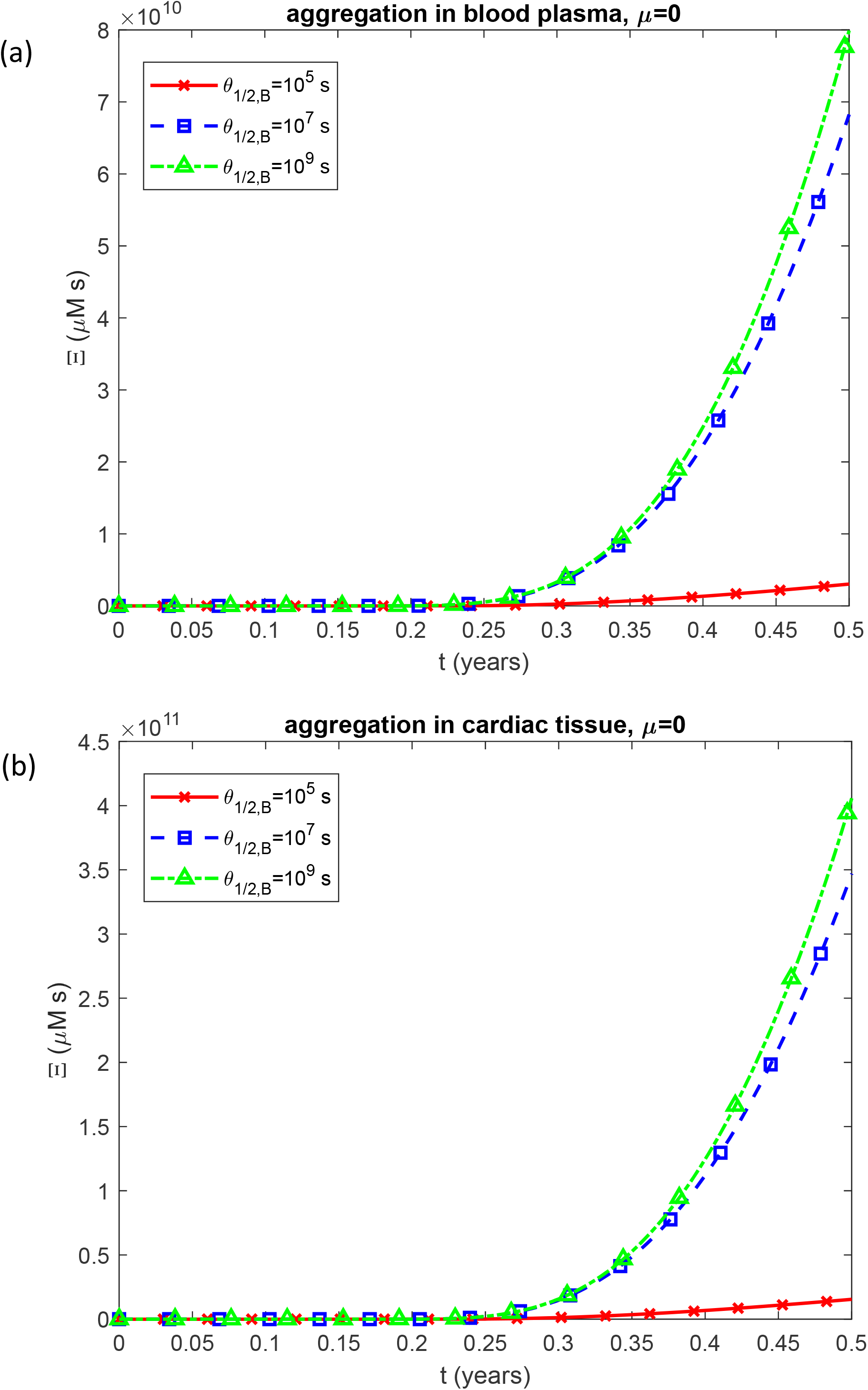
Time-dependent criterion for accumulated cardiotoxicity induced by LC oligomers, Ξ, vs time, *t*, for two scenarios: (a) aggregation occurring in the blood plasma and (b) aggregation occurring within the cardiac tissue. Results are shown for three different values of the half-deposition time for free LC oligomers to be incorporated into LC fibrils, *θ* _1/2,*B*_ . *k*_1_ =10^−6^ s^-1^, *k*_2_ = 10^−6^ *μ*M^-1^ s^-1^, *k*_*u*_ = 4 ×10^−4^ s^-1^, 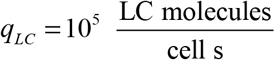, and *θ*_1/ 2,*I*_ = 10^7^ s. Results correspond to simulations without anticancer therapy.

**Fig. 8.**
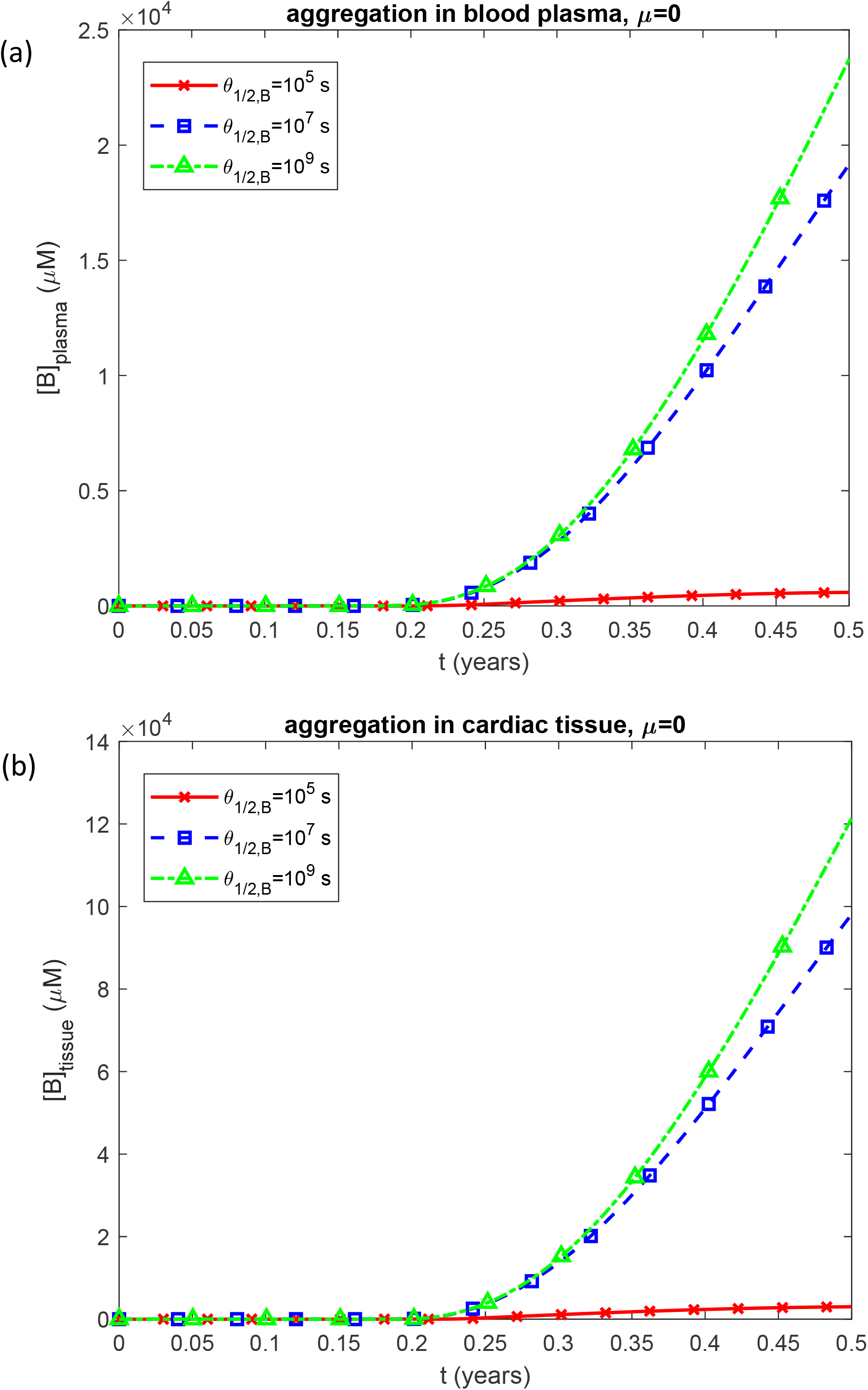
(a) Molar concentration of free LC oligomers in blood plasma, [*B*]_*plasma*_, vs time, *t*, when aggregation occurs in the plasma, and (b) molar concentration of free LC oligomers in cardiac tissue, [*B*]_*tissue*_, vs time when aggregation occurs within the tissue. Results are shown for three different values of the half-deposition time for free LC oligomers to be incorporated into LC fibrils, *θ*_1/2,*B*_ . *k*_1_ =10^−6^ s^-1^, *k*_2_ = 10^−6^ *μ*M^-1^ s^-1^, *k*_*u*_ = 4 ×10^−4^ s^-1^, 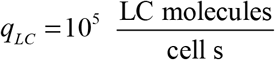, and *θ*_1/ 2,*I*_ = 10^7^ s. Results correspond to simulations without anticancer therapy.

The second component reflects the fraction of myocardial volume occupied by fibrils, *V*_*fibrils*_ / *V*_*heart*_ . As *θ*_1/ 2,*B*_ increases, this fraction decreases (Fig. 9), since a larger *θ*_1/ 2,*B*_ corresponds to more stable oligomers that convert into fibrils more slowly. This trend is consistent with the reduced concentration of LC oligomers deposited into amyloid protofibrils for increased *θ*_1/ 2,*B*_ (Fig. 10).

**Fig. 9.**
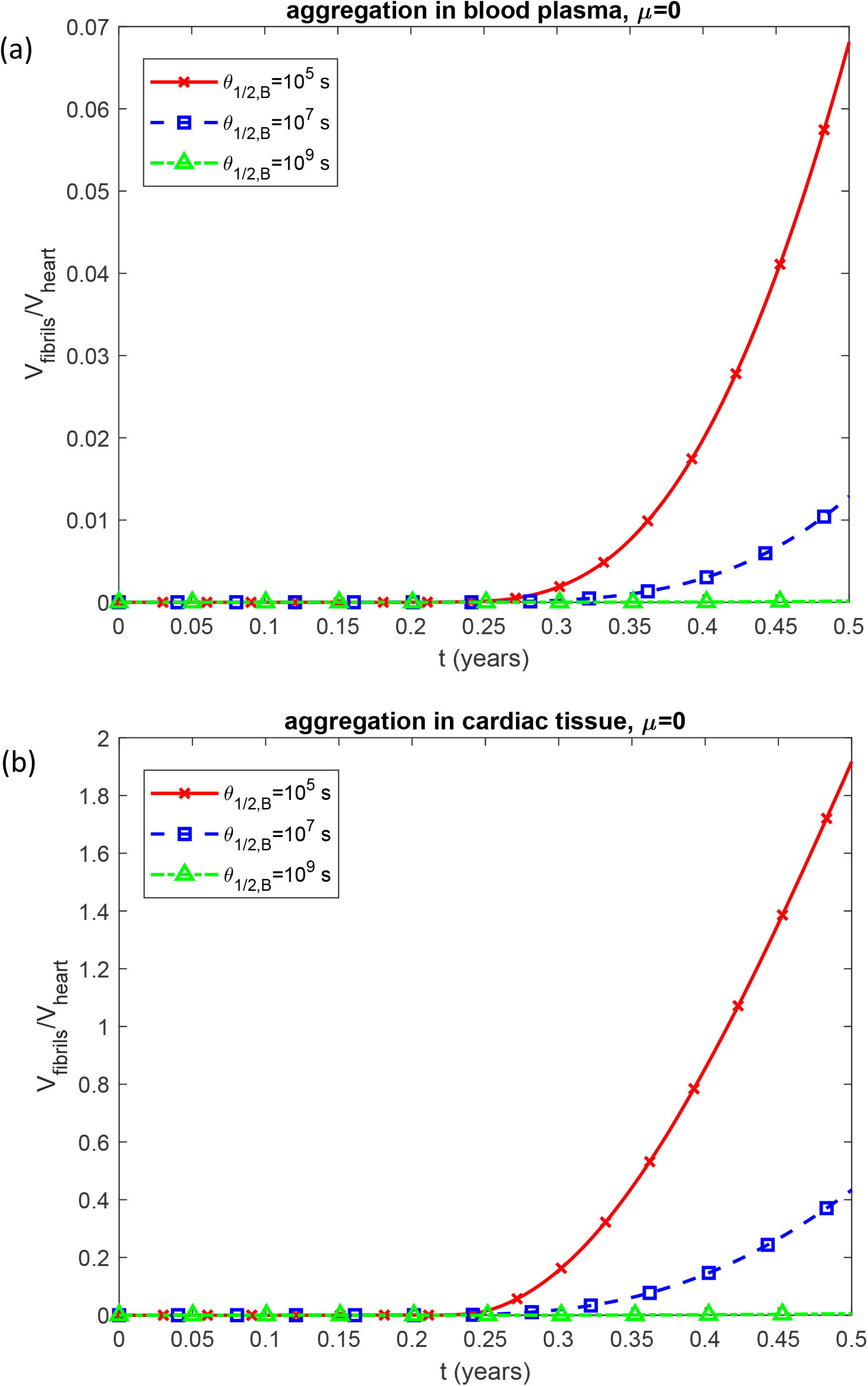
Fraction of myocardial volume occupied by LC fibrils, *V*_*fibrils*_ / *V*_*heart*_, vs time, *t*, for two scenarios: aggregation occurring in the blood plasma and (b) aggregation occurring within the cardiac tissue. Results are shown for three different values of the half-deposition time for free LC oligomers to be incorporated into LC fibrils, *θ*_1/2,*B*_ . *k*_1_ =10^−6^ s^-1^, *k*_2_ = 10^−6^ *μ*M^-1^ s^-1^, *k*_*u*_ = 4 ×10^−4^ s^-1^, 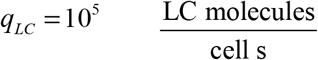, and *θ*_1/ 2,*I*_ = 10^7^ s. Results correspond to simulations without anticancer therapy.

**Fig. 10.**
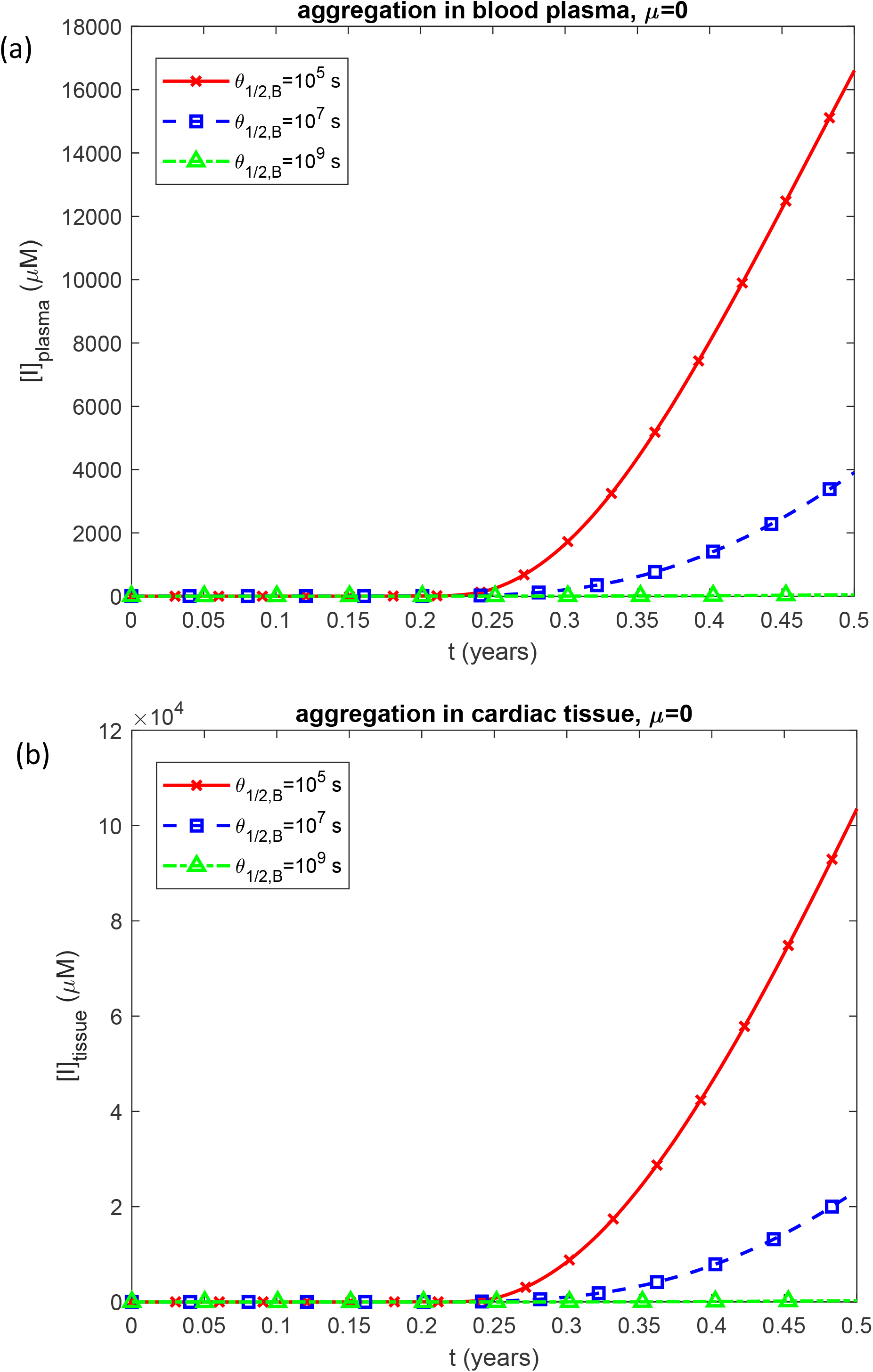
Molar concentration of LC oligomers incorporated into amyloid fibrils infiltrating the cardiac tissue, [*I*]_*tissue*_, vs time, *t*, for two scenarios: (a) aggregation occurring in the blood plasma and (b) aggregation occurring within the cardiac tissue. Results are shown for three different values of the half-deposition time for free LC oligomers to be incorporated into LC fibrils, *θ*_1/2,*B*_ . *k*_1_ =10^−6^ s^-1^, *k*_2_ = 10^−6^ *μ*M^- 1^ s^-1^, *k*_*u*_ = 4 ×10^−4^ s^-1^, 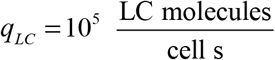, and *θ*_1/ 2,*I*_ = 10^7^ s. Results correspond to simulations without anticancer therapy.

Thus, the combined criterion characterizing cardiac damage, Ξ _*comb*_, comprises two opposing effects: one that increases with *θ*_1/ 2,*B*_ and another that decreases with *θ*_1/ 2,*B*_ . In Figs. 5a and 6a, which represent the scenario of LC aggregation occurring in the blood plasma, these opposing effects nearly cancel each other, resulting in the apparent independence of Ξ _*comb*_ from *θ*_1/ 2,*B*_ .

For the scenario in which LC aggregation occurs in the blood plasma, at a secretion rate of *q*_*LC*_ =10^5^ LC molecules per plasma cell per second, the sensitivity of Ξ _*comb*_ to *θ*_1/ 2,*B*_ is positive within the range of approximately 10^5^ s < *θ*_1/ 2,*B*_ < 10^6^ s (Fig. 11a), corresponding to the slight increase of Ξ _*comb*_ with *θ*_1/ 2,*B*_ observed in Fig. 6a in this range (dashed blue line). The sensitivity becomes negative in the range of approximately 10^6^ s < *θ*_1/ 2,*B*_ < 10^9^ s (Fig. 11a), consistent with the slight decrease of Ξ_*comb*_ with *θ*_1/ 2,*B*_ in this range in Fig. 6a. Within the range of approximately 10^6^ s < *θ* _1/2,*B*_ < 10^9^ s, the sensitivity approaches zero (Fig. 11a), aligning with the nearly horizontal, *θ*_1/2,*B*_ -independent behavior of Ξ _*comb*_ shown in Fig. 6a in this range.

**Fig. 11.**
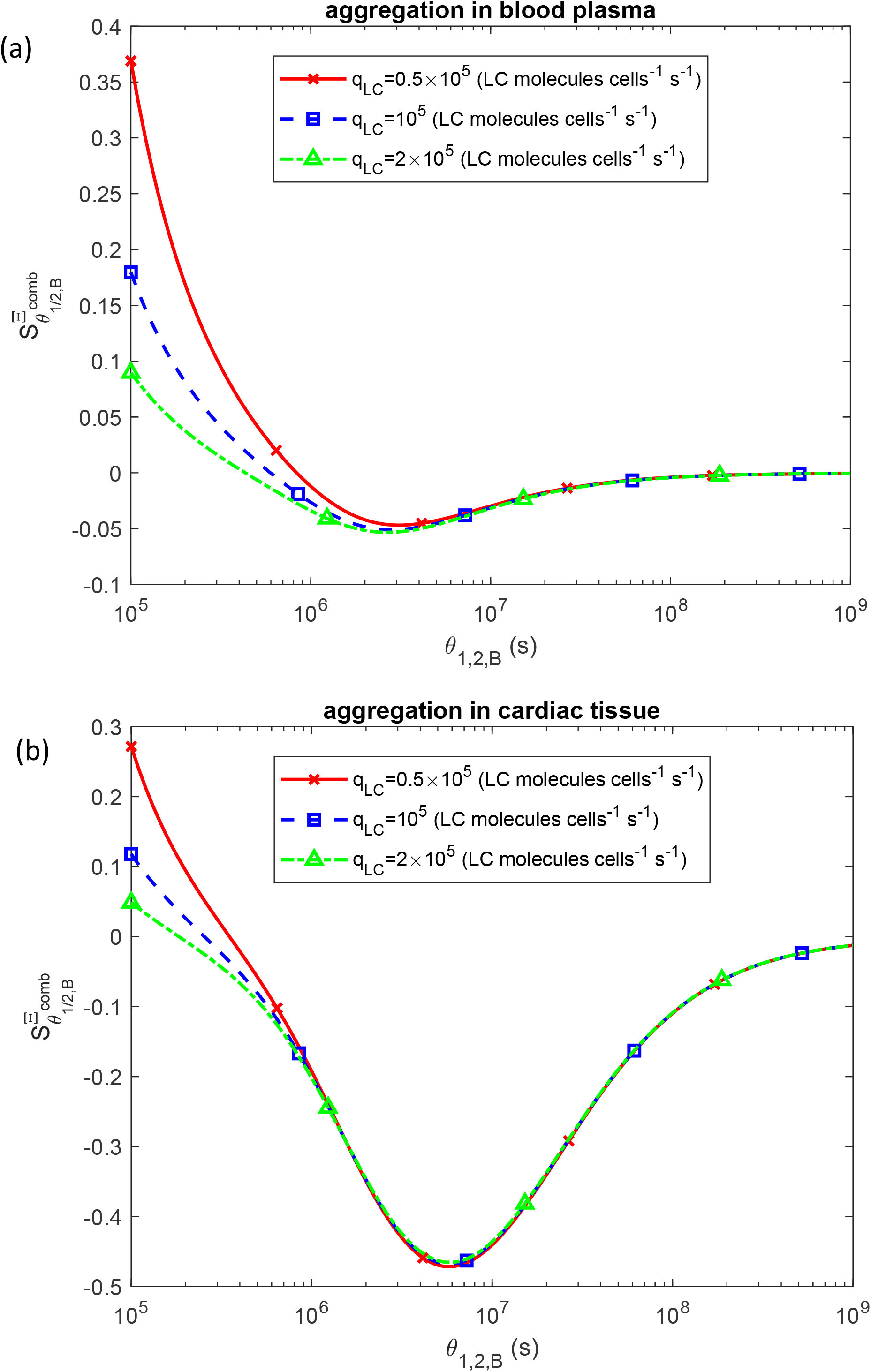
Dimensionless sensitivity of the criterion characterizing the combined effects of cytotoxicity and myocardial stiffening to the half-deposition time for free LC oligomers to be incorporated into LC fibrils, 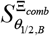, vs *θ*_1/2,*B*_ . Results are presented for three different values of the LC secretion rate per clonal plasma cell, *q*_*LC*_ . *k*_1_ =10^−6^ s^-1^, *k*_2_ = 10^−6^ *μ*M^-1^ s^-1^, *k*_*u*_ = 4 ×10^−4^ s^-1^, and *θ*_1/ 2,*I*_ = 10^7^ s. Results correspond to simulations without anticancer therapy.

For the scenario in which LC aggregation occurs within the cardiac tissue, the sensitivity is negative throughout the range of approximately 2 ×10^5^ s < *θ* _1/2,*B*_ < 10^9^ s (Fig. 11b), consistent with the decrease of Ξ _*comb*_ ^with^ *θ*_1/ 2,*B*_ observed in Fig. 6b in this range.

The case involving anticancer treatment is analyzed in Figs. S27–S35 and Fig. 12. The initial number of clonal plasma cells was assumed to be 5 ×10^10^ cells. During therapy, the clone population progressively declined and was almost completely eliminated after six months of treatment (Fig. S27a). Because folded LC monomers are secreted by clonal plasma cells, their concentration likewise decreased over time (Fig. S27b).

**Fig. 12.**
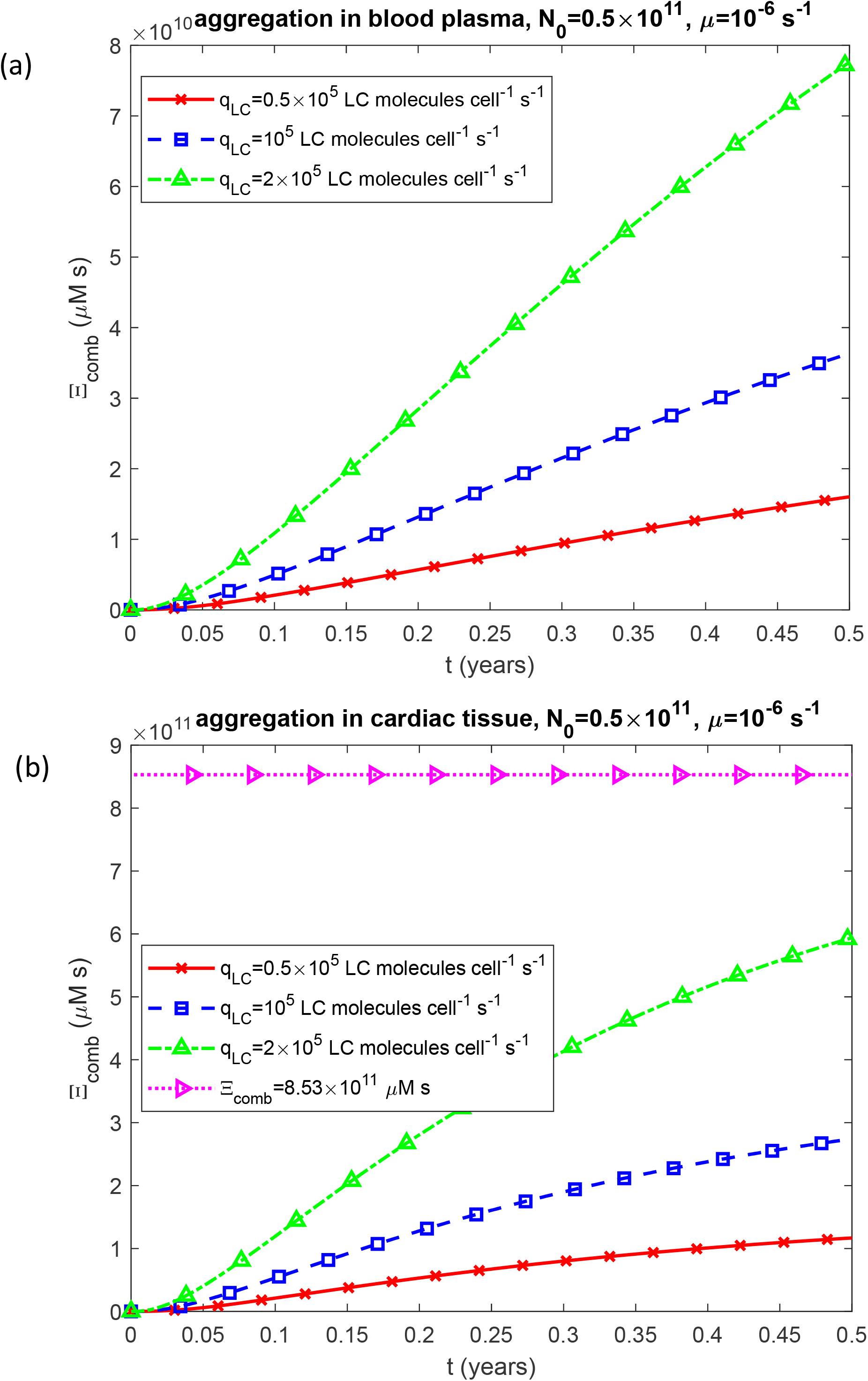
Time-dependent criterion characterizing the combined effect of cytotoxicity and myocardial stiffening, Ξ _*comb*_, vs time, *t*, for two scenarios: (a) aggregation occurring in the blood plasma and (b) aggregation occurring within the cardiac tissue. Results are presented for three different values of the LC secretion rate per clonal plasma cell, *q*_*LC*_ . *k*_1_ =10^−6^ s^-1^, *k*_2_ = 10^−6^ *μ*M^-1^ s^-1^, *k*_*u*_ = 4 ×10^−4^ s^-1^, *θ*_1/ 2,*B*_ = 10^7^ s, and *θ*_1/ 2,*I*_ = 10^7^ s. Results correspond to simulations with anticancer therapy.

The concentration of unfolded LC monomers susceptible to aggregation declined sharply, nearly disappearing within the first month (Figs. S28 and S29). The concentration of free oligomers initially increased during the first month due to conversion of unfolded monomers into oligomers but subsequently decreased as new oligomer formation became negligible and the remaining oligomers were progressively incorporated into fibrils (Fig. S30).

The concentration of oligomers incorporated into fibrils continued to rise over the six-month period, indicating ongoing deposition of free oligomers into fibrillar structures (Figs. S31 and S32). In Fig. S31, the concentration of fibril-incorporated oligomers circulating in the blood plasma approaches an asymptotic value at longer times as the oligomer population declines (Fig. S30a). The myocardial volume fraction occupied by fibrils also increased gradually over time; however, for a secretion rate of *q*_*LC*_ = 10^5^ LC molecules per plasma cell per second, this fraction reached only approximately 13% after six months in the scenario where LC aggregation occurred within cardiac tissue (Fig. S33).

Accumulated cardiotoxicity (Fig. S34) increased monotonically despite the decline in oligomer concentration after the first month (Fig. S30), because cardiotoxicity is defined as the time integral of oligomer concentration and therefore continues to accumulate even as oligomer levels decrease (Eqs. (27a) and (27b)).

Overall, anticancer therapy slowed the progression of the combined parameter representing cardiac damage, Ξ _*comb*_, such that after six months it did not reach the threshold associated with heart failure, even for the case of LC aggregation within the myocardium (Fig. 12). For *q*_*LC*_ = 10^5^ LC molecules per plasma cell per second, the corresponding biological age after six months was 77 years—well below the threshold of 100 years associated with patient death (Fig. S35). Thus, although anticancer therapy substantially mitigates cardiac damage by reducing the number of clonal plasma cells, residual damage from LC aggregation remains significant, albeit markedly less than in the absence of therapy.

Table 3 categorizes the model parameters into three groups based on the sensitivity of the combined cardiac damage criterion, Ξ_*comb*_, to each parameter. The combined cardiac damage criterion, Ξ_*comb*_, shows high sensitivity to several model parameters: the kinetic constant governing anticancer treatment–induced killing of clonal plasma cells, *μ* ; total myocardial tissue volume, *V*_*heart*_ ; the LC secretion rate per clonal plasma cell, *q*_*LC*_ ; the half-life of unfolded LC monomers in the blood plasma, *T*_1/ 2,*U, plasma*_ ; and, for scenario 2 only, the mass-transfer coefficient governing the transfer of unfolded LC monomers to the endocardial surface, *h*_*U*_ . Moderate sensitivity is observed for *k*_1_, *k*_2_, *k*_*cl*_, *k*_*u*_, *T*_1/ 2, *B*_, *T*_1/ 2, *I*_, *θ*_1/ 2,*B*_, and *θ*_1/ 2,*I*_, while Ξ_*comb*_ shows low sensitivity to *T*_1/ 2,*U,tissue*_ .

**Table 3.** Sensitivity of the criterion characterizing the combined effects of cytotoxicity and myocardial stiffening to various model parameters at the end of the six-month modeled terminal cardiac-progression interval. Simulations were performed using the following baseline parameter values: *k*_1_ =10^−6^ s^-1^, *k*_2_ = 10^−6^ *μ*M^-1^ s^-1^, *k*_*cl*_ = 4.81×10^−5^ s^−1^, *k*_*u*_ = 4 ×10^−4^ s^-1^, 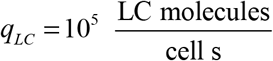, *T*_1/ 2,*U, plasma*_ = 1.44 × 10^4^ s, *T*_1/ 2, *B*_ = 6.5 × 10^7^ s, *T*_1/ 2, *I*_ = 6.5 × 10^7^ s, *T*_1/ 2,*U,tissue*_ = 6.5 × 10^7^ s, *θ*_1/ 2, *B*_ = 10^7^ s, *θ*_1/ 2,*I*_ = 10^7^ s, and *h*_*U*_ = 5×10^−9^ m s^-1^.

| Parameter | The scenario involving LC aggregation in the plasma. Dimensionless sensitivity of $\Xi_{comb}$ to the parameter given in this row at the end of the six-month modeled terminal cardiac-progression interval. $\mu=0$ (no anticancer treatment). | The scenario involving LC aggregation in the plasma. Dimensionless sensitivity of $\Xi_{comb}$ to the parameter given in this row at the end of the six-month modeled terminal cardiac-progression interval. With anticancer treatment, $\mu=10^{-6} \text{ s}^{-1}$. | The scenario involving LC aggregation in the cardiac tissue. Dimensionless sensitivity of $\Xi_{comb}$ to the parameter given in this row at the end of the six-month modeled terminal cardiac-progression interval. $\mu=0$ (no anticancer treatment). | The scenario involving LC aggregation in the cardiac tissue. Dimensionless sensitivity of $\Xi_{comb}$ to the parameter given in this row at the end of the six-month modeled terminal cardiac-progression interval. With anticancer treatment, $\mu=10^{-6} \text{ s}^{-1}$. |
| --- | --- | --- | --- | --- |
| Parameters to which $\Xi_{comb}$ exhibits high sensitivity | | | | |
| $\mu$ | 0 | -1.30 | 0 | -1.39 |
| $q_{LC}$ | 1.04 | 1.13 | 1.04 | 1.16 |
| $V_{heart}$ | N/A | N/A | -0.995 | -0.981 |
| $T_{1/2,U,plasma}$ | 0.0460 | 0.147 | 0.784 | 0.906 |
| $h_U$ | N/A | N/A | 0.740 | 0.729 |
| Parameters to which $\Xi_{comb}$ exhibits moderate sensitivity | | | | |
| $k_1$ | 0.00530 | 0.0185 | 0.00447 | 0.0185 |
| $k_2$ | 0.0408 | 0.128 | 0.0435 | 0.159 |
| $k_{cl}$ | -0.112 | -0.121 | -0.112 | -0.124 |
| $k_u$ | 0.112 | 0.121 | 0.113 | 0.124 |
| $T_{1/2,B}$ | 0.0227 | 0.0565 | 0.0233 | 0.0627 |
| $T_{1/2,I}$ | 0.00724 | 0.0381 | 0.0155 | 0.0503 |
| $\theta_{1/2,B}$ | 0.0227 | 0.0565 | 0.0233 | 0.0627 |
| $\theta_{1/2,I}$ | 0.00724 | 0.0381 | N/A | N/A |
| Parameters to which $\Xi_{comb}$ exhibits low sensitivity | | | | |
| $T_{1/2,U,tissue}$ | N/A | N/A | $2.06 \times 10^{-6}$ | $7.63 \times 10^{-6}$ |

## 4. Discussion, limitations of the model, and future directions

AL amyloidosis is a progressive and dynamic disease (Eisele et al. 2015) characterized by the expansion of clonal plasma cell populations, production of folded LC monomers by these cells, conversion of folded LCs into unfolded aggregation-prone monomers, aggregation of unfolded monomers into oligomers, and subsequent deposition of oligomers into amyloid fibrils. Depending on the site of aggregation, either unfolded LC monomers or LC fibrils infiltrate the cardiac tissue, contributing to increased myocardial stiffness. In addition, LC oligomers are considered the primary mediators of cardiotoxicity. Consequently, two pathological processes act concurrently to damage the heart: structural stiffening due to LC fibril infiltration and direct cardiotoxicity induced by LC oligomers.

Given the rapid progression of AL amyloidosis, early diagnosis is critical (Palladini et al. 2020), underscoring the importance of understanding the disease’s underlying dynamic processes.

The mathematical framework developed here shares several features with models of amyloid aggregation developed for Alzheimer’s disease and other protein-misfolding disorders. In particular, both classes of models represent aggregation as a nonlinear kinetic process involving the generation of aggregation-competent species followed by autocatalytic amplification. The Finke–Watzky formulation used here represents this behavior through two effective steps: primary formation of oligomers and their subsequent autocatalytic conversion from monomeric LC species. This minimal representation has also been applied to other amyloid and prion aggregation systems and provides a compact description of sigmoidal aggregation kinetics.

More detailed kinetic models of amyloid-β aggregation explicitly distinguish among primary nucleation, fibril elongation, monomer-dependent secondary nucleation, and fibril fragmentation (Knowles et al. 2009; Meisl et al. 2014). In particular, combined experimental and theoretical studies have shown that fibril-catalyzed secondary nucleation can become a dominant pathway for the generation of new Aβ aggregates (Cohen et al. 2013; Meisl et al. 2014). For Aβ42, this mechanism also provides an important link between the accumulation of mature fibrils and the production of soluble oligomeric species, because existing fibril surfaces catalyze the formation of new oligomers from monomeric peptide (Cohen et al. 2013; Michaels et al. 2020).

The present model differs from these Aβ frameworks in two important respects. First, the aggregation substrate is not produced at a fixed rate but originates from a pathogenic plasma-cell clone. LC production therefore provides an upstream source term that couples hematologic disease to protein aggregation. Second, the model explicitly represents transport/deposition between compartments and distinguishes aggregation occurring in the blood plasma from aggregation occurring within cardiac tissue. This compartmental structure allows the model to examine how the effective distribution volume influences local LC concentration and, through the autocatalytic aggregation term, the rate of oligomer formation.

A further difference is that AL amyloidosis involves immunoglobulin light chains whose native folded state must undergo destabilization or unfolding before efficient aggregation can occur. Thus, the present model includes an explicit folded-to-unfolded LC conversion step upstream of aggregation. In contrast, many Aβ models begin with an aggregation-competent peptide pool and focus primarily on the subsequent self-assembly processes.

The simplicity of the F–W representation is both an advantage and a limitation. It permits coupling of aggregation kinetics to plasma-cell production, systemic clearance, tissue deposition, and cardiac injury without introducing a large number of poorly constrained microscopic rate constants. At the same time, the model does not explicitly resolve the molecular heterogeneity of LC oligomers, fibril lengths, fragmentation, or distinct secondary-nucleation mechanisms. More detailed descriptions analogous to those developed for Aβ could be incorporated when sufficient quantitative LC-specific kinetic data become available.

The present framework is conceptually related to mathematical models previously developed for Aβ aggregation, in which the dynamics of soluble monomers, oligomeric intermediates, and fibrillar aggregates are used to relate microscopic aggregation processes to the accumulation of Aβ at larger scales (Pallitto and Murphy 2001; Craft et al. 2002; Michaels et al. 2020). More detailed kinetic descriptions explicitly distinguish primary nucleation, fibril elongation, fibril-catalyzed secondary nucleation, and fragmentation, thereby providing a mechanistic representation of the nonlinear amplification of amyloid aggregates (Knowles et al. 2009; Cohen et al. 2013; Meisl et al. 2014). The present AL model adopts a more reduced description of the aggregation process because its primary objective is to couple LC aggregation to upstream production by the plasma-cell clone and downstream cardiac deposition and injury. Nevertheless, the common kinetic structure of these aggregation frameworks suggests that the present model could be extended to incorporate additional microscopic pathways as sufficiently detailed LC-specific experimental data become available.

Experimental studies of amyloidogenic LCs indicate that fibril formation can proceed through multiple molecular pathways involving distinct non-native conformations and oligomeric intermediates, and that the resulting aggregation kinetics depend strongly on LC sequence, thermodynamic stability, and domain interactions (Rennella, Morgan, Yan, et al. 2019; Kazman et al. 2021; Pradhan et al. 2023; Wong et al. 2023). In vitro experiments have further demonstrated a nonlinear dependence of LC aggregation kinetics on protein concentration and acceleration of fibril formation by pre-existing fibrillar seeds (Blancas-Mejía and Ramirez-Alvarado 2016; Wong et al. 2023). The latter observation is consistent with an autocatalytic contribution to LC fibril formation, in which existing aggregates promote the recruitment of soluble LC species into additional fibrillar material (Blancas-Mejía and Ramirez-Alvarado 2016). Taken together, the pronounced sequence- and conformation-dependent variability observed experimentally suggests that the kinetic coefficients employed in a reduced-order description should be interpreted as effective, LC-specific parameters rather than universal constants (Rennella, Morgan, Yan, et al. 2019; Rennella, Morgan, Kelly, et al. 2019; Pradhan et al. 2023; Wong et al. 2023).

The model predicts that LC aggregation is highly nonlinear during the modeled terminal cardiac-progression interval. Oligomer concentrations remain relatively low during the initial portion of this interval and subsequently increase rapidly as autocatalytic oligomer formation becomes dominant. This result should be interpreted only as a prediction of the late-stage aggregation dynamics represented by the model and should not be extrapolated to the earlier, unmodeled course of AL amyloidosis.

The model indicates that when LC aggregation occurs within cardiac tissue, the fraction of the myocardium occupied by LC fibrils is approximately 30 times greater than when aggregation occurs in blood plasma. This difference highlights the importance of compartmentalization in nonlinear aggregation kinetics. Because the F–W autocatalytic term depends on the local concentration of aggregation-competent species, the substantially smaller volume of cardiac tissue increases local oligomer concentrations and thereby accelerates autocatalytic oligomer formation and subsequent fibril accumulation, even when the total amount of secreted LC is unchanged. Thus, in the present model, the site of aggregation affects the kinetics through a concentration-dependent positive-feedback mechanism rather than serving merely as a geometric parameter. Consistent with the higher oligomer concentrations in cardiac tissue, the criterion representing accumulated oligomer-induced cardiotoxicity is approximately fivefold greater than in the plasma-aggregation scenario.

A criterion is proposed to quantify the combined cardiac damage resulting from oligomer-induced cardiotoxicity and myocardial stiffening. This criterion attains values approximately tenfold higher when LC aggregation occurs within the cardiac tissue compared with aggregation in the blood plasma. It is further proposed that this criterion may be used as a measure of biological age.

An interesting finding is the near independence of Ξ _*comb*_ from *θ*_1/2,*B*_ when LC aggregation occurs in blood plasma. To explain this behavior, the two components of Ξ _*comb*_ were examined separately. The cardiotoxicity component Ξ increases with *θ*_1/ 2,*B*_ because longer oligomer residence times elevate their concentrations and toxicity, whereas the myocardial volume fraction occupied by fibrils *V*_*fibrils*_ / *V*_*heart*_, which characterizes cardiac stiffening, decreases as oligomers convert into fibrils more slowly. These opposing effects nearly cancel each other, making the combined cardiac damage criterion Ξ _*comb*_ effectively independent of *θ*_1/ 2,*B*_ .

The scenario involving anticancer therapy markedly slowed the increase of the combined parameter representing cardiac damage Ξ _*comb*_ . The predicted beneficial effect of reducing the plasma-cell population is consistent with the central therapeutic principle of AL amyloidosis, in which suppression of the pathogenic plasma-cell clone reduces the circulating supply of amyloidogenic LC. Experimental and clinical observations that normalization of circulating free LC concentrations is associated with subsequent regression or stabilization of amyloid deposits provide biological support for treating LC production as an upstream determinant of tissue amyloid burden (van Gameren et al. 2009). However, Ξ _*comb*_ remains appreciable—though substantially reduced compared with the untreated case—indicating that treatment should be initiated as early as possible to minimize residual cardiac injury resulting from LC aggregation.

The sensitivity analysis identifies a relatively small group of parameters as the principal determinants of the combined cardiac damage criterion, Ξ_*comb*_ . These include the efficacy of clone-directed therapy, represented by the plasma-cell killing rate constant *μ* ; myocardial tissue volume *V*_*heart*_ ; LC secretion by individual clonal plasma cells *q*_*LC*_ ; the plasma half-life of unfolded LC monomers *T*_1/ 2,*U, plasma*_ ; and, specifically for scenario 2, the mass-transfer coefficient controlling the delivery of unfolded LC monomers to the endocardial surface *h*_*U*_ . The pronounced sensitivity to the plasma-cell killing rate and LC secretion rate highlights the importance of clone-directed therapy and precursor supply in determining cardiac damage during the modeled interval, further emphasizing their relevance as therapeutic targets. The findings presented here provide a theoretical framework for identifying model parameters and dynamic variables that may be informative for future biomarker development. However, validation against longitudinal clinical measurements of circulating free LC concentrations, cardiac biomarkers, and tissue amyloid burden will be required before the proposed cardiac-damage criterion can be considered a clinically useful biomarker.

Several limitations should be noted. First, AL amyloidosis is highly heterogeneous, and each patient possesses a structurally distinct pathogenic LC with potentially different folding stability, aggregation propensity, and tissue tropism. The present model therefore represents an effective amyloidogenic LC rather than a specific patient-derived sequence. Future studies could address this heterogeneity by tailoring parameters governing LC folding and aggregation to individual patients or by explicitly simulating distinct LC variants. Second, the plasma-cell clone is represented using a phenomenological Gompertz-type function. This formulation is not intended to reproduce the clonal biology of multiple myeloma or to imply myeloma-like proliferation in AL amyloidosis; rather, it provides an effective time-dependent source term for LC production. Future models incorporating longitudinal measurements of clonal plasma-cell burden and LC production could replace this phenomenological representation with patient-specific plasma-cell kinetics. Third, the model focuses on the advanced cardiac-progression phase and does not reconstruct the potentially prolonged preclinical period between the emergence of the pathogenic plasma-cell clone and clinically overt cardiac involvement. Accordingly, the six-month interval used for model calibration represents a characteristic terminal cardiac-progression timescale and should not be interpreted as the total duration of AL amyloidosis. Fourth, LC clearance is represented by an effective systemic clearance constant and does not explicitly resolve individual physiological clearance pathways or differences between κ and λ LCs. Finally, cardiac volume is represented by a population-level average rather than a patient-specific value. Future studies could address this limitation by incorporating patient-specific myocardial mass or volume obtained from echocardiography or cardiac magnetic resonance imaging.

## Abbreviations

AL: amyloid light chain
dFLC: difference between the plasma concentrations of involved and uninvolved free LCs
F–W: Finke–Watzky
Ig: immunoglobulin
LC: light chain

## Acknowledgment

This research was supported by the National Science Foundation (grant DMS-2451660) and the Alexander von Humboldt Foundation through the Humboldt Research Award.

## Supplemental Materials

### S1. Gompertzian function

When *μ* = 0 in Eq. (1), corresponding to the absence of anticancer treatment, the solution of Eq. (1) with initial condition (2) takes the form of the Gompertzian function:

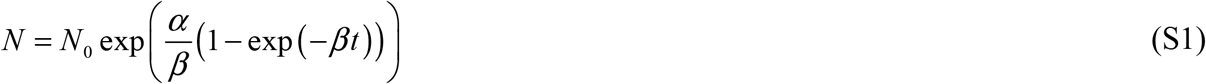

In the absence of anticancer treatment Eqs. (1) and (2) can be reformulated as (Sullivan and Salmon 1972):

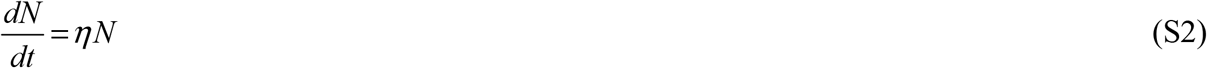

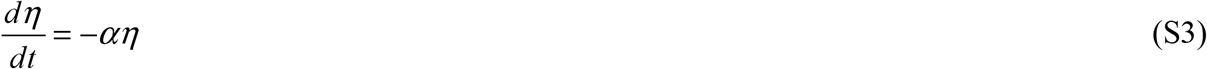

Eqs. (S2) and (S3) are solved subject to the following initial conditions:

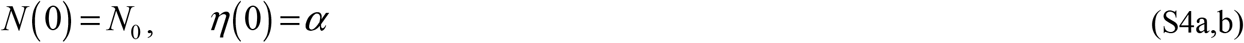

The solution of Eqs. (S2)–(S4) is expressed by Eq. (S1), which gives *N* (*t*), together with the following solution for *η*(*t*) :

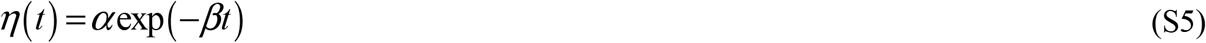

Accordingly, Eq. (S2) can be reformulated as:

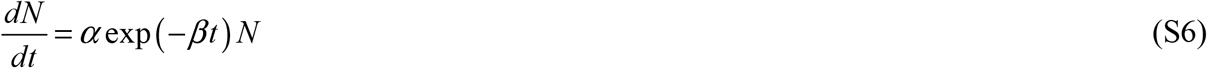

to be solved under the initial condition (S4a).

Eq. (S6) demonstrates that, in the Gompertzian model, the exponential decay rate—equal to *α* at *t* = 0— decreases exponentially at a constant rate *β* .

### S2. Numerical solution

Algebraic Eq. (7) was solved numerically using the FZERO root-finding algorithm in MATLAB (R2024a, MathWorks, Natick, MA, USA). Initial value problems for ordinary differential equations were addressed using MATLAB’s ODE45 solver (R2024a, MathWorks, Natick, MA, USA). Specifically, Eq. (1) was integrated with initial condition (2), and Eq. (9) was solved with initial condition (10). For scenario 1, Eqs. (13)–(16) were solved under initial conditions (17), whereas for scenario 2, Eqs. (18)– (21) were solved with initial conditions (22). The ODE45 solver, which implements an adaptive step-size Runge–Kutta method, was selected for its balance between accuracy and computational efficiency. To ensure high numerical precision, both the relative and absolute tolerances were set to 1e-10.

### S3. Supplementary figures

**Fig. S1.**
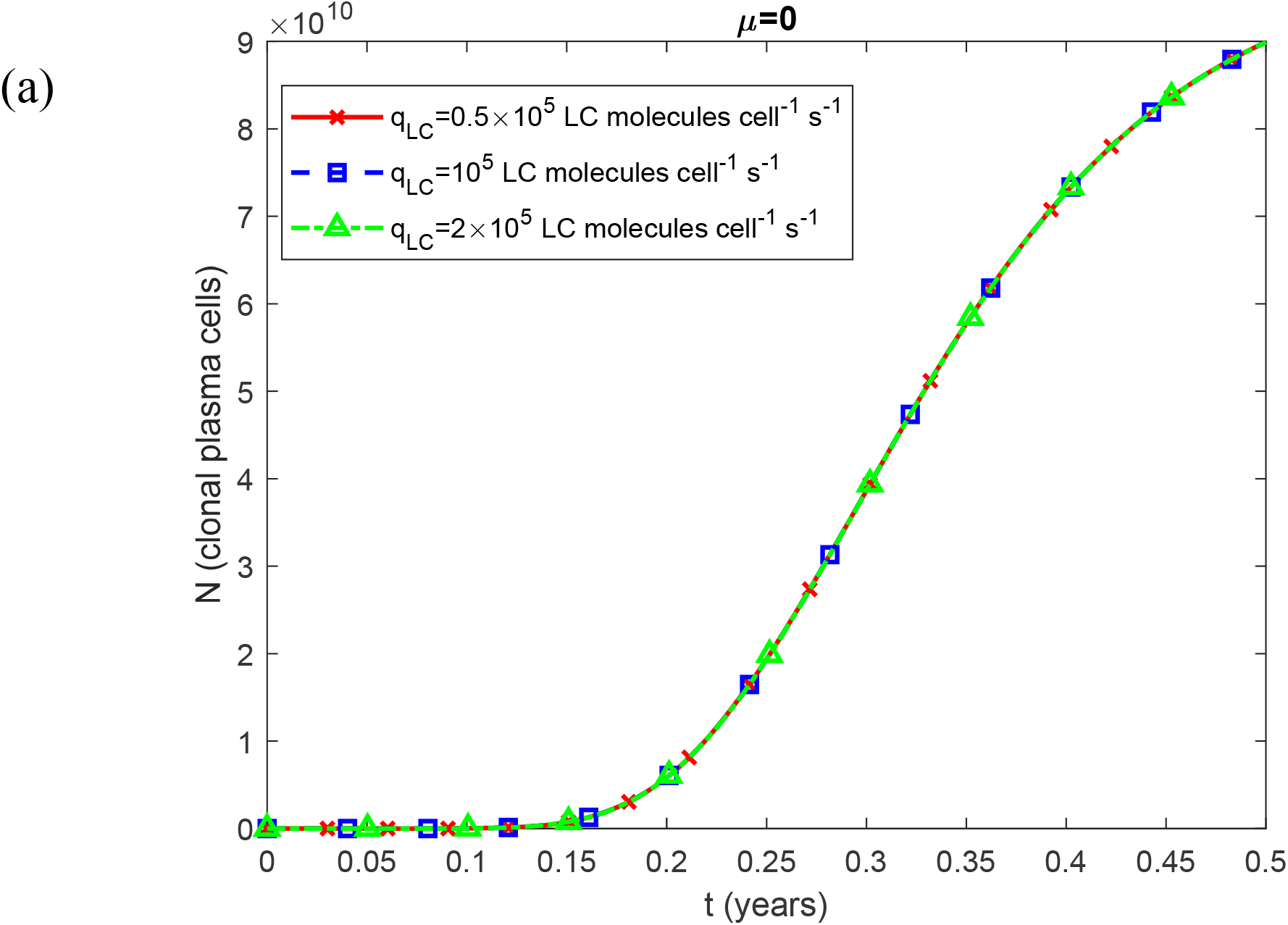
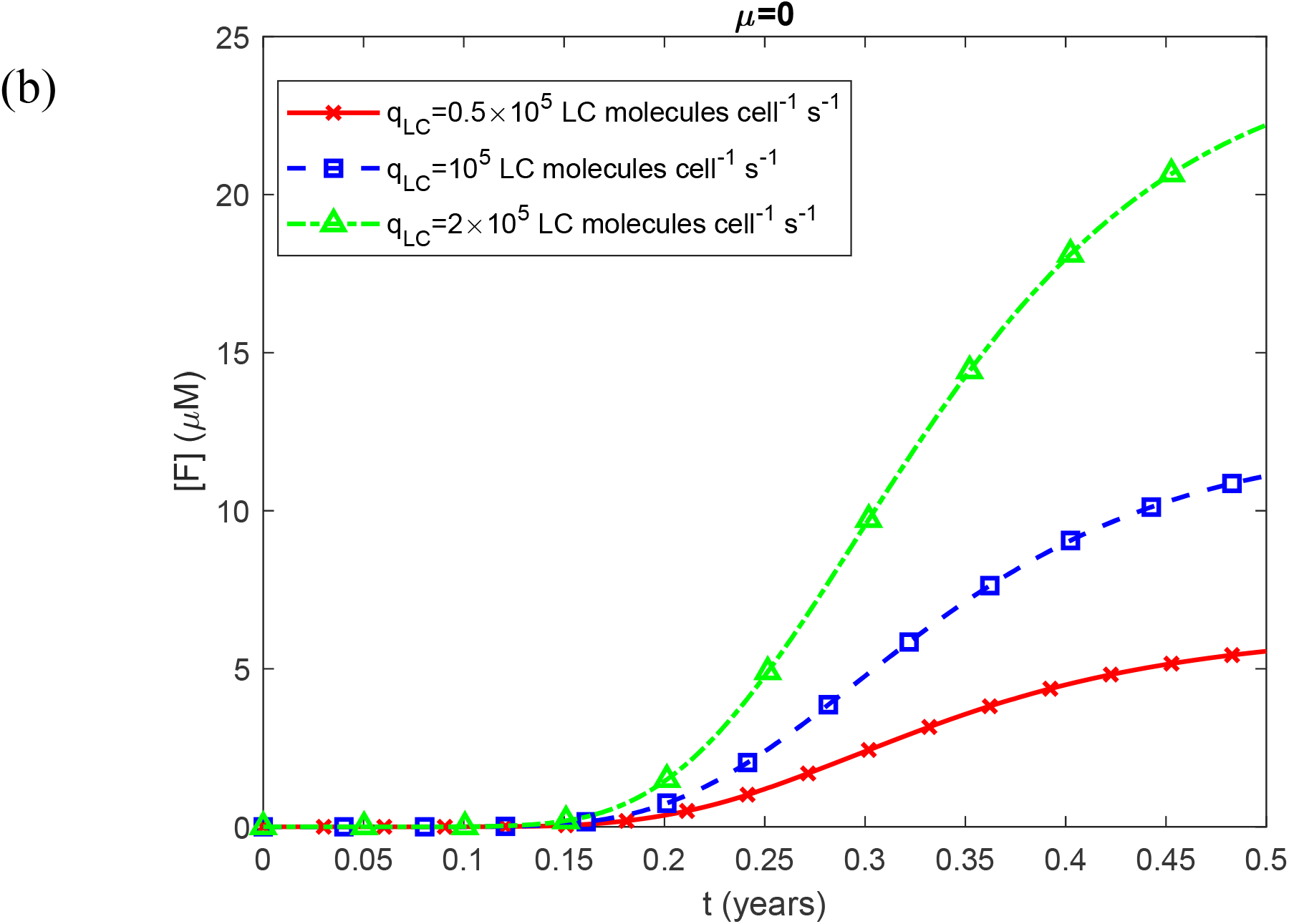
(a) Number of clonal plasma cells, *N*, vs time, *t*. The results indicate that *N* is independent of *q* _*LC*_ . (b) Plasma molar concentration of natively folded Ig LC monomers secreted by clonal plasma cells vs time for three different values of the secretion rate of folded LCs per single plasma cell belonging to a clonal population, *q* _*LC*_ . Computations were performed for *k*_*u*_ = 4 ×10^−4^ s^-1^. Results correspond to simulations without anticancer therapy.

**Fig. S2.**
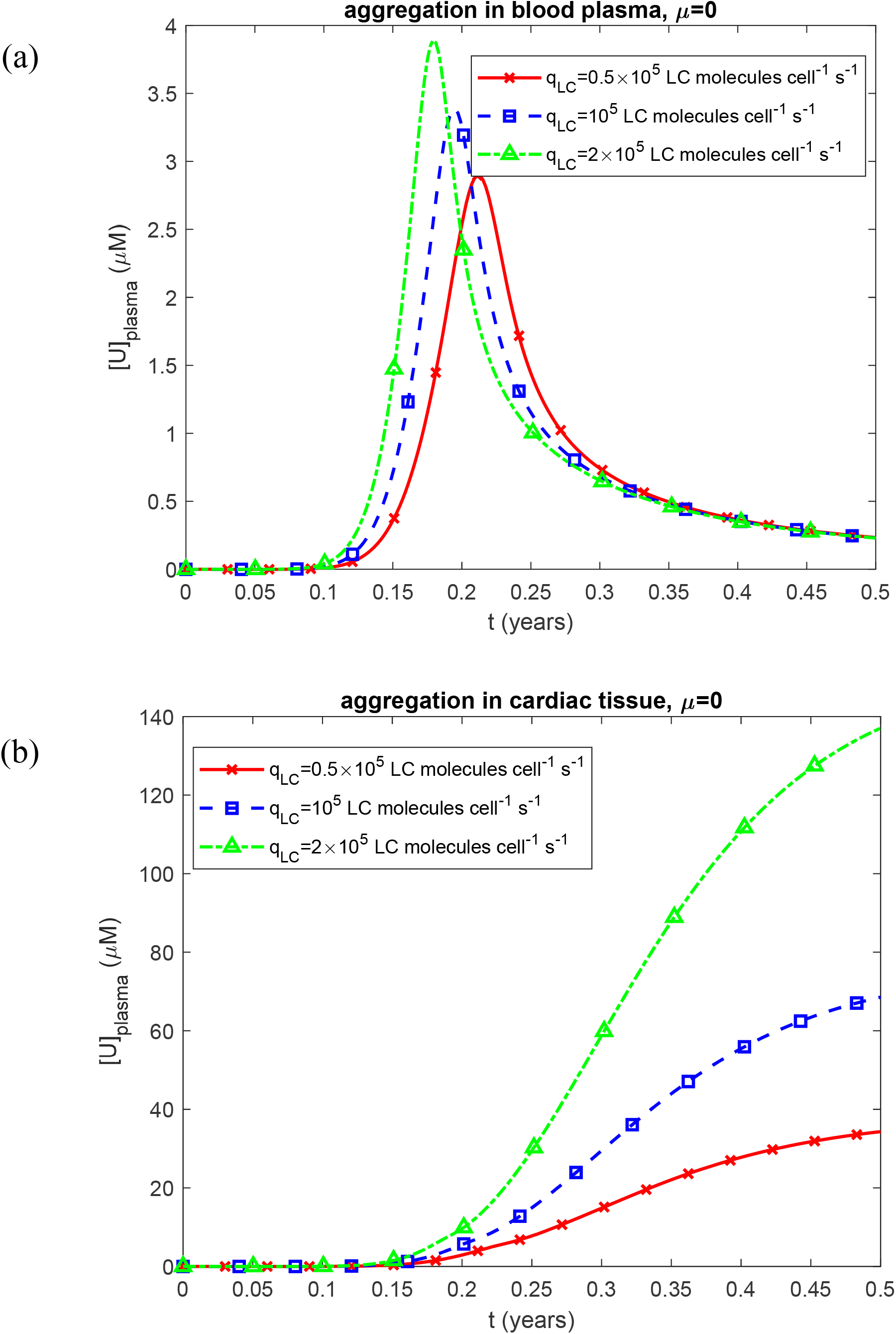
Plasma molar concentration of unfolded Ig LC monomers susceptible to aggregation, [*U*]_*plasma*_, vs time, *t*, for the scenarios of (a) aggregation occurring in the blood plasma and (b) aggregation occurring within the cardiac tissue. Results are shown for three different values of the LC secretion rate per clonal plasma cell, *q*_*LC*_ . Computations were performed for *k*_1_ =10^−6^ s^-1^, *k*_2_ = 10^−6^ *μ*M^-1^ s^-1^, *k*_*u*_ = 4 ×10^−4^ s^-1^, *θ*_1/ 2,*B*_ = 10^7^ s, and *θ*_1/ 2, *I*_ = 10^7^ s. Results correspond to simulations without anticancer therapy.

**Fig. S3.**
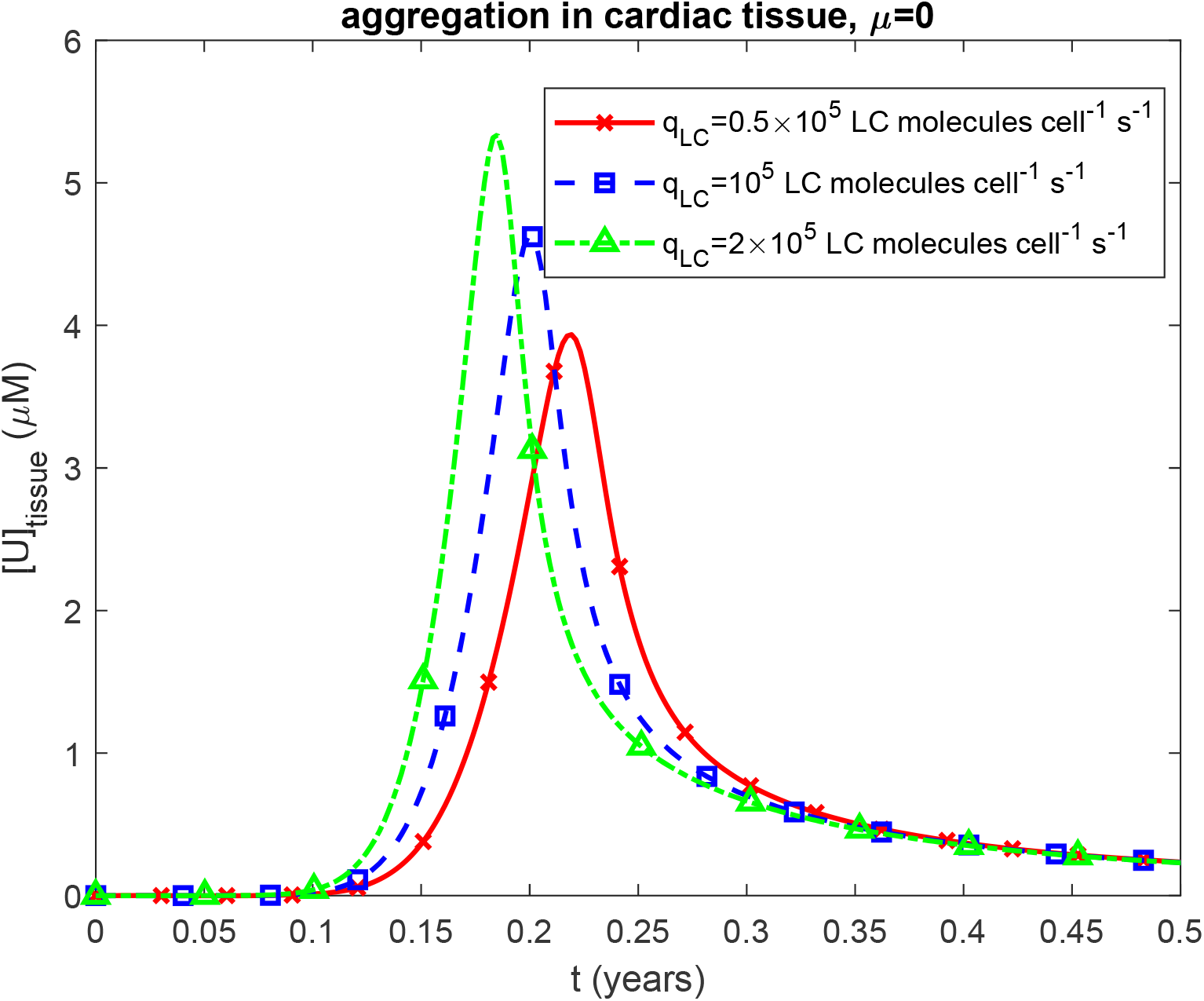
Molar concentration of unfolded Ig LC monomers in cardiac tissue, [*U*]_*tissue*_, vs time, *t*, for the scenario where aggregation occurs within the tissue. Results are shown for three different values of the LC secretion rate per clonal plasma cell, *q*_*LC*_ . Computations were performed for *k*_1_ =10^−6^ s^-1^, *k*_2_ = 10^−6^ *μ*M^-1^ s^-1^, *k*_*u*_ = 4 ×10^−4^ s^-1^, and *θ*_1/ 2,*B*_ = 10^7^ s. Results correspond to simulations without anticancer therapy.

**Fig. S4.**
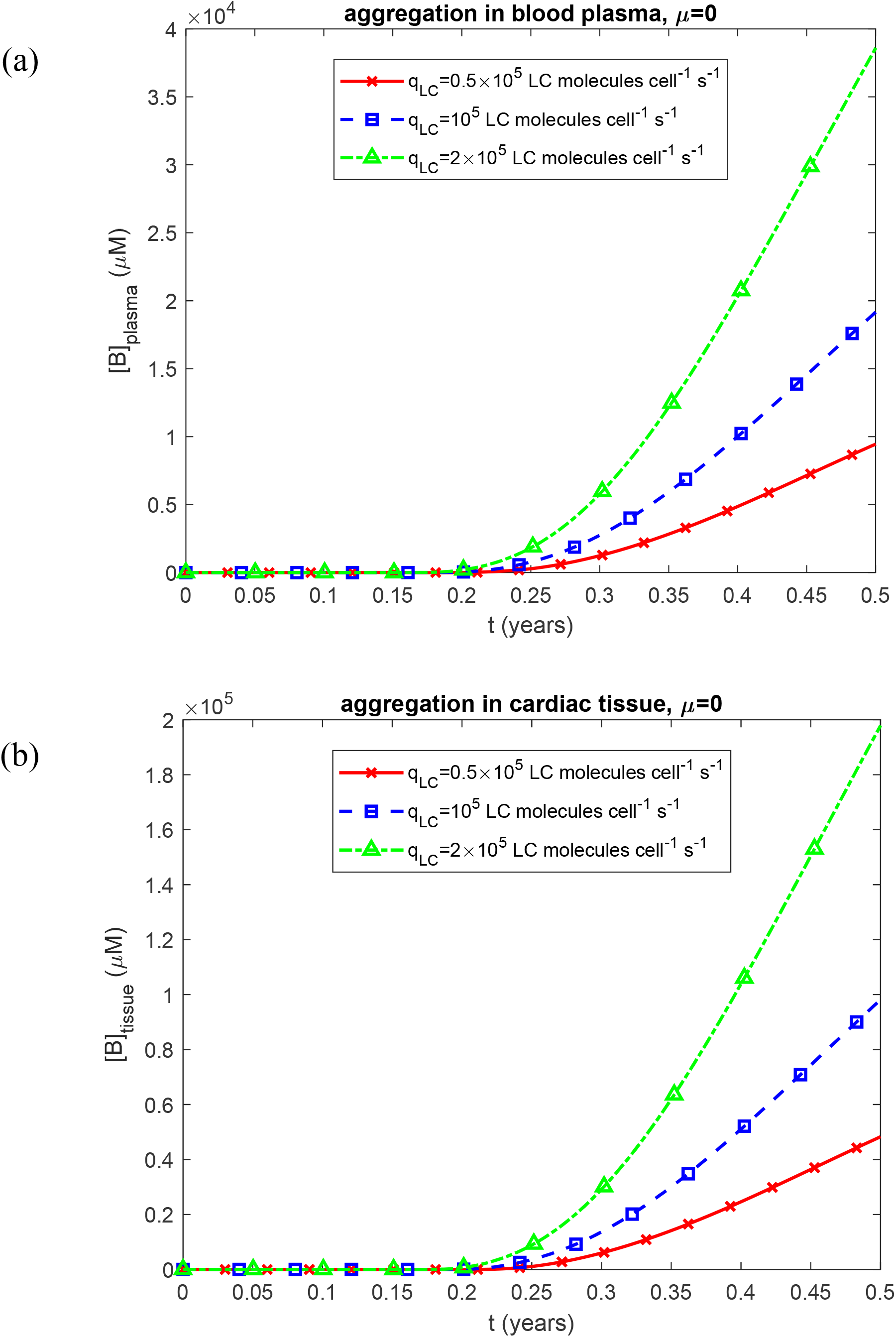
(a) Molar concentration of free LC oligomers in blood plasma, [*B*]_*plasma*_, vs time, *t*, when aggregation occurs in the plasma, and (b) molar concentration of free LC oligomers in cardiac tissue, [*B*]_*tissue*_, vs time, *t*, when aggregation occurs within the tissue. Results are shown for three different values of the LC secretion rate per clonal plasma cell, *q*_*LC*_ . Computations were performed for *k*_1_ =10^−6^ s^-1^, *k*_2_ = 10^−6^ *μ*M^-1^ s^-1^, *k*_*u*_ = 4 ×10^−4^ s^-1^, *θ*_1/ 2,*B*_ = 10^7^ s, and *θ*_1/ 2, *I*_ = 10^7^ s. Results correspond to simulations without anticancer therapy.

**Fig. S5.**
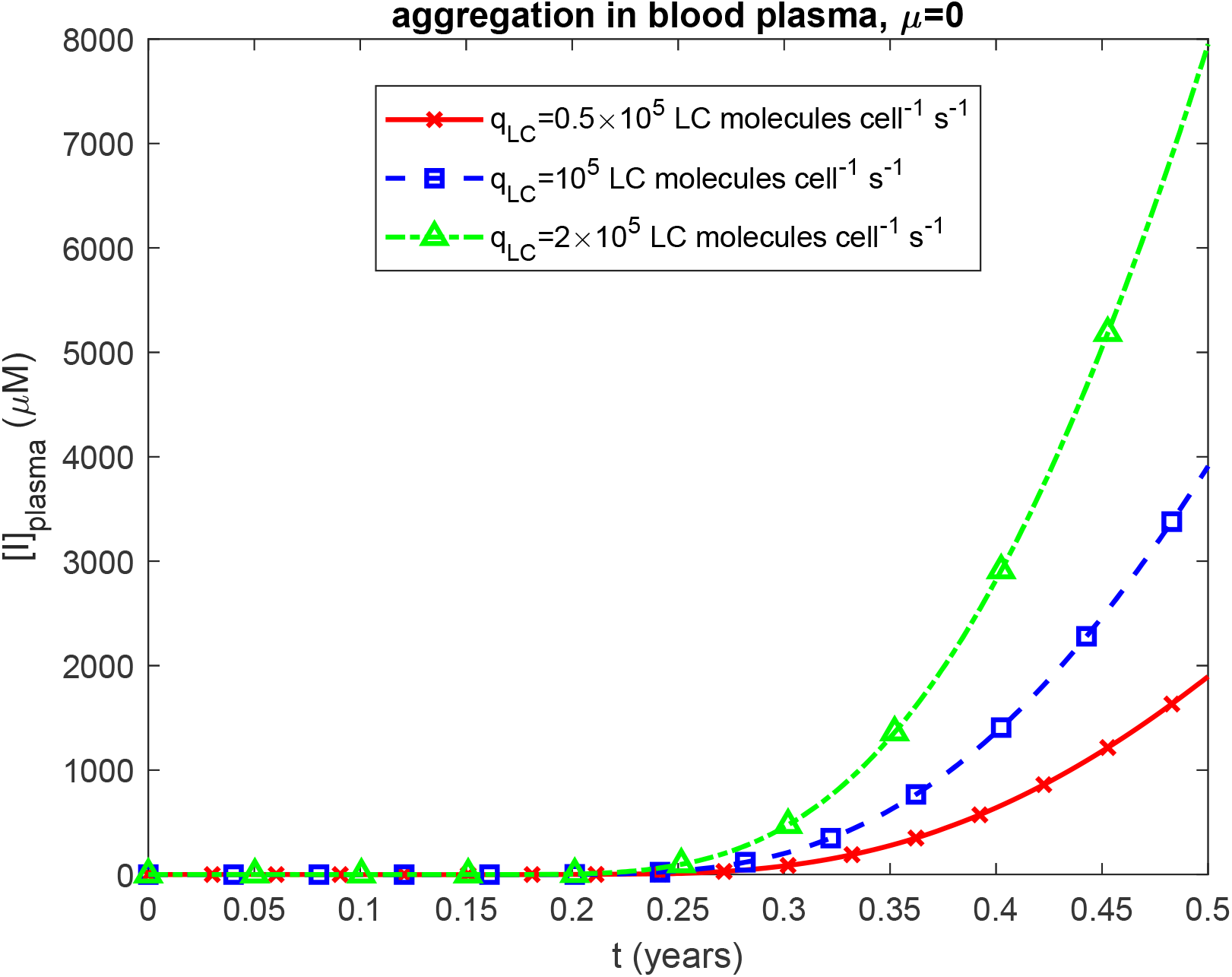
Molar concentration of LC oligomers deposited into amyloid protofibrils circulating within blood plasma, [*I*]_*plasma*_, vs time, *t*. Results are shown for three different values of the LC secretion rate per clonal plasma cell, *q*_*LC*_ . Computations were performed for *k*_1_ =10^−6^ s^-1^, *k*_2_ = 10^−6^ *μ*M^-1^ s^-1^, *k*_*u*_ = 4 ×10^−4^ s^-1^, *θ*_1/ 2,*B*_ = 10^7^ s, and *θ*_1/ 2, *I*_ = 10^7^ s. Results correspond to simulations without anticancer therapy.

**Fig. S6.**
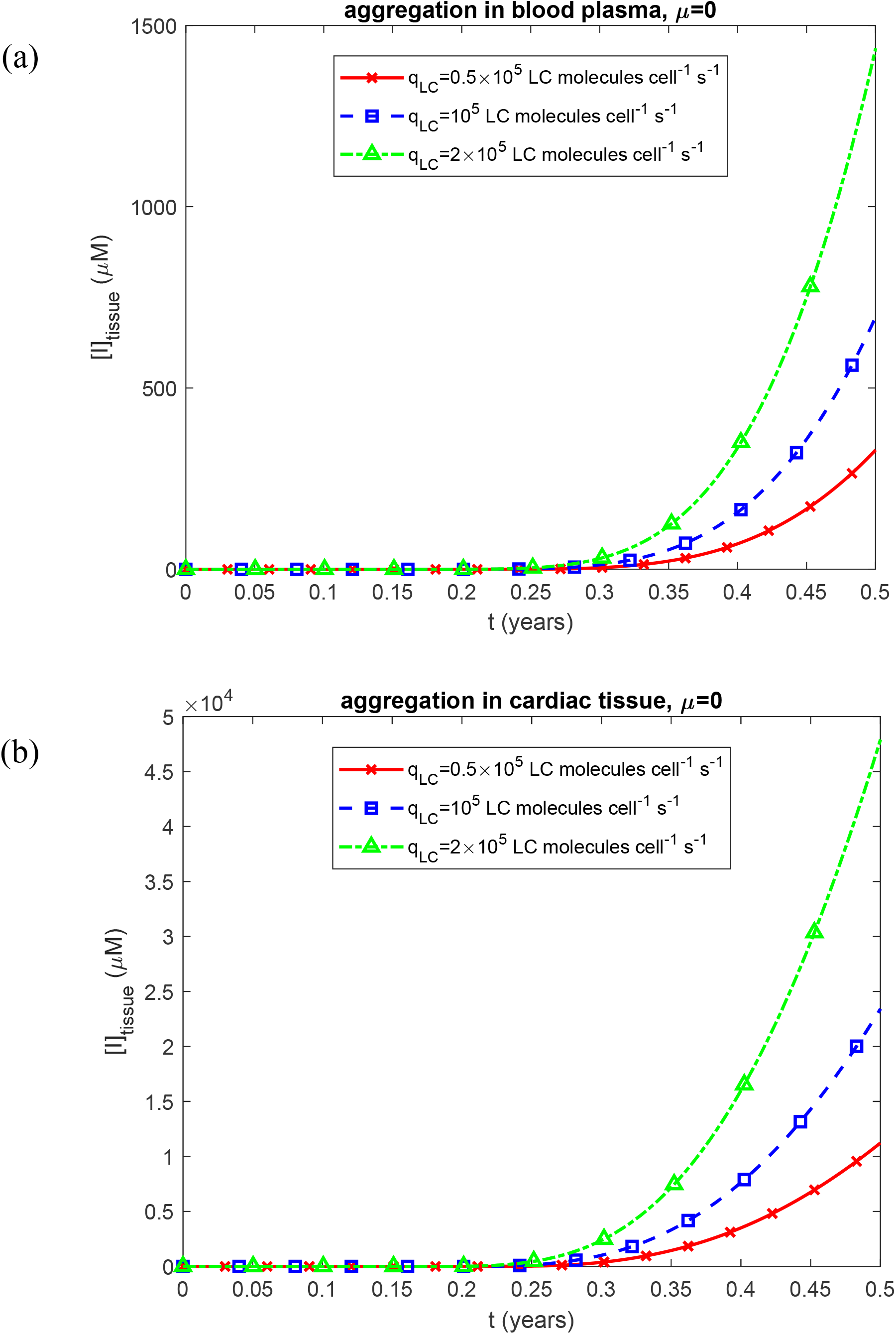
Molar concentration of LC oligomers incorporated into amyloid fibrils infiltrating the cardiac tissue, [*I*]_*tissue*_, vs time, *t*, for two scenarios: (a) aggregation occurring in the blood plasma and (b) aggregation occurring within the cardiac tissue. Results are shown for three different values of the LC secretion rate per clonal plasma cell, *q* _*LC*_ . Computations were performed for *k*_1_ =10^−6^ s^-1^, *k*_2_ = 10^−6^ *μ*M^-1^ s^-1^, *k*_*u*_ = 4 ×10^−4^ s^-1^, *θ*_1/ 2,*B*_ = 10^7^ s, and *θ*_1/ 2,*I*_ = 10^7^ s. Results correspond to simulations without anticancer therapy.

**Fig. S7.**
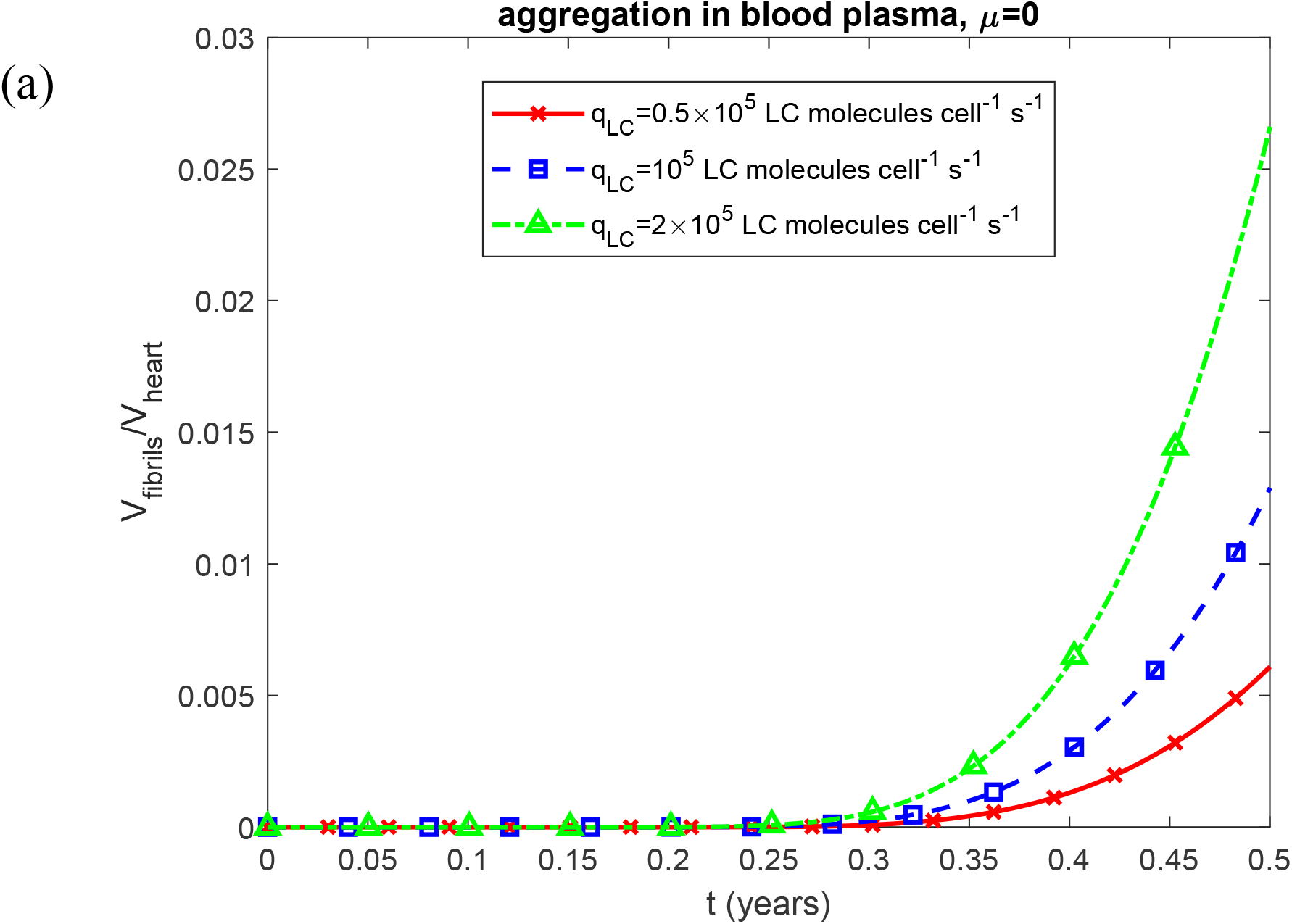
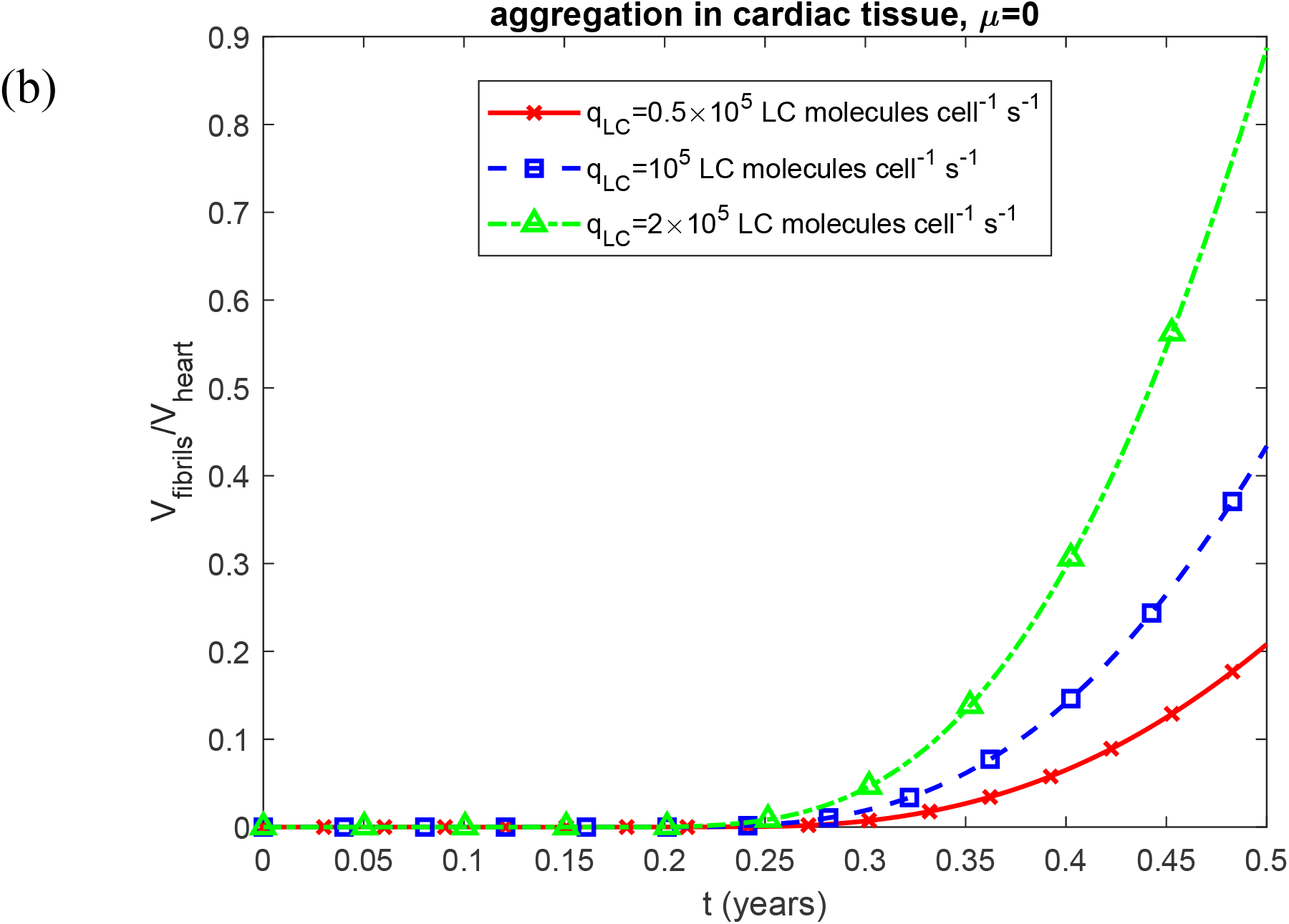
Fraction of myocardial volume occupied by LC fibrils, *V*_*fibrils*_ / *V*_*heart*_, vs time, *t*, for two scenarios: (a) aggregation occurring in the blood plasma and (b) aggregation occurring within the cardiac tissue. Results are presented for three different values of the LC secretion rate per clonal plasma cell, *q* _*LC*_ . Computations were performed for *k*_1_ =10^−6^ s^-1^, *k*_2_ = 10^−6^ *μ*M^-1^ s^-1^, *k*_*u*_ = 4 ×10^−4^ s^-1^, *θ*_1/ 2, *B*_ = 10^7^ s, and *θ*_1/ 2,*I*_ = 10^7^ s. Results correspond to simulations without anticancer therapy.

**Fig. S8.**
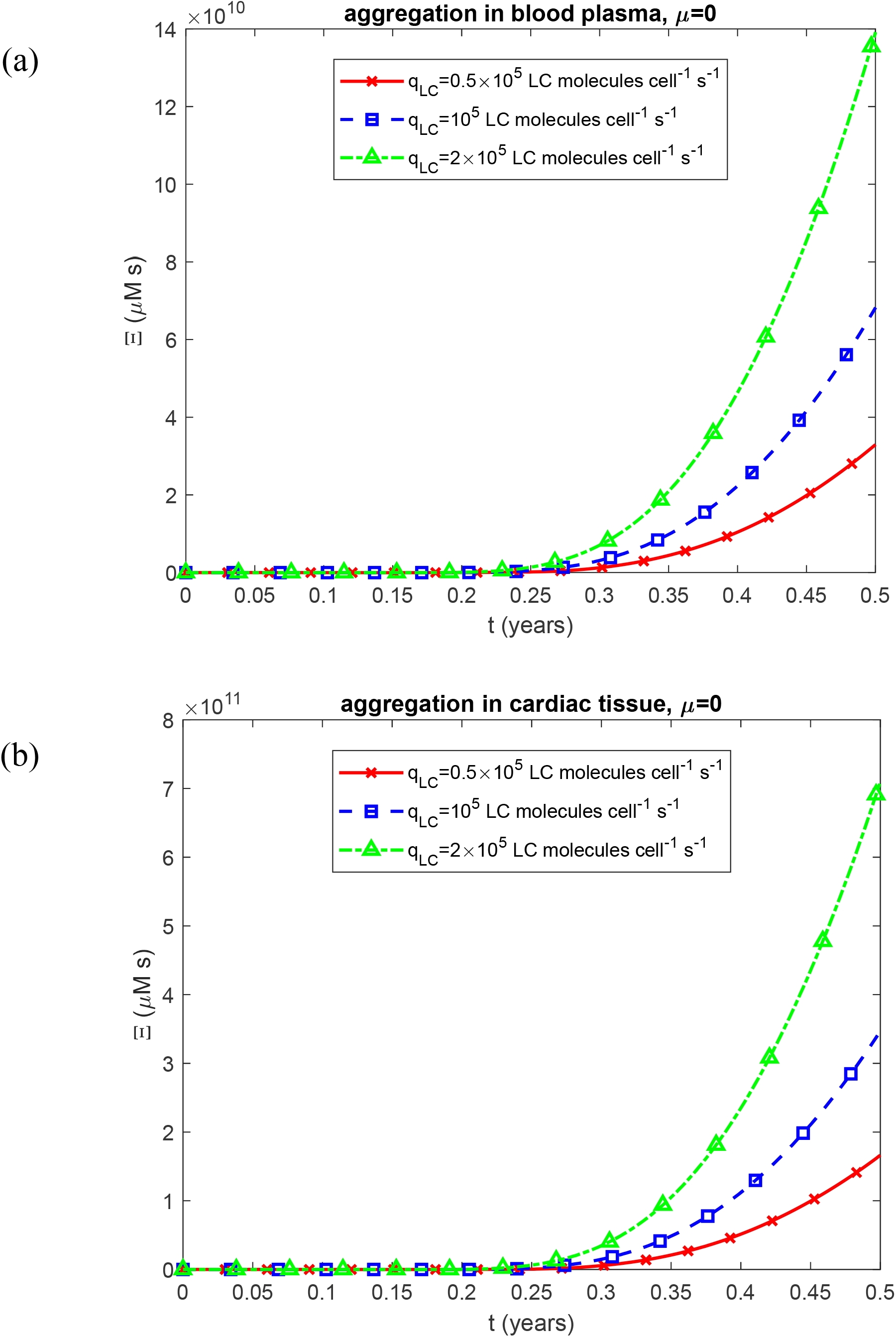
Time-dependent criterion for accumulated cardiotoxicity induced by LC oligomers, Ξ, vs time, *t*, for two scenarios: (a) aggregation occurring in the blood plasma and (b) aggregation occurring within the cardiac tissue. Results are presented for three different values of the LC secretion rate per clonal plasma cell, *q*_*LC*_ . Computations were performed for *k*_1_ =10^−6^ s^-1^, *k*_2_ = 10^−6^ *μ*M^-1^ s^-1^, *k*_*u*_ = 4 ×10^−4^ s^-1^, *θ* _1/ 2,*B*_ = 10^7^ s, and *θ*_1/ 2, *I*_ = 10^7^ s. Results correspond to simulations without anticancer therapy.

**Fig. S9.**
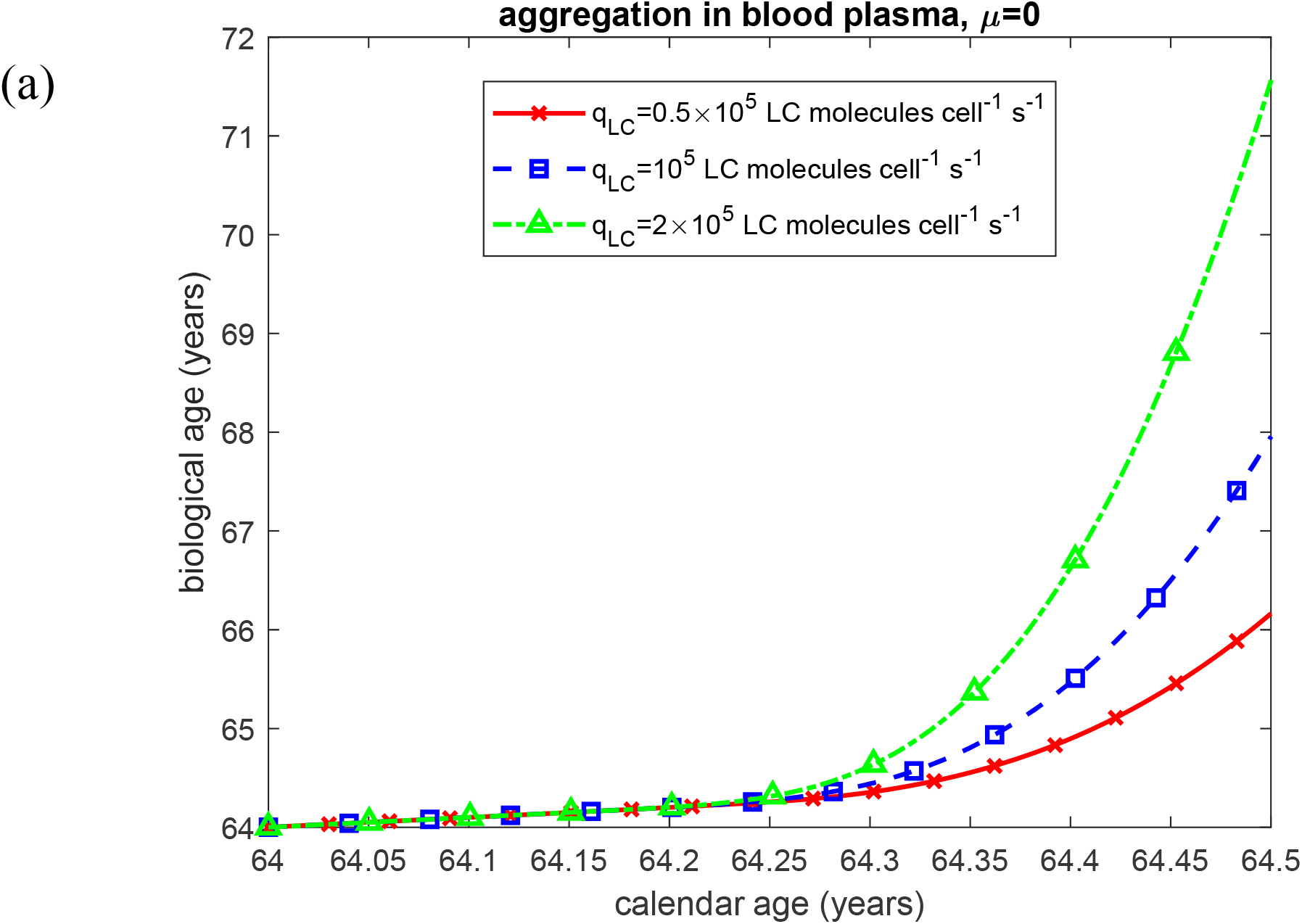
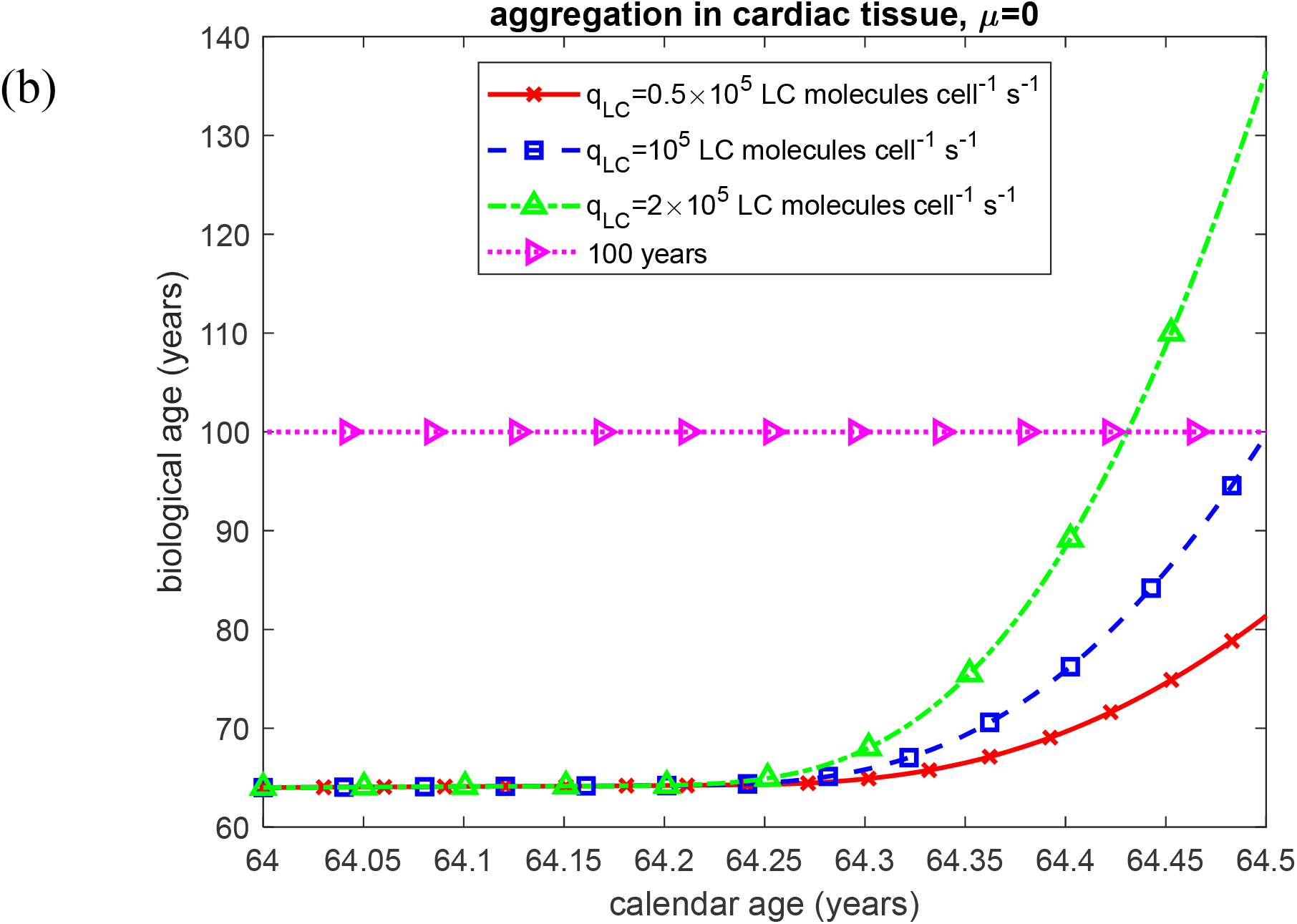
Biological age vs calendar age under two scenarios: (a) aggregation occurring in the blood plasma and (b) aggregation occurring within the cardiac tissue. Results are presented for three different values of the LC secretion rate per clonal plasma cell, *q*_*LC*_ . Computations were performed for *k*_1_ =10^−6^ s^-1^, *K*_2_ = 10^−6^ *μ*M^-1^ s^-1^, *k*_*u*_ = 4 ×10^−4^ s^-1^, *θ*_1/ 2,*B*_ = 10^7^ s, and *θ*_1/ 2,*I*_ = 10^7^ s. Results correspond to simulations without anticancer therapy.

**Fig. S10.**
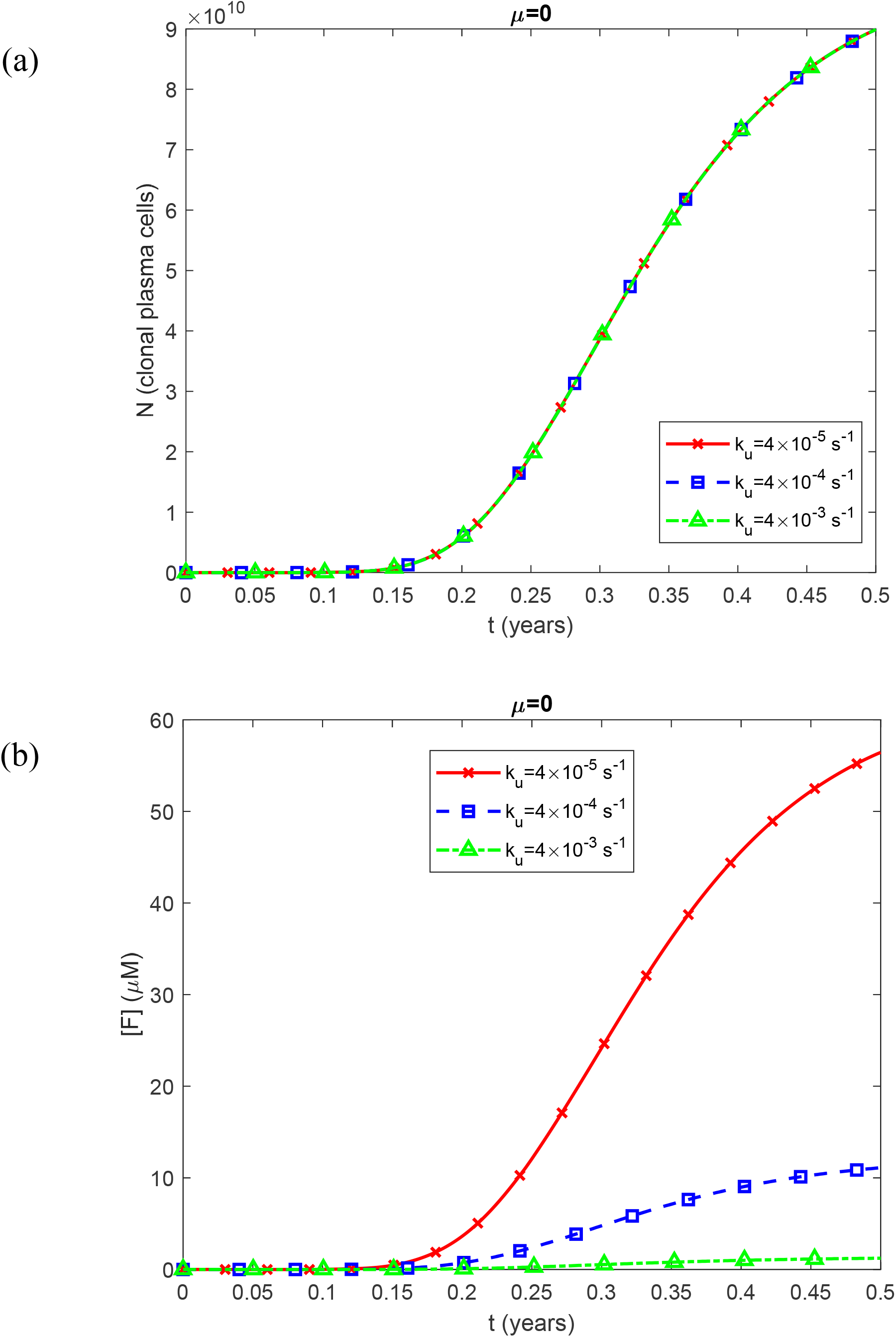
(a) Number of clonal plasma cells, *N*, vs time, *t*. The results indicate that *N* is independent of *k*_*u*_ . (b) Plasma molar concentration of natively folded Ig LC monomers secreted by clonal plasma cells, [*F*], vs time, *t*, for three different values of the rate of unfolding of LCs, *k*_*u*_ . Computations were performed fo 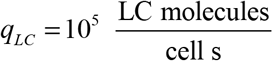. Results correspond to simulations without anticancer therapy.

**Fig. S11.**
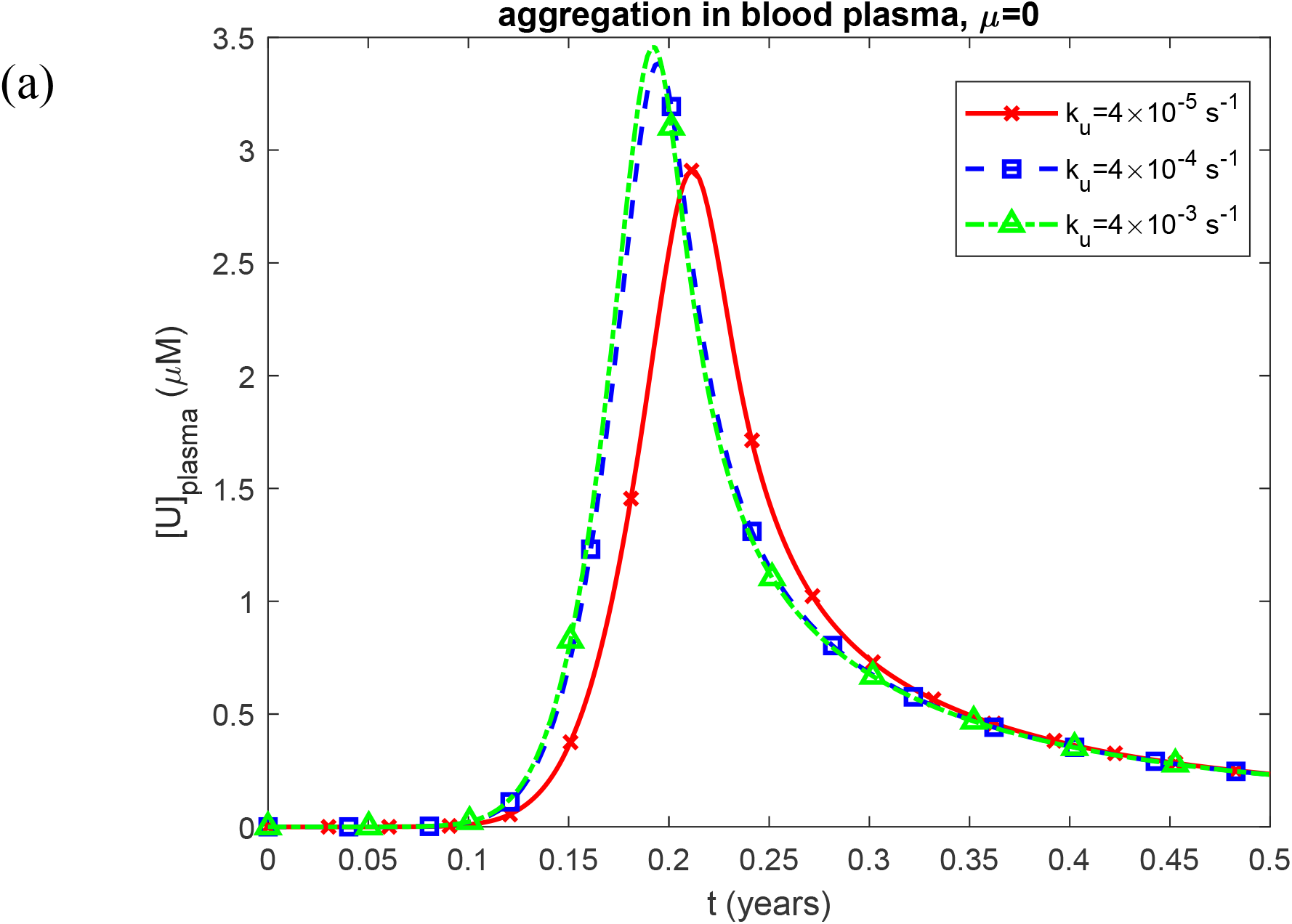
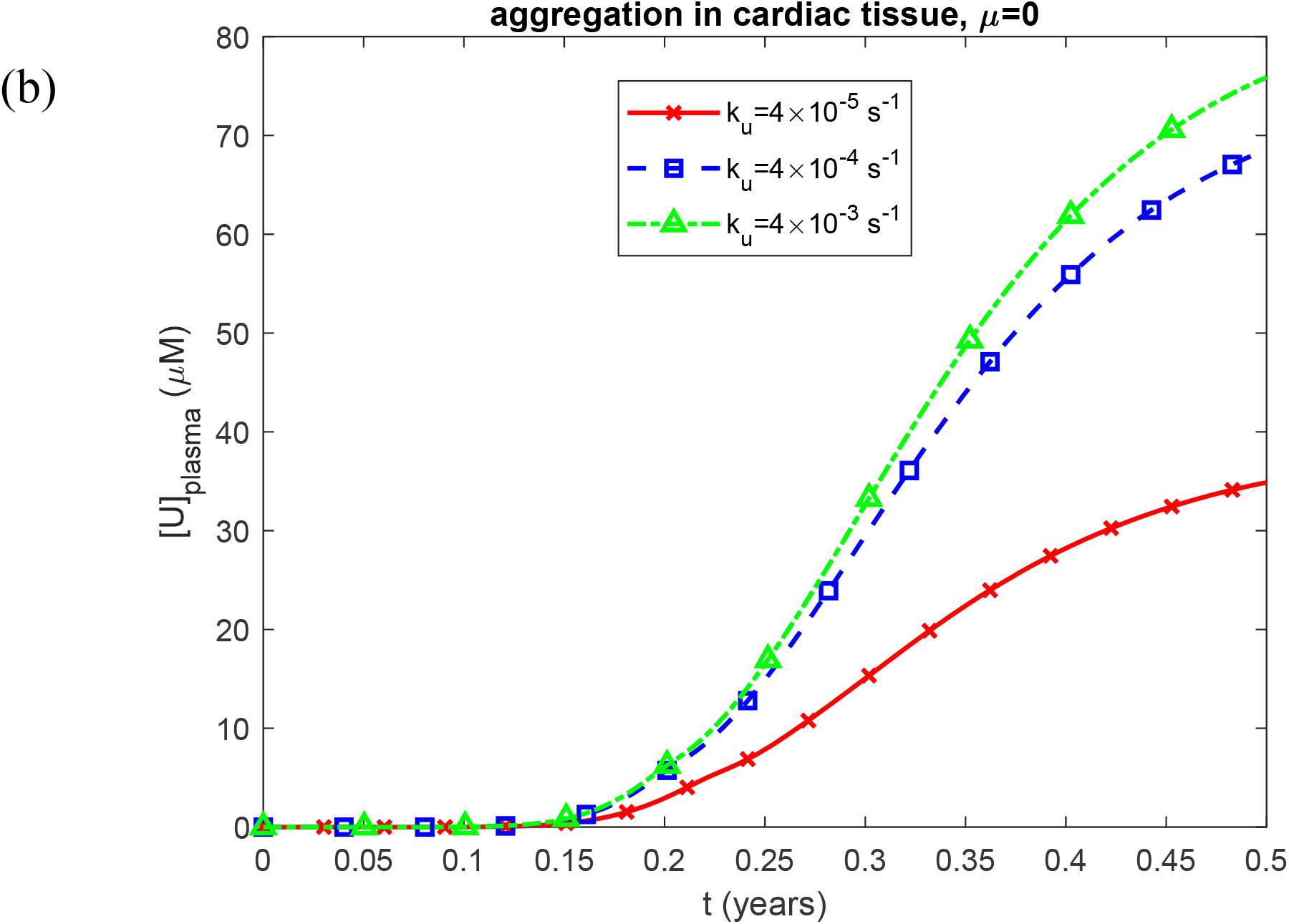
Plasma molar concentration of unfolded Ig LC monomers susceptible to aggregation, [*U*]_*plasma*_, vs time, *t*, for the scenarios of (a) aggregation occurring in the blood plasma and (b) aggregation occurring within the cardiac tissue. Results are shown for three different values of the rate of unfolding of LCs, *k* . Computations were performed for *k* =10^−6^ s^-1^, *k* = 10^−6^ *μ*M^-1^ s^-1^, 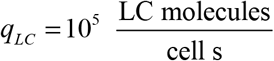, *θ*_1/ 2, *B*_ = 10^7^ s, and *θ*_1/ 2, *I*_ = 10^7^ s. Results correspond to simulations without anticancer therapy.

**Fig. S12.**
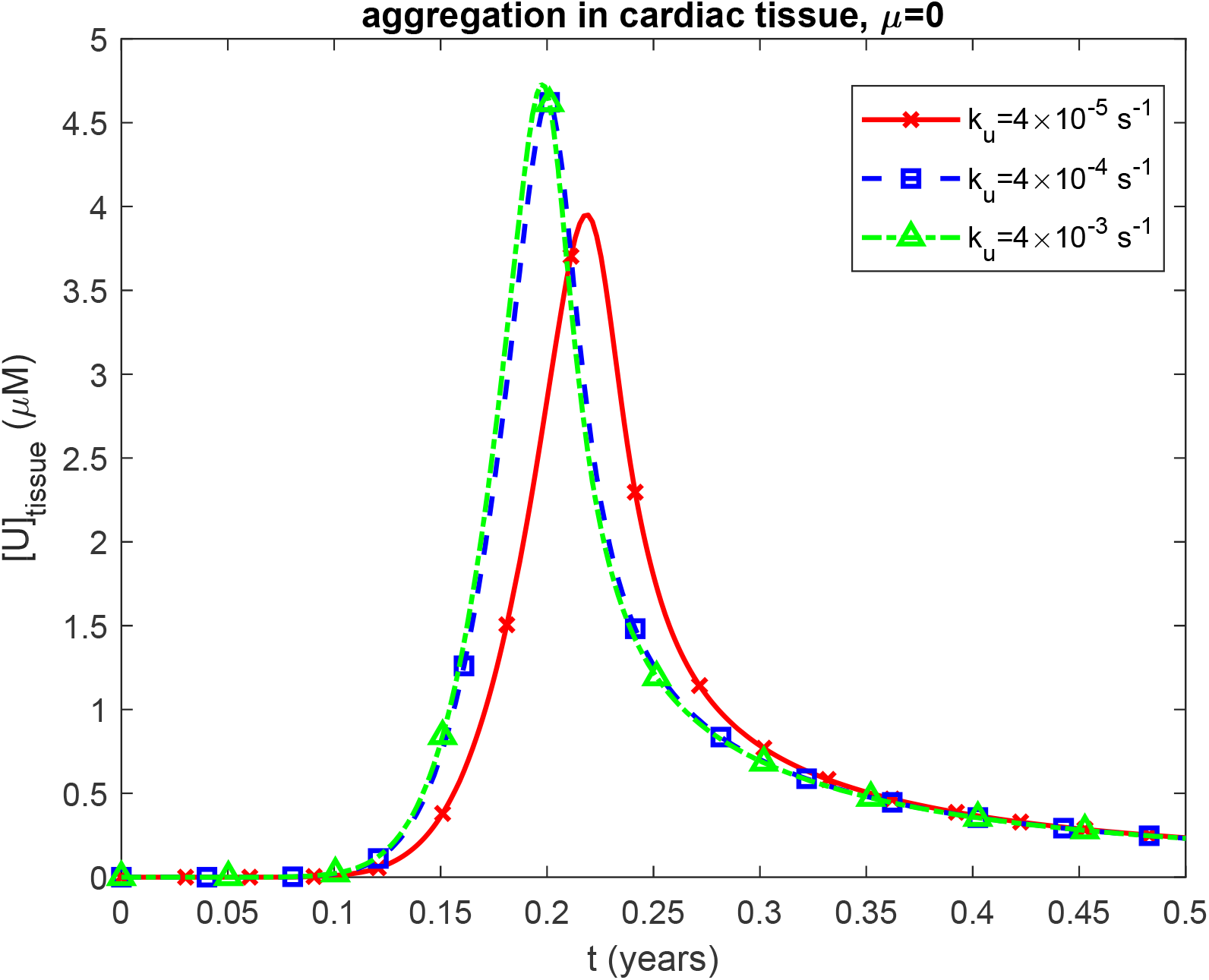
Molar concentration of unfolded Ig LC monomers in cardiac tissue, [*U*]_*tissue*_, vs time, *t*, for the scenario where aggregation occurs within the tissue. Results are shown for three different values of the rate of unfolding of LCs, *k*_*u*_ . Computations were performed for *k*_1_ =10^−6^ s^-1^, *k*_2_ = 10^−6^ *μ*M^-1^ s^-1^, 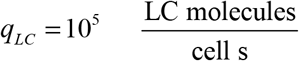, and *θ*_1/ 2, *B*_ = 10^7^ s. Results correspond to simulations without anticancer therapy.

**Fig. S13.**
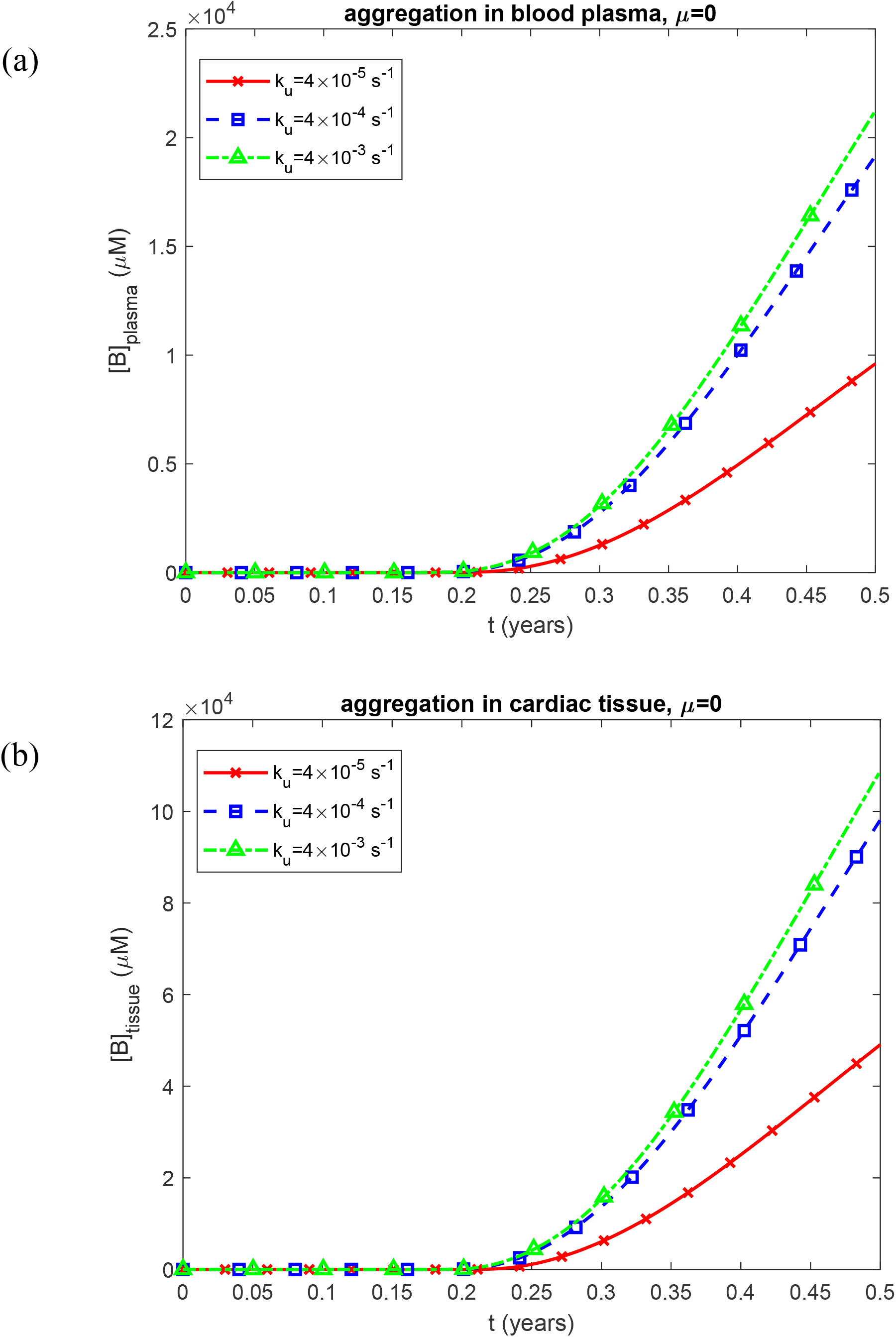
(a) Molar concentration of free LC oligomers in blood plasma, [*B*]_*plasma*_, vs time, *t*, when aggregation occurs in the plasma and (b) molar concentration of free LC oligomers in cardiac tissue, [*B*]_*tissue*_, vs time, *t*, when aggregation occurs within the tissue. Results are shown for three different values of the rate of unfolding of LCs, *k*_*u*_ . Computations were performed for *k*_1_ =10^−6^ s^-1^, *k*_2_ = 10^−6^ *μ*M^-1^ s^-1^, 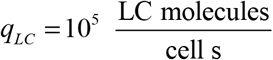, *θ* _1/ 2, *B*_ = 10^7^ s, and *θ*_1/ 2, *I*_ = 10^7^ s. Results correspond to simulations without anticancer therapy.

**Fig. S14.**
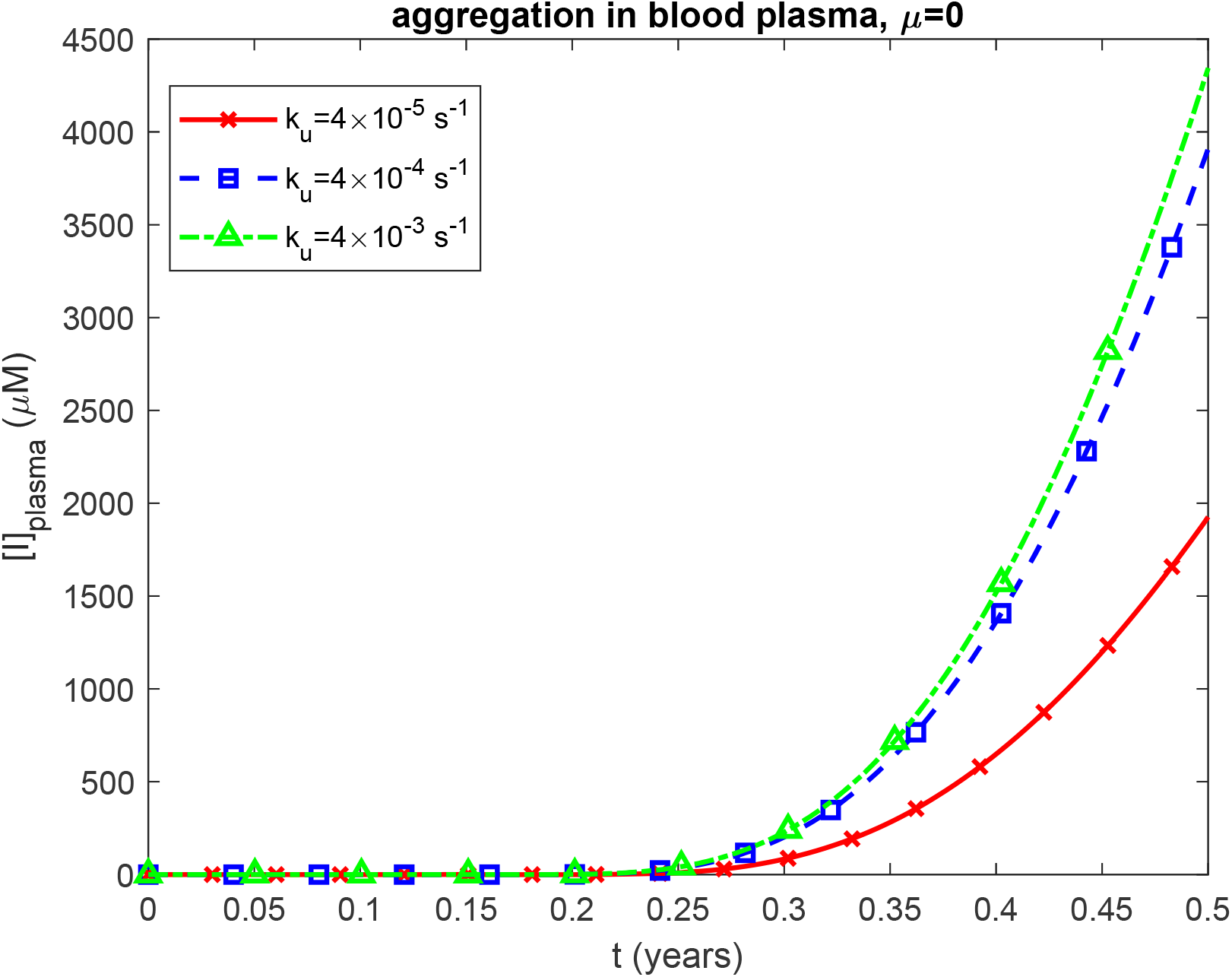
Molar concentration of LC oligomers deposited into amyloid protofibrils circulating within blood plasma, [*I*]_*plasma*_, vs time, *t*. Results are shown for three different values of the rate of unfolding of LCs, *k*_*u*_ . Computations were performed for *k*_1_ =10^−6^ s^-1^, *k*_2_ = 10^−6^ *μ*M^-1^ s^-1^, 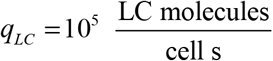, *θ*_1/ 2, *B*_ = 10^7^ s, and *θ* _1/ 2, *BI*_= 10^7^ s. Results correspond to simulations without anticancer therapy.

**Fig. S15.**
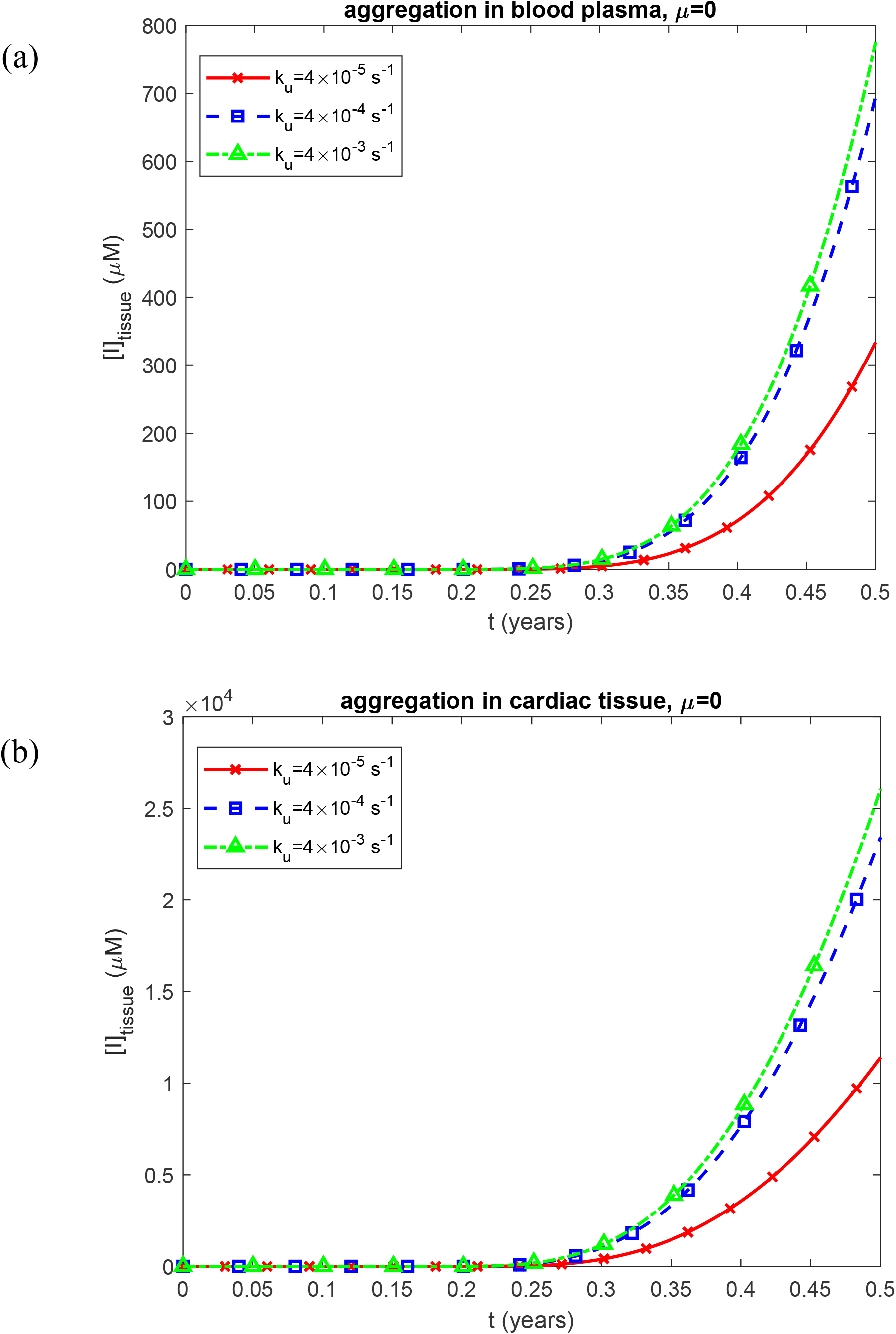
Molar concentration of LC oligomers incorporated into amyloid fibrils infiltrating the cardiac tissue, [*I*]_*tissue*_, vs time, *t*, for two scenarios: (a) aggregation occurring in the blood plasma and (b) aggregation occurring within the cardiac tissue. Results are shown for three different values of the rate of unfolding of LCs, *k*_*u*_ . Computations were performed for *k*_1_ =10^−6^ s^-1^, *k*_2_ = 10^−6^ *μ*M^-1^ s^-1^, 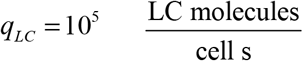, *θ*_1/ 2, *B*_ = 10^7^ s, and *θ*_1/ 2, *I*_ = 10^7^ s. Results correspond to simulations without anticancer therapy.

**Fig. S16.**
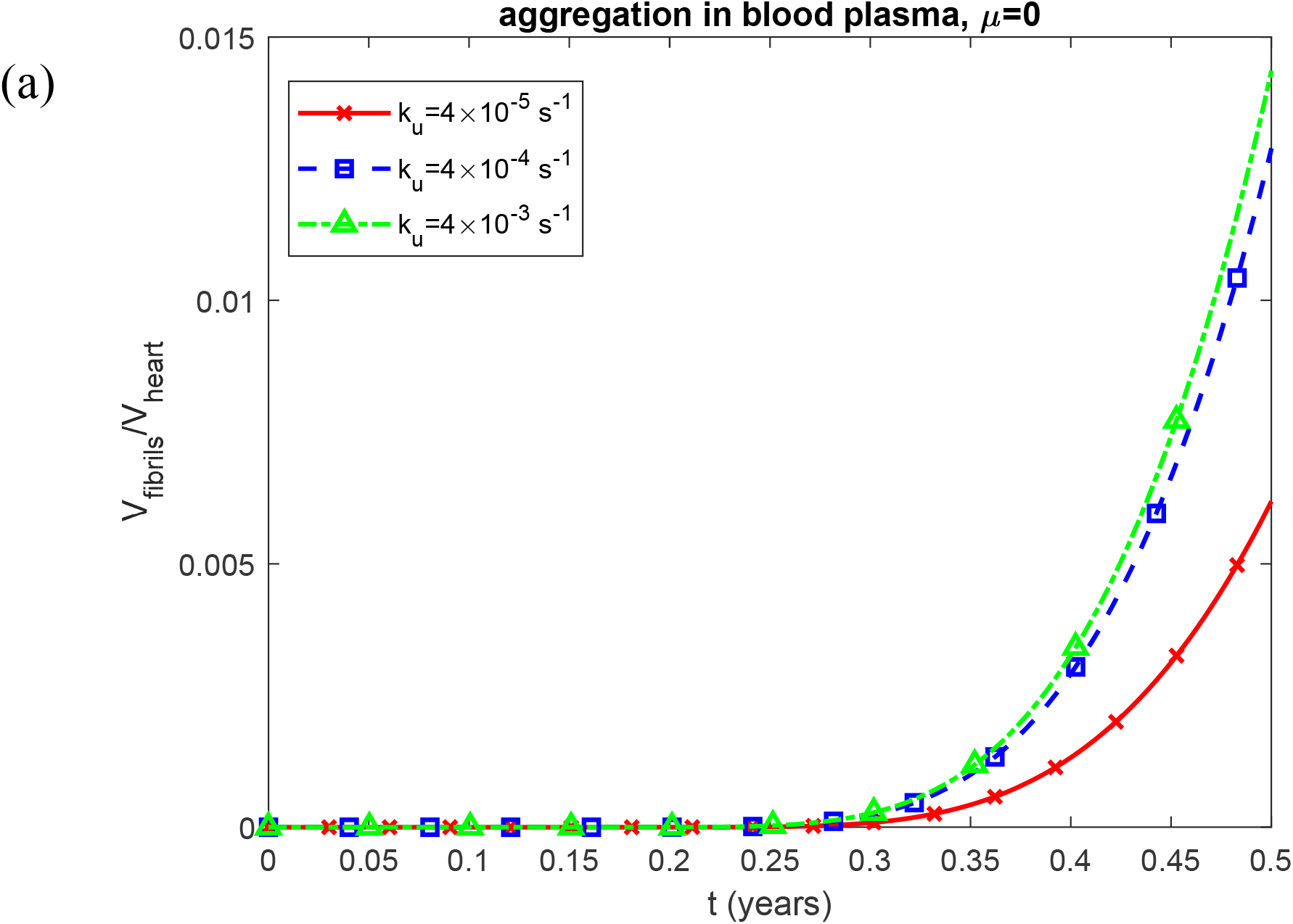
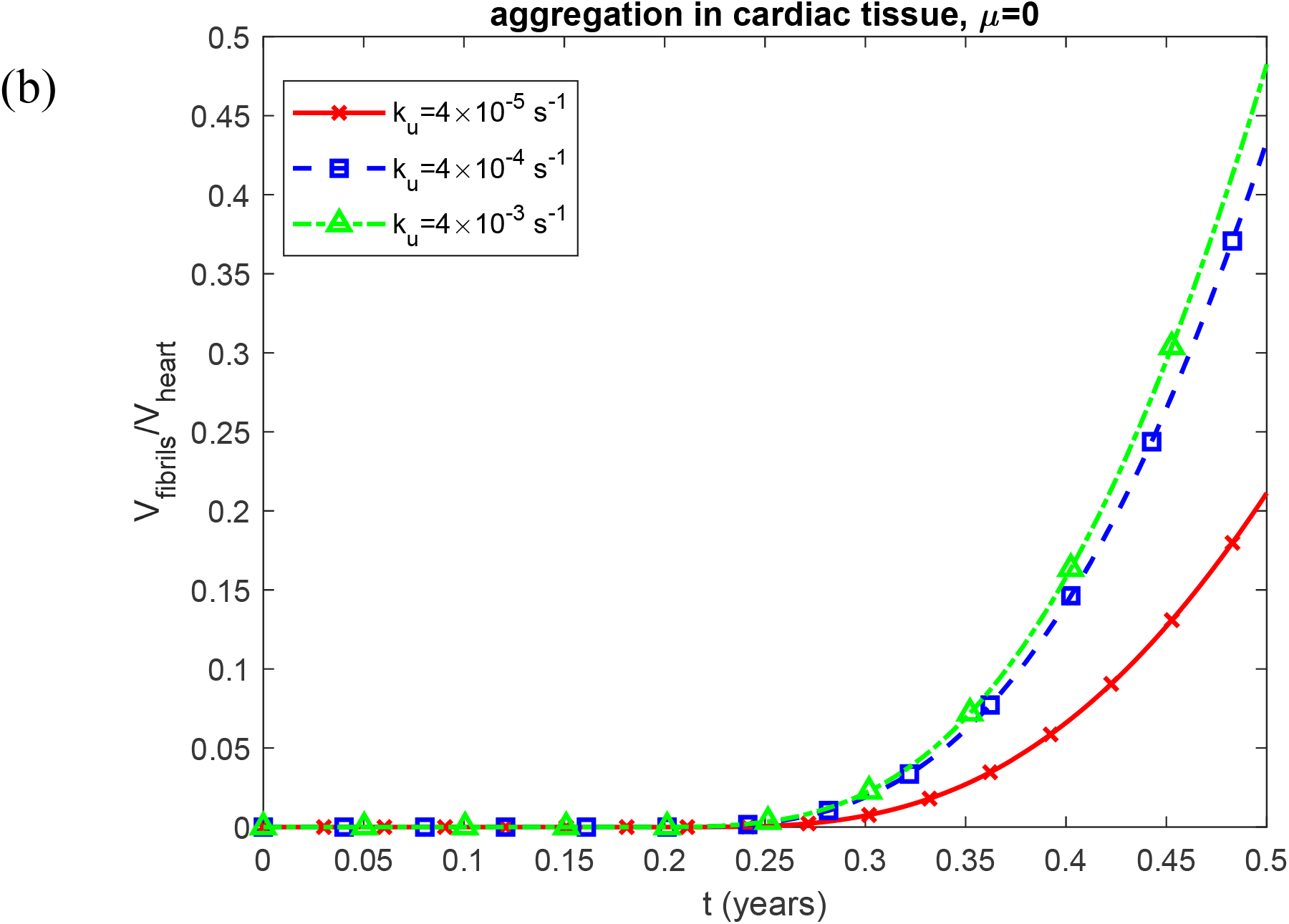
Fraction of myocardial volume occupied by LC fibrils, *V*_*fibrils*_ / *V*_*heart*_, vs time, *t*, for two scenarios: aggregation occurring in the blood plasma and (b) aggregation occurring within the cardiac tissue. Results are shown for three different values of the rate of unfolding of LCs, *k*_*u*_ . Computations were performed for *k*_1_ =10^−6^ s^-1^, *k*_2_ = 10^−6^ *μ*M^-1^ s^-1^, 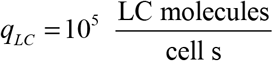, *θ*_1/ 2, *B*_ = 10^7^ s, and *θ*_1/ 2, *I*_ = 10^7^ s. Results correspond to simulations without anticancer therapy.

**Fig. S17.**
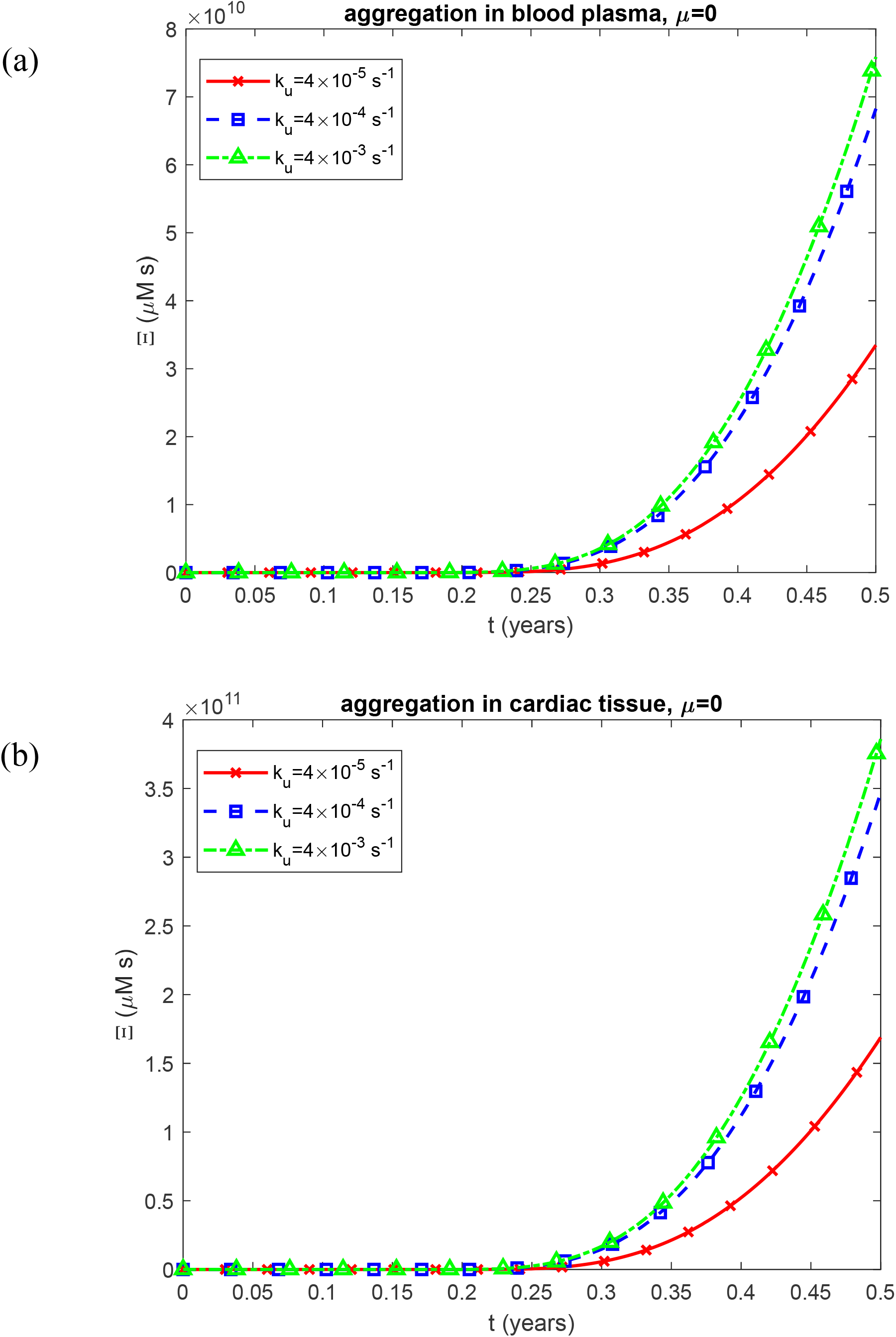
Time-dependent criterion for accumulated cardiotoxicity induced by LC oligomers, Ξ, vs time, *t*, for two scenarios: (a) aggregation occurring in the blood plasma and (b) aggregation occurring within the cardiac tissue. Results are shown for three different values of the rate of unfolding of LCs, *k*_*u*_ . Computations were performed for *k*_1_ =10^−6^ s^-1^, *k*_2_ = 10^−6^ *μ*M^-1^ s^-1^, 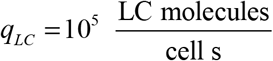, *θ* _1/ 2, *B*_ = 10^7^ s, and *θ*_1/ 2, *I*_ = 10^7^ s. Results correspond to simulations without anticancer therapy.

**Fig. S18.**
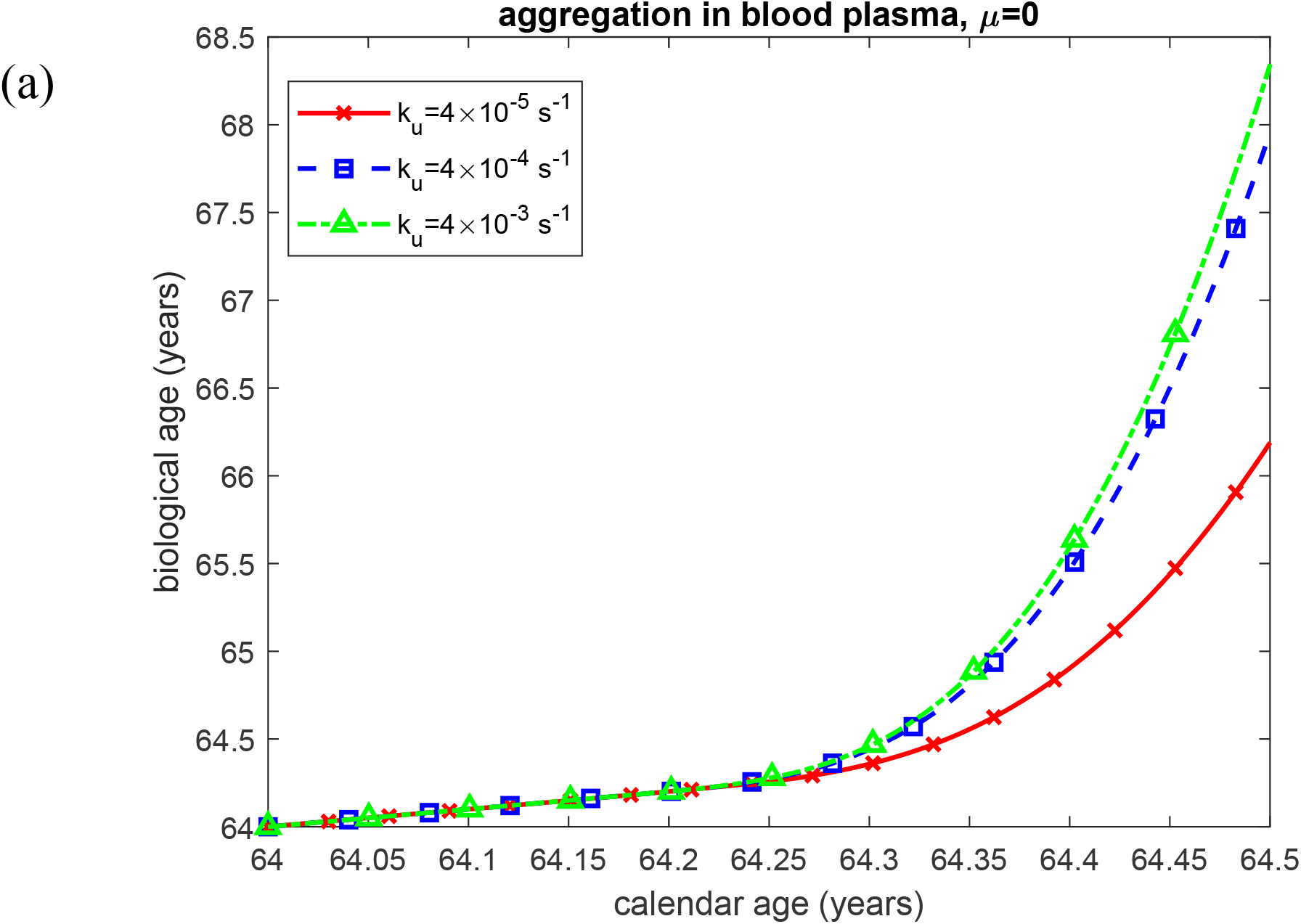
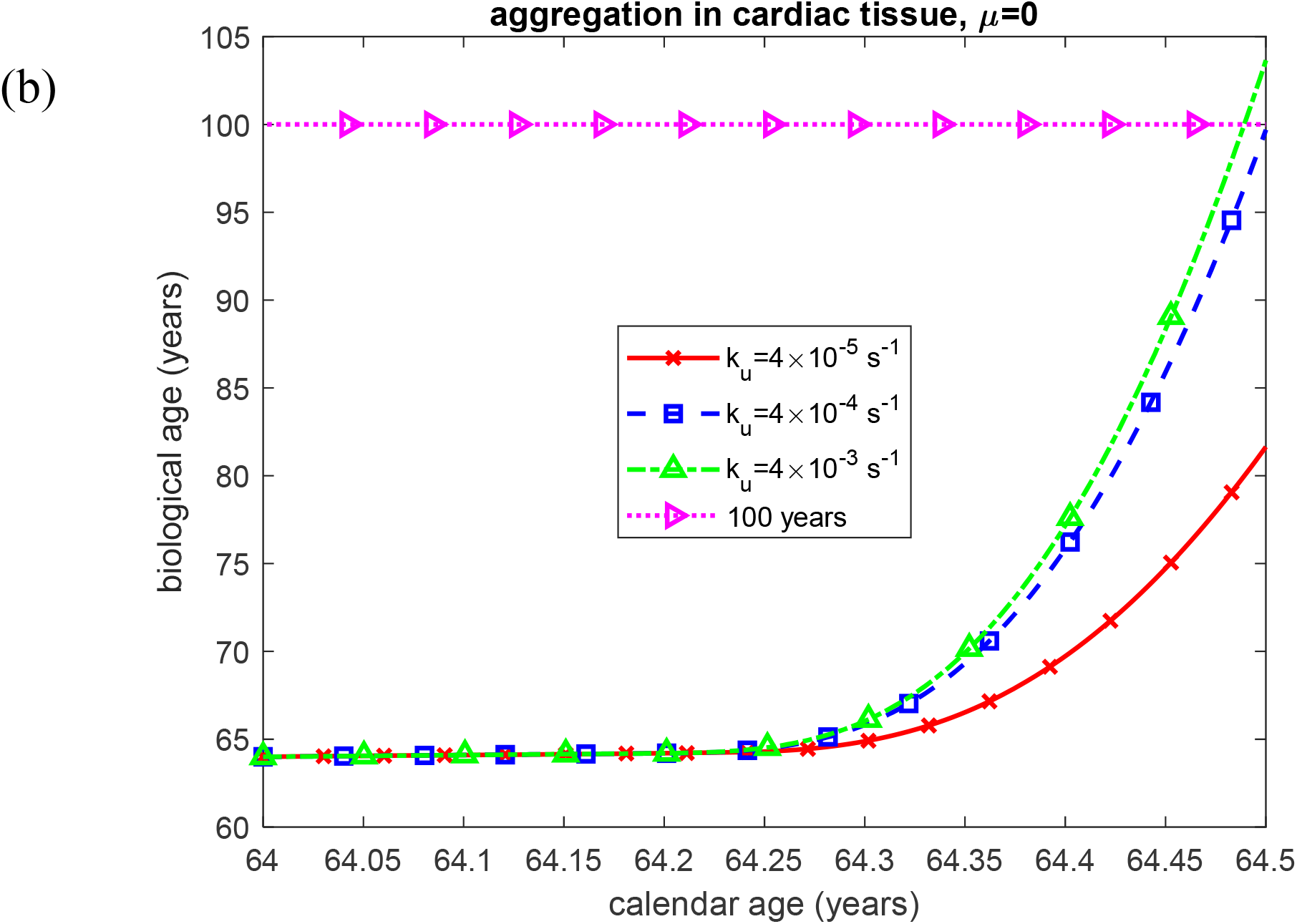
Biological age vs calendar age under two scenarios: (a) aggregation occurring in the blood plasma and (b) aggregation occurring within the cardiac tissue. Results are shown for three different values of the rate of unfolding of LCs, *k*_*u*_ . Computations were performed for *k*_1_ =10^−6^ s^-1^, *k*_2_ = 10^−6^ *μ*M^-1^ s^-1^, 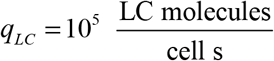, *θ*_1/ 2, *B*_ = 10^7^ s, and *θ*_1/ 2, *I*_ = 10^7^ s. Results correspond to simulations without anticancer therapy.

**Fig. S19.**
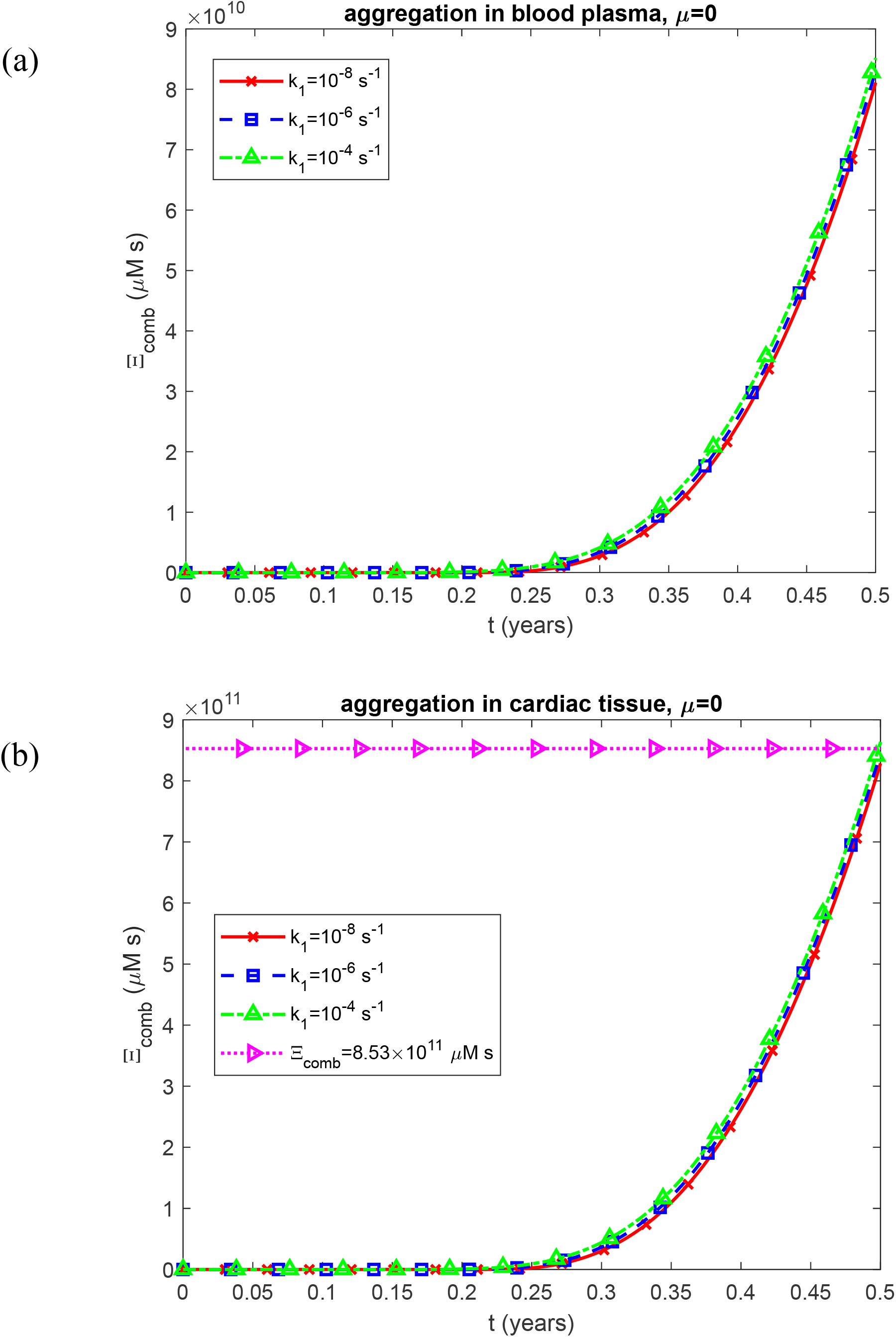
Time-dependent criterion characterizing the combined effect of cytotoxicity and myocardial stiffening, Ξ _*comb*_, vs time, *t*, for two scenarios: (a) aggregation occurring in the blood plasma and (b) aggregation occurring within the cardiac tissue. Results are shown for three different values of the rate constant describing the first pseudoelementary (nucleation) step in the F–W model of LC oligomer formation, *k*_1_ . Computations were performed for *k*_2_ = 10^−6^ *μ*M^-1^ s^-1^, *k*_*u*_ = 4 ×10^−4^ s^-1^, 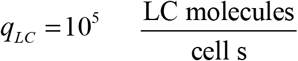, *θ*_1/ 2, *B*_ = 10^7^ s, and *θ* _1/ 2, *I*_ = 10^7^ s. Results correspond to simulations without anticancer therapy.

**Fig. S20.**
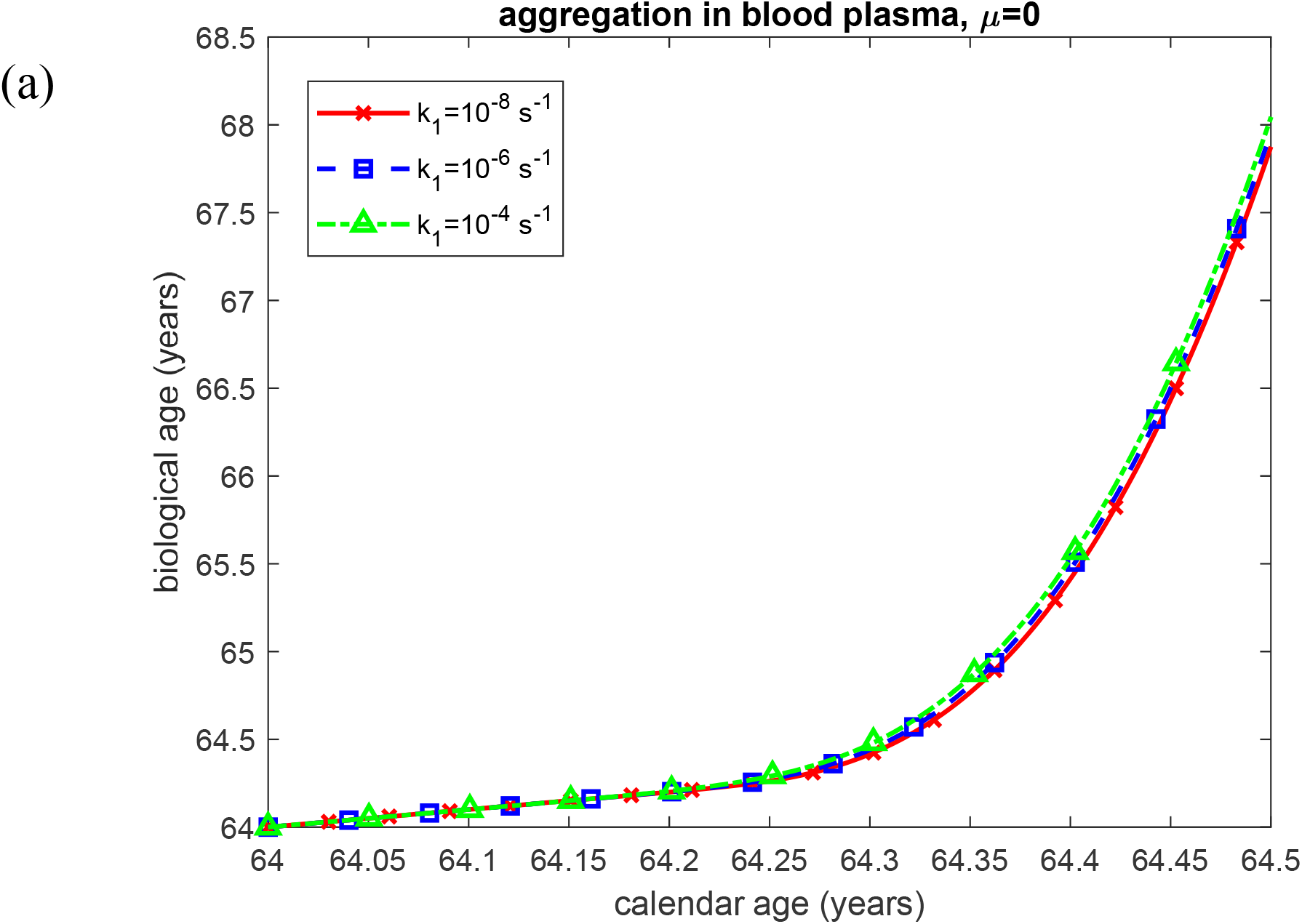
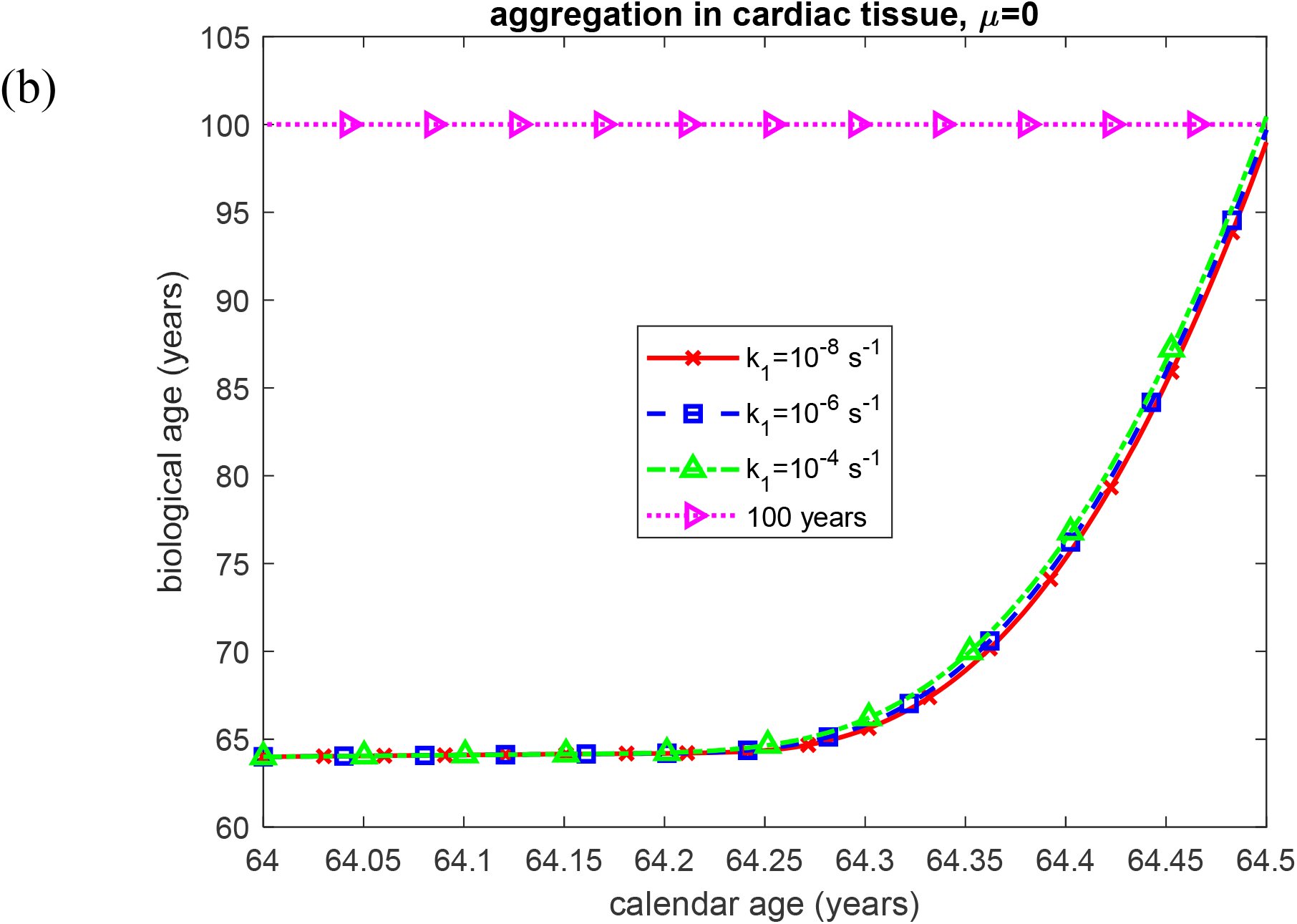
Biological age vs calendar age under two scenarios: (a) aggregation occurring in the blood plasma and (b) aggregation occurring within the cardiac tissue. Results are shown for three different values of the rate constant describing the first pseudoelementary (nucleation) step in the F–W model of LC oligomer formation, *k*_1_ . Computations were performed for *k*_2_ = 10^−6^ *μ*M^-1^ s^-1^, *k*_*u*_ = 4 ×10^−4^ s^-1^, 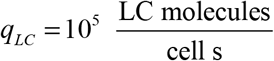, *θ*_1/ 2, *B*_ = 10^7^ s, and *θ*_1/ 2, *I*_ = 10^7^ s. Results correspond to simulations without anticancer therapy.

**Fig. S21.**
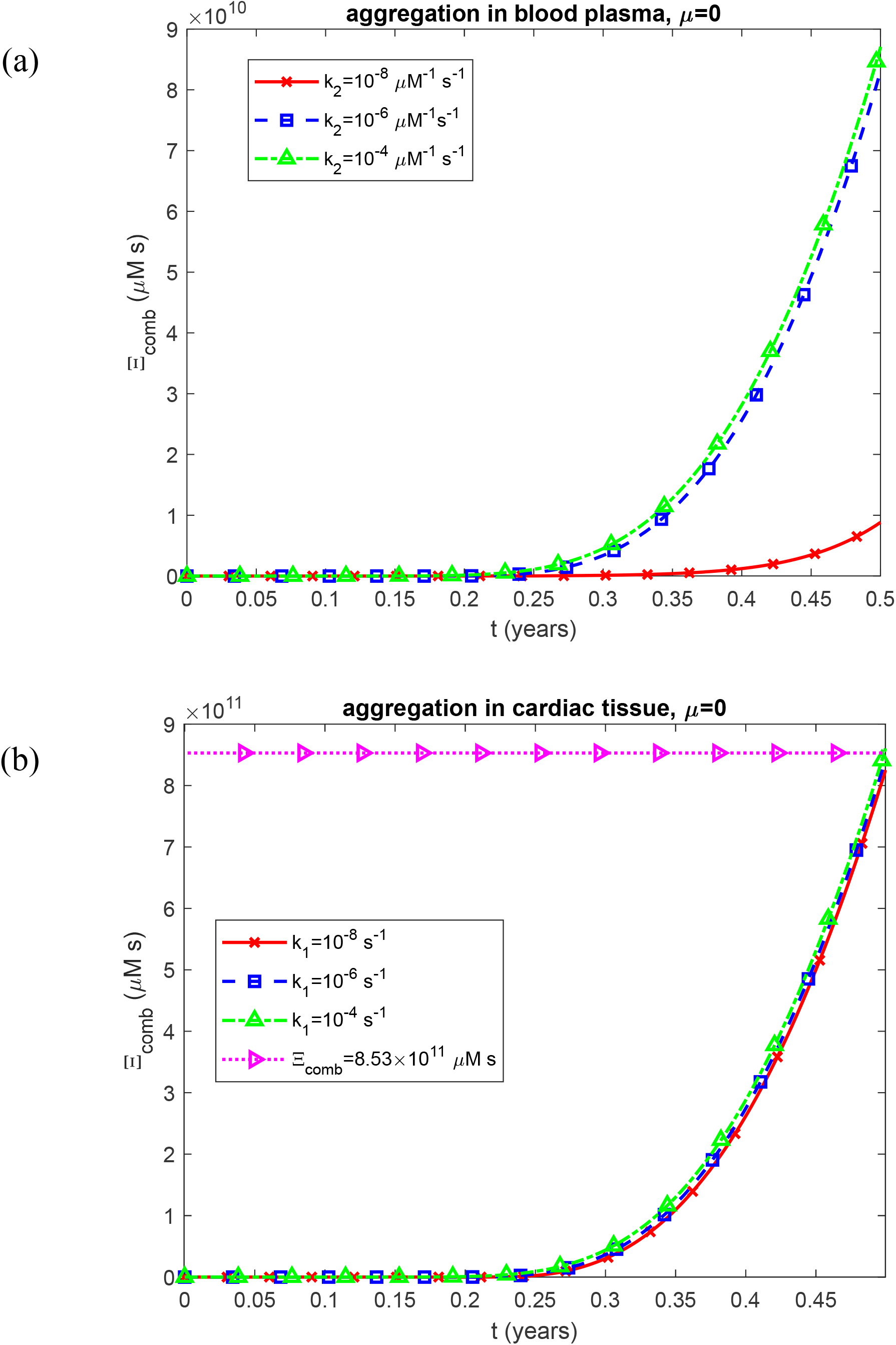
Time-dependent criterion characterizing the combined effect of cytotoxicity and myocardial stiffening, Ξ _*comb*_, vs time, *t*, for two scenarios: (a) aggregation occurring in the blood plasma and (b) aggregation occurring within the cardiac tissue. Results are shown for three different values of the rate constant describing the second pseudoelementary (autocatalytic growth) step in the F–W model of LC oligomer formation, *k*_2_ . Computations were performed for *k*_1_ =10^−6^ s^-1^, *k*_*u*_ = 4 ×10^−4^ s^-1^, 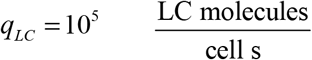, *θ*_1/ 2, *B*_ = 10^7^ s, and *θ*_1/ 2, *I*_ = 10^7^ s. Results correspond to simulations without anticancer therapy.

**Fig. S22.**
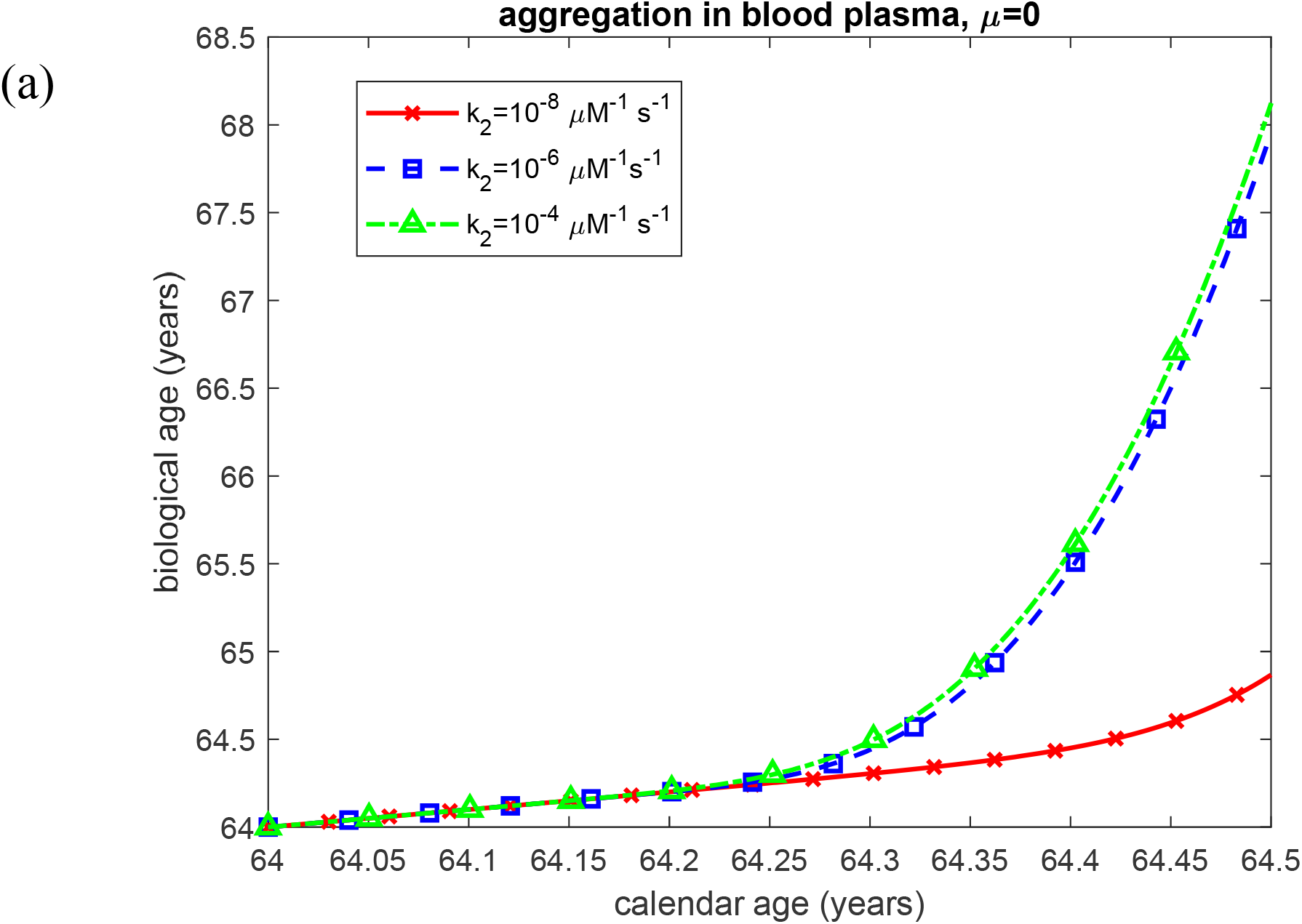
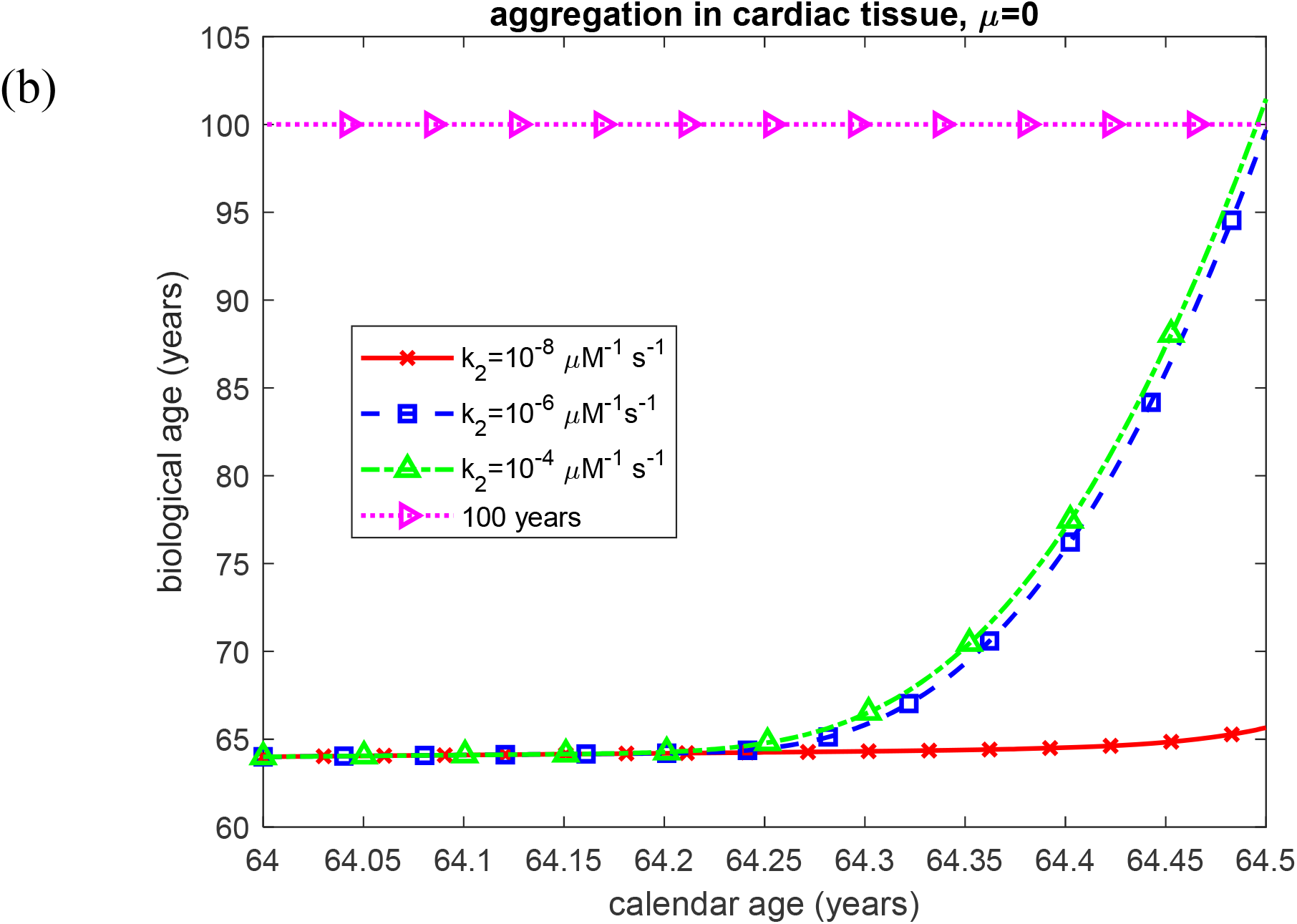
Biological age vs calendar age for two scenarios: (a) aggregation occurring in the blood plasma and (b) aggregation occurring within the cardiac tissue. Results are shown for three different values of the rate constant describing the second pseudoelementary (autocatalytic growth) step in the F–W model of LC oligomer formation, *k*_2_ . Computations were performed for *k*_1_ =10^−6^ s^-1^, *k*_*u*_ = 4 ×10^−4^ s^-1^, 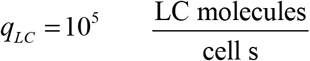, *θ*_1/ 2, *B*_ = 10^7^ s, and *θ*_1/ 2, *I*_ = 10^7^ s. Results correspond to simulations without anticancer therapy.

**Fig. S23.**
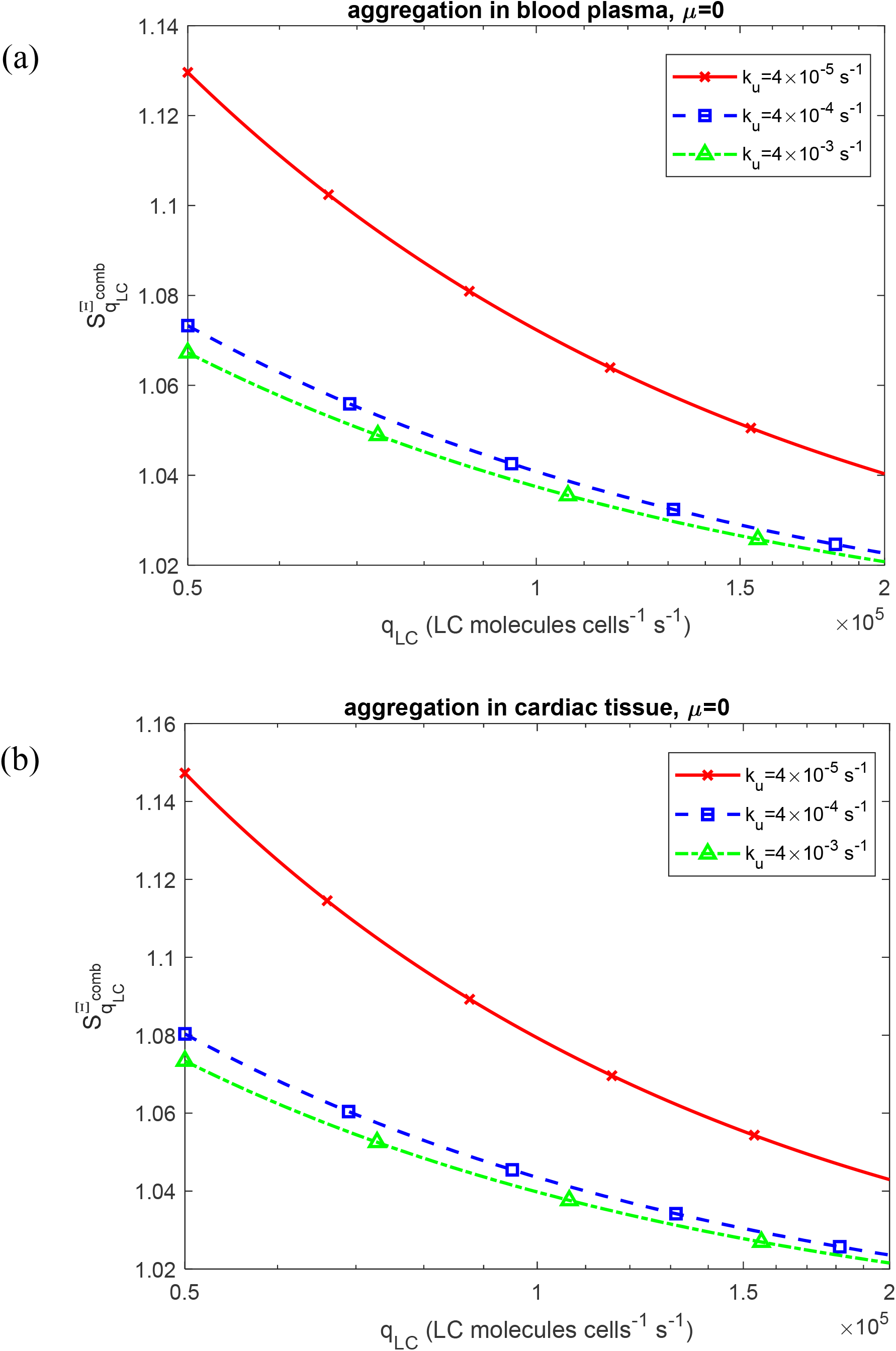
Dimensionless sensitivity of the criterion characterizing the combined effects of cytotoxicity and myocardial stiffening to the rate of secretion of folded LCs by a single plasma cell belonging to a clonal population, 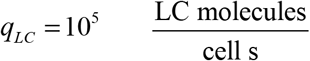, vs *q*_*LC*_ . Results are presented at the end of the six-month terminal cardiac-progression interval for three different values of the LC unfolding rate constant, *k*_*u*_ . Computations were performed for *k*_1_ =10^−6^ s^-1^, *k*_2_ = 10^−6^ *μ*M^-1^ s^-1^, *θ* _1/ 2, *B*_ = 10^7^ s, and *θ* _1/ 2, *I*_ = 10^7^ s. Results correspond to simulations without anticancer therapy.

**Fig. S24.**
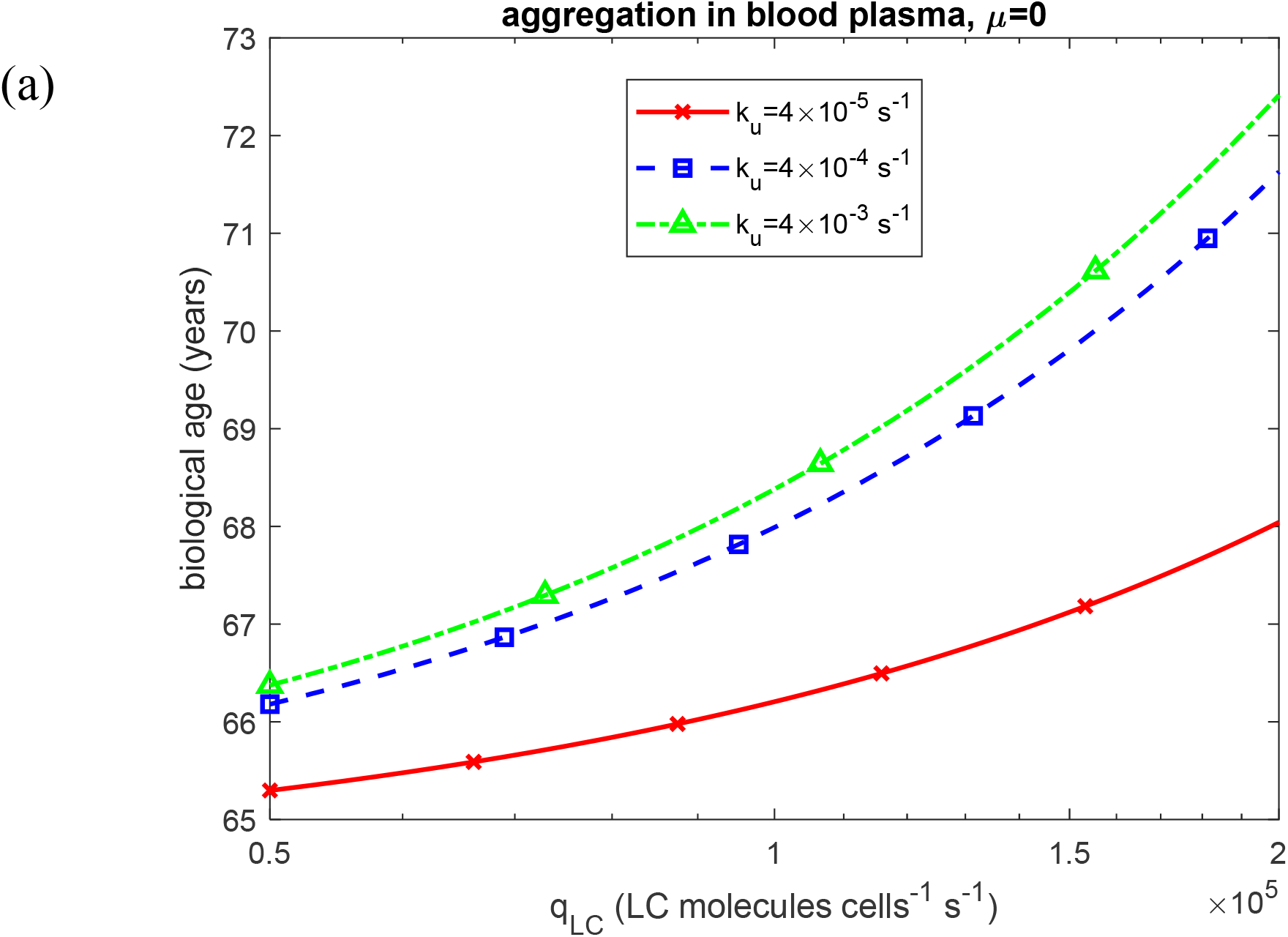
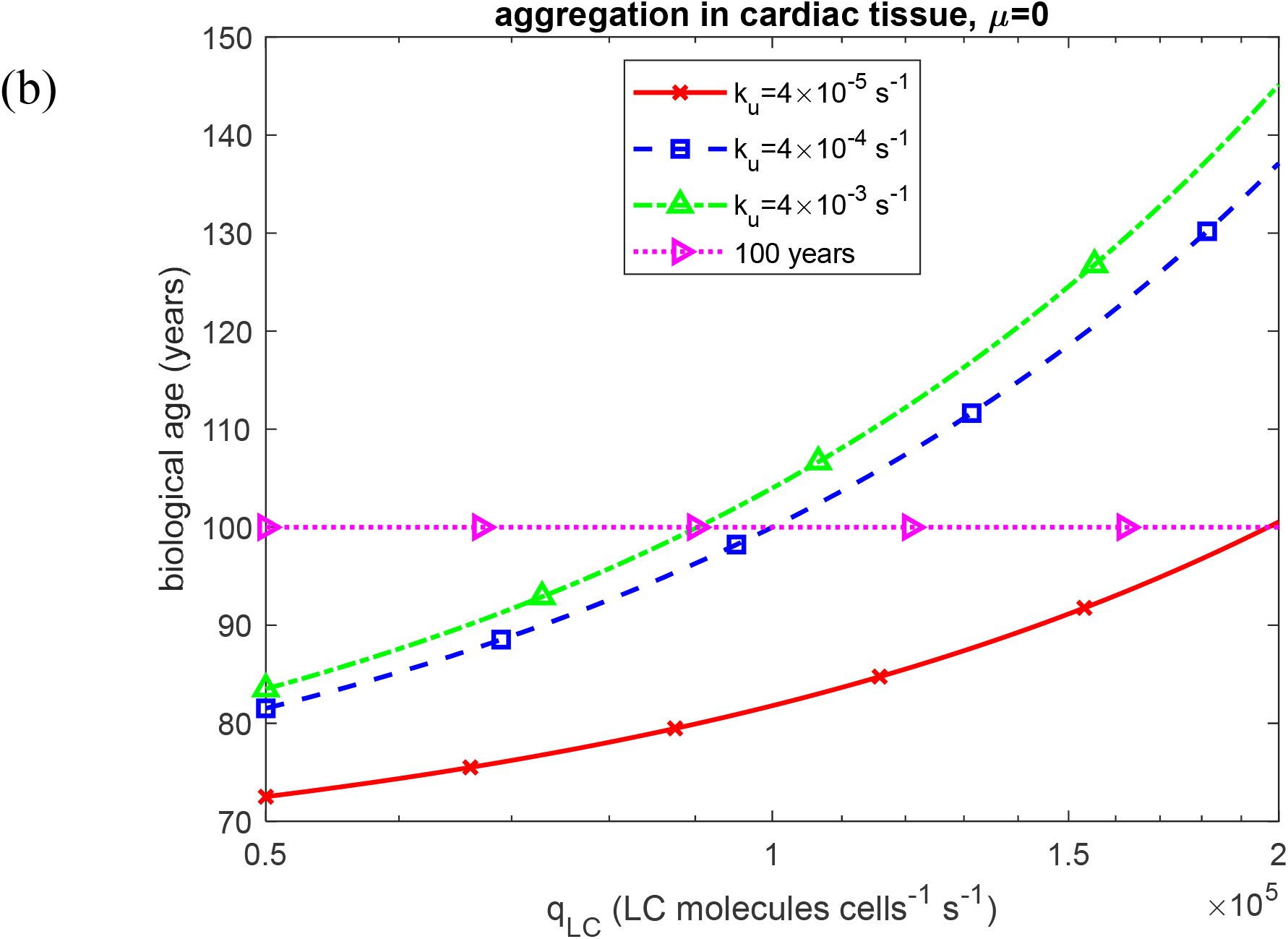
Biological age vs the rate of secretion of folded LCs by a single plasma cell belonging to a clonal population at the end of the six-month modeled terminal cardiac-progression interval for two scenarios: (a) aggregation occurring in the blood plasma and (b) aggregation occurring within the cardiac tissue. Results are shown for three different values of the rate of unfolding of LCs, *k*_*u*_ . Computations were performed for *k*_1_ =10^−6^ s^-1^, *k*_2_ = 10^−6^ *μ*M^-1^ s^-1^, *θ*_1/ 2, *B*_ = 10^7^ s, and *θ*_1/ 2, *I*_ = 10^7^ s. Results correspond to simulations without anticancer therapy.

**Fig. S25.**
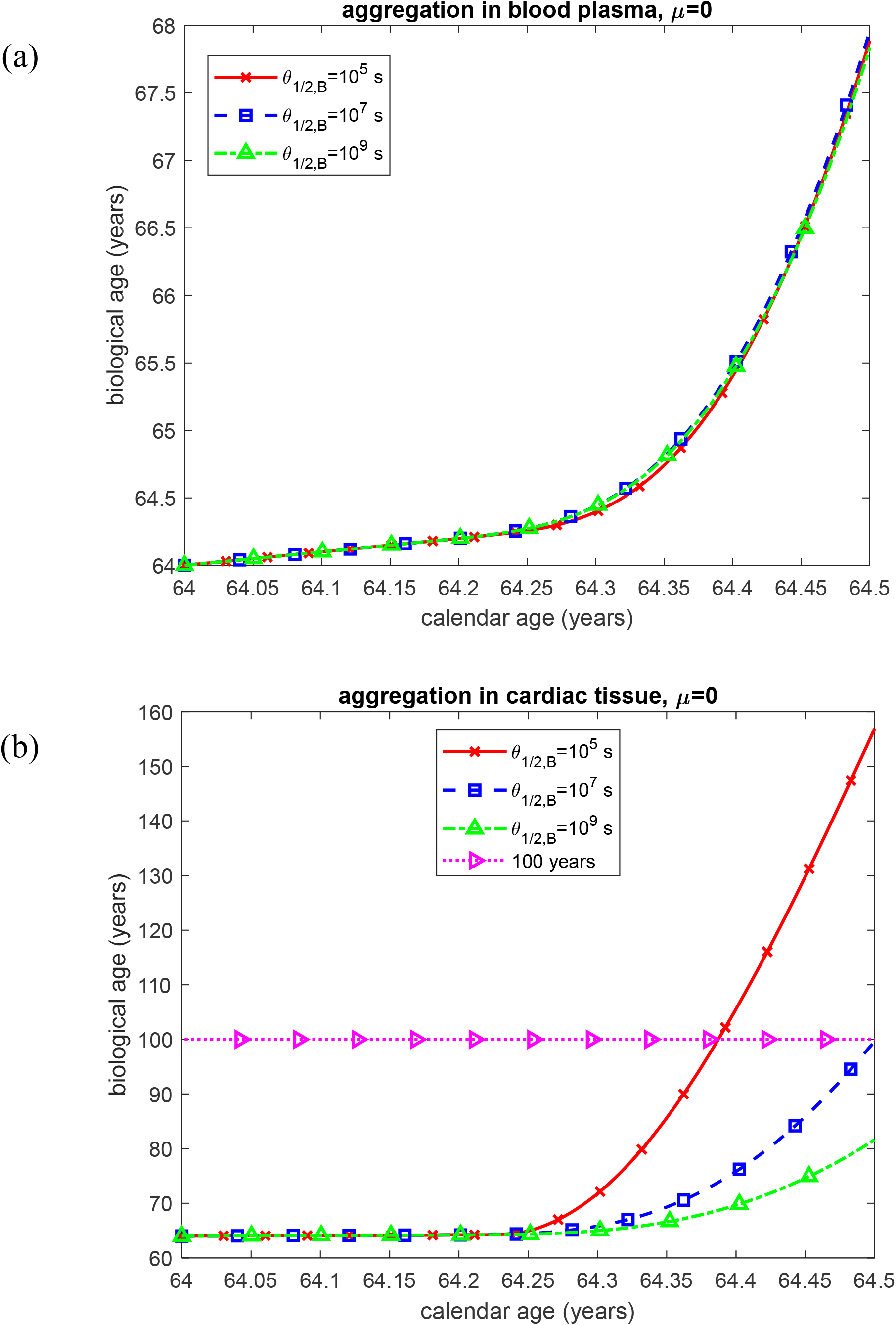
Biological age vs calendar age for two scenarios: (a) aggregation occurring in the blood plasma and (b) aggregation occurring within the cardiac tissue. Results are shown for three different values of the half-deposition time for free LC oligomers to be incorporated into LC fibrils, *θ*_1/2,*B*_ . Computations were performed for *k*_1_ =10^−6^ s^-1^, *k*_2_ = 10^−6^ *μ*M^-1^ s^-1^, *k*_*u*_ = 4 ×10^−4^ s^-1^, 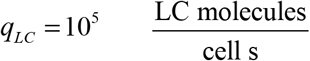, and *θ*_1/ 2, *I*_ = 10^7^ s. Results correspond to simulations without anticancer therapy.

**Fig. S26.**
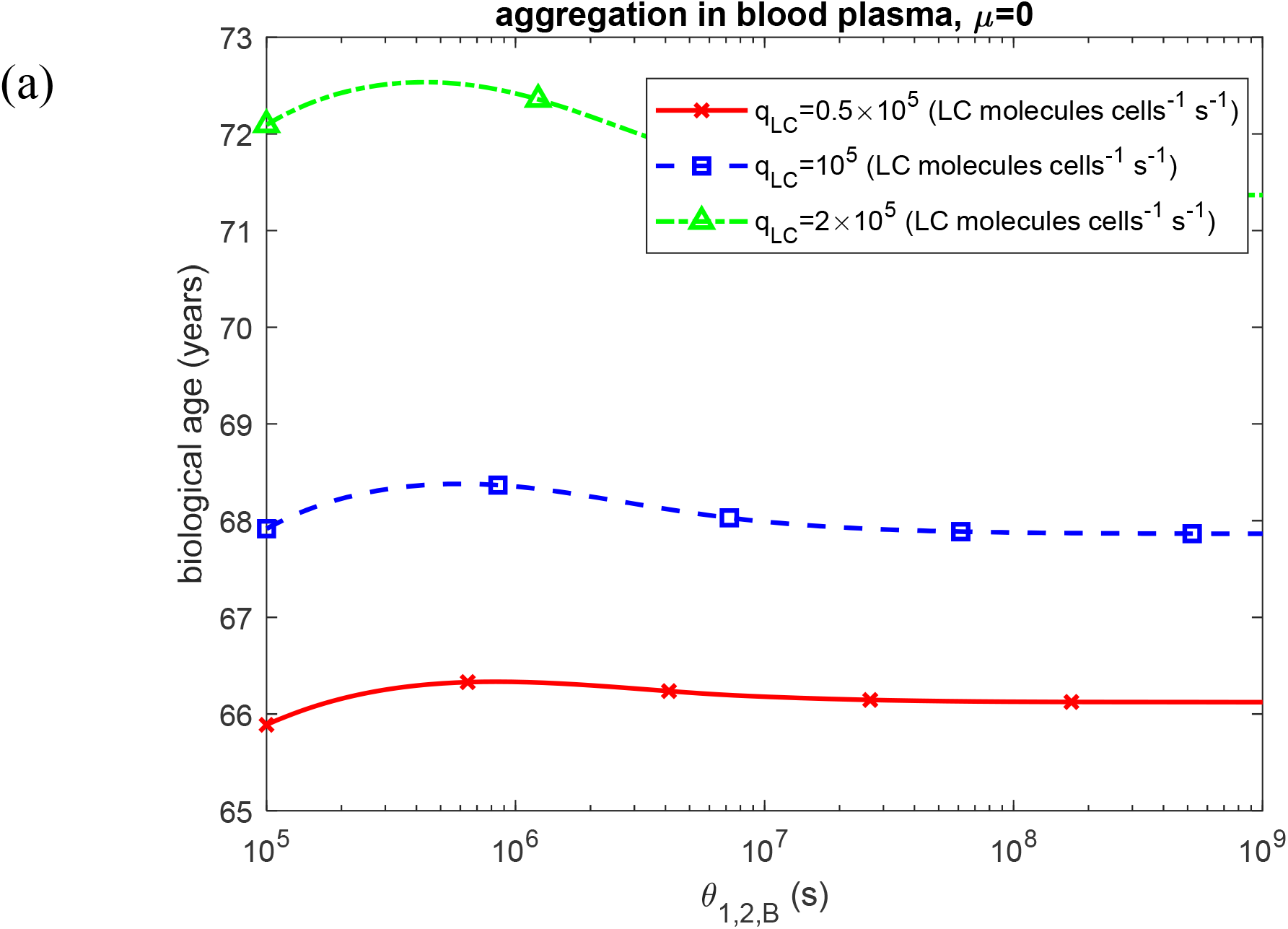
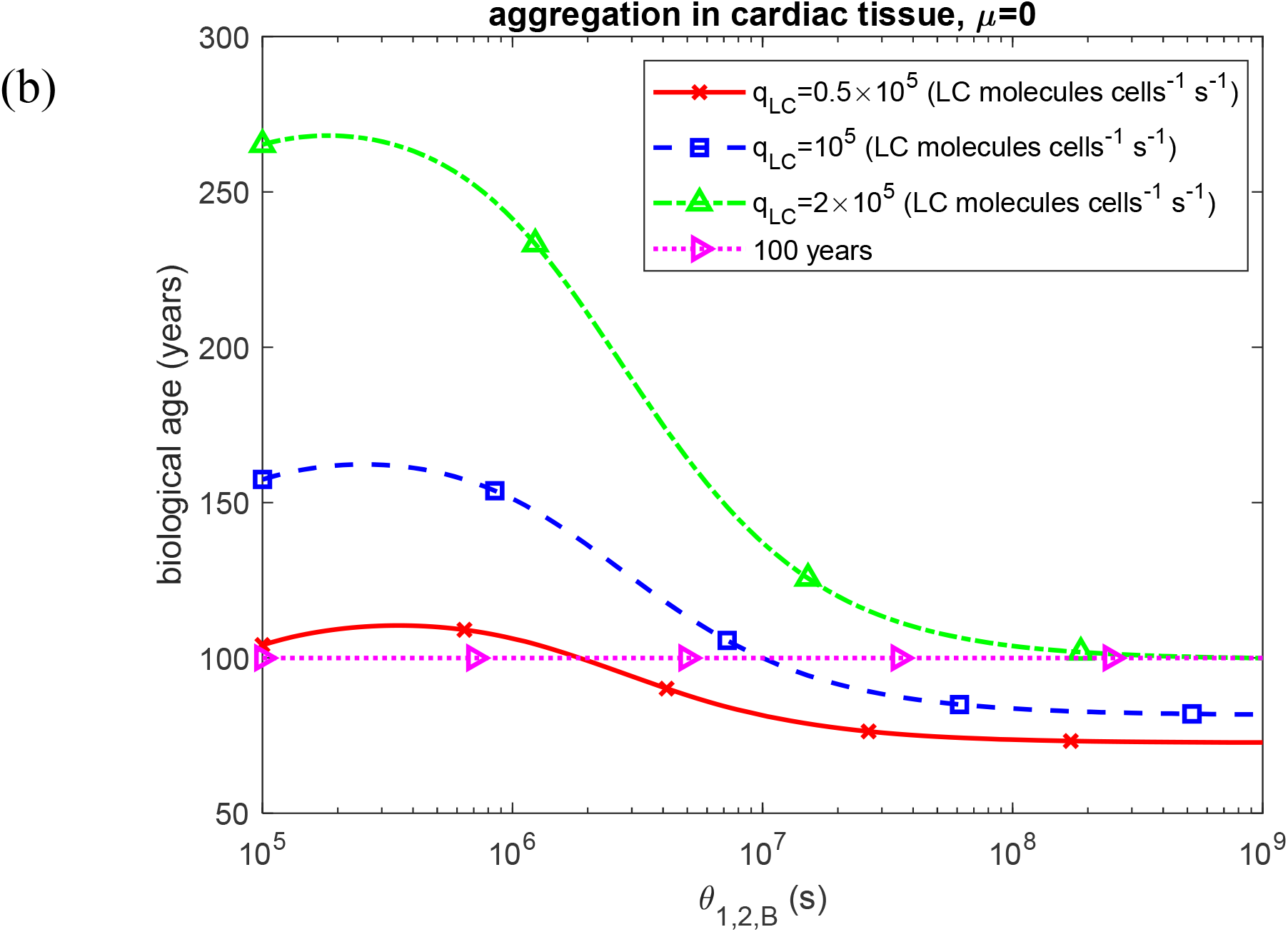
Biological age vs the half-deposition time for free LC oligomers to be incorporated into LC fibrils at the end of the six-month modeled terminal cardiac-progression interval for two scenarios: (a) aggregation occurring in the blood plasma and (b) aggregation occurring within the cardiac tissue. Results are presented for three different values of the LC secretion rate per clonal plasma cell, *q* _*LC*_ . Computations were performed for *k*_1_ =10^−6^ s^-1^, *k*_2_ = 10^−6^ *μ*M^-1^ s^-1^, *k*_*u*_ = 4 ×10^−4^ s^-1^, and *θ*_1/ 2, *I*_ = 10^7^ s. Results correspond to simulations without anticancer therapy.

**Fig. S27.**
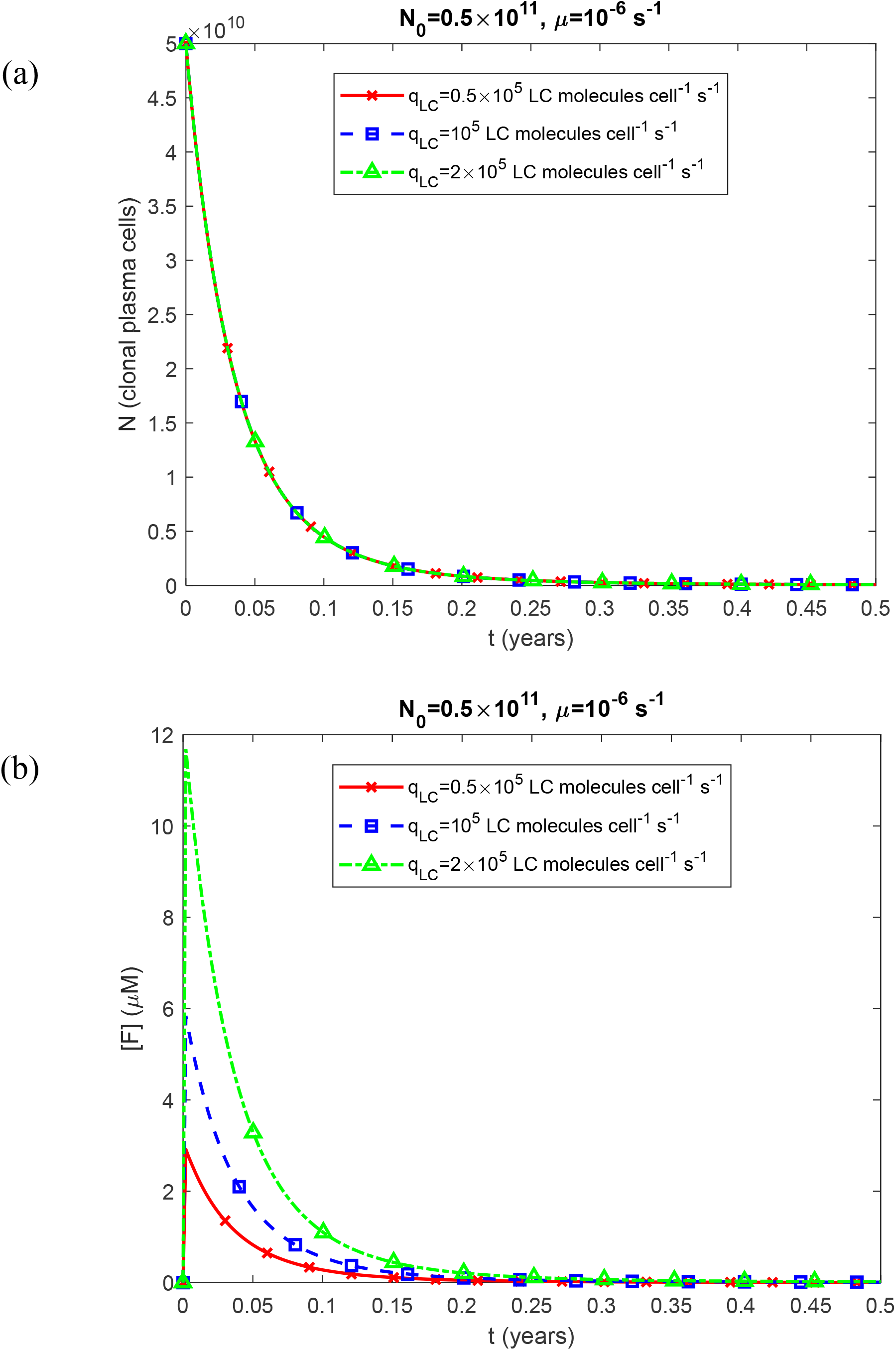
(a) Number of clonal plasma cells, *N*, vs time, *t*. The results indicate that *N* is independent of *q* _*LC*_ . (b) Plasma molar concentration of natively folded Ig LC monomers secreted by clonal plasma cells vs time for three different values of the secretion rate of folded LCs per single plasma cell belonging to a clonal population, *q*_*LC*_ . Computations were performed for *k*_*u*_ = 4 ×10^−4^ s^-1^. Results correspond to simulations with anticancer therapy.

**Fig. S28.**
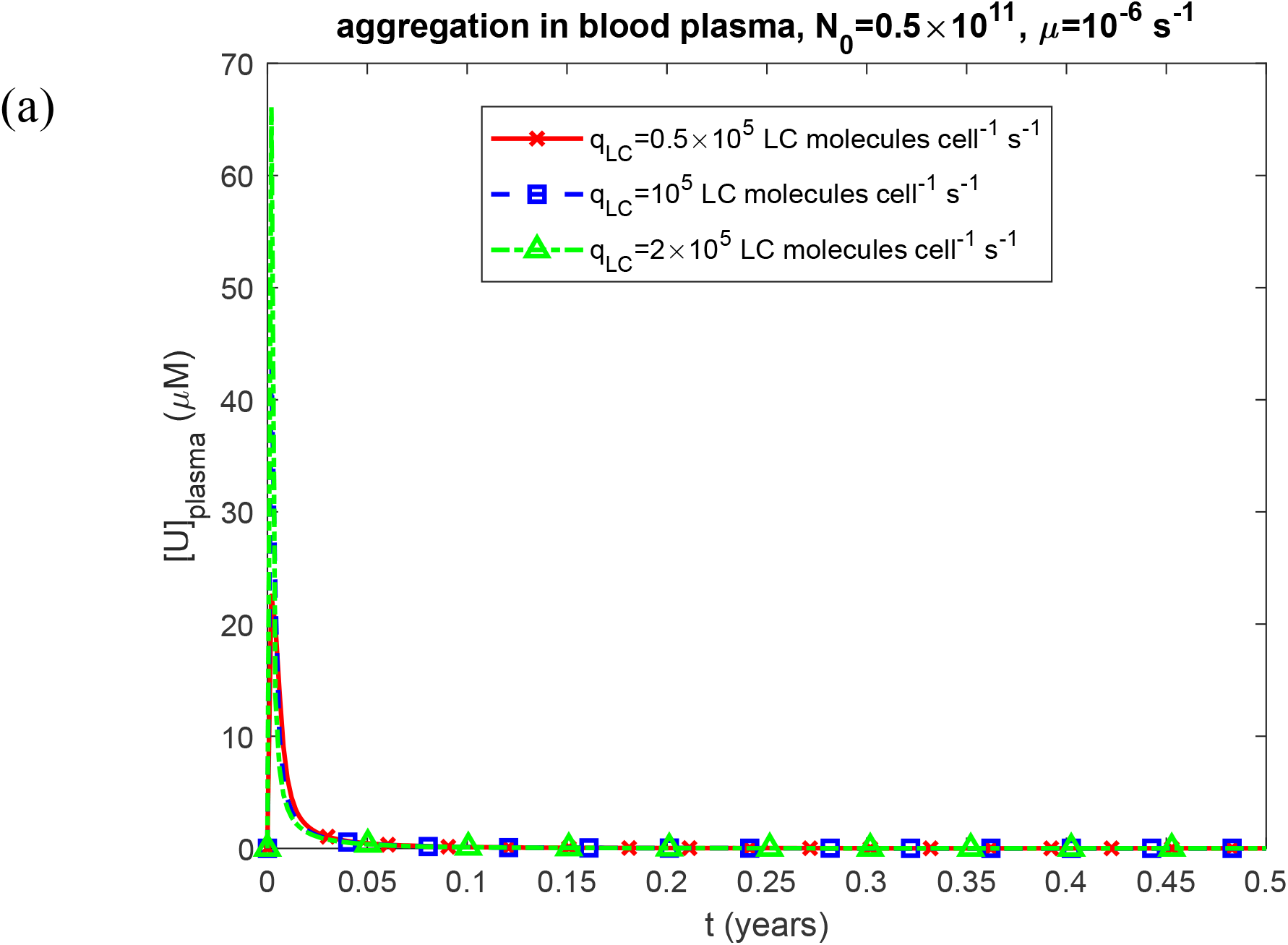
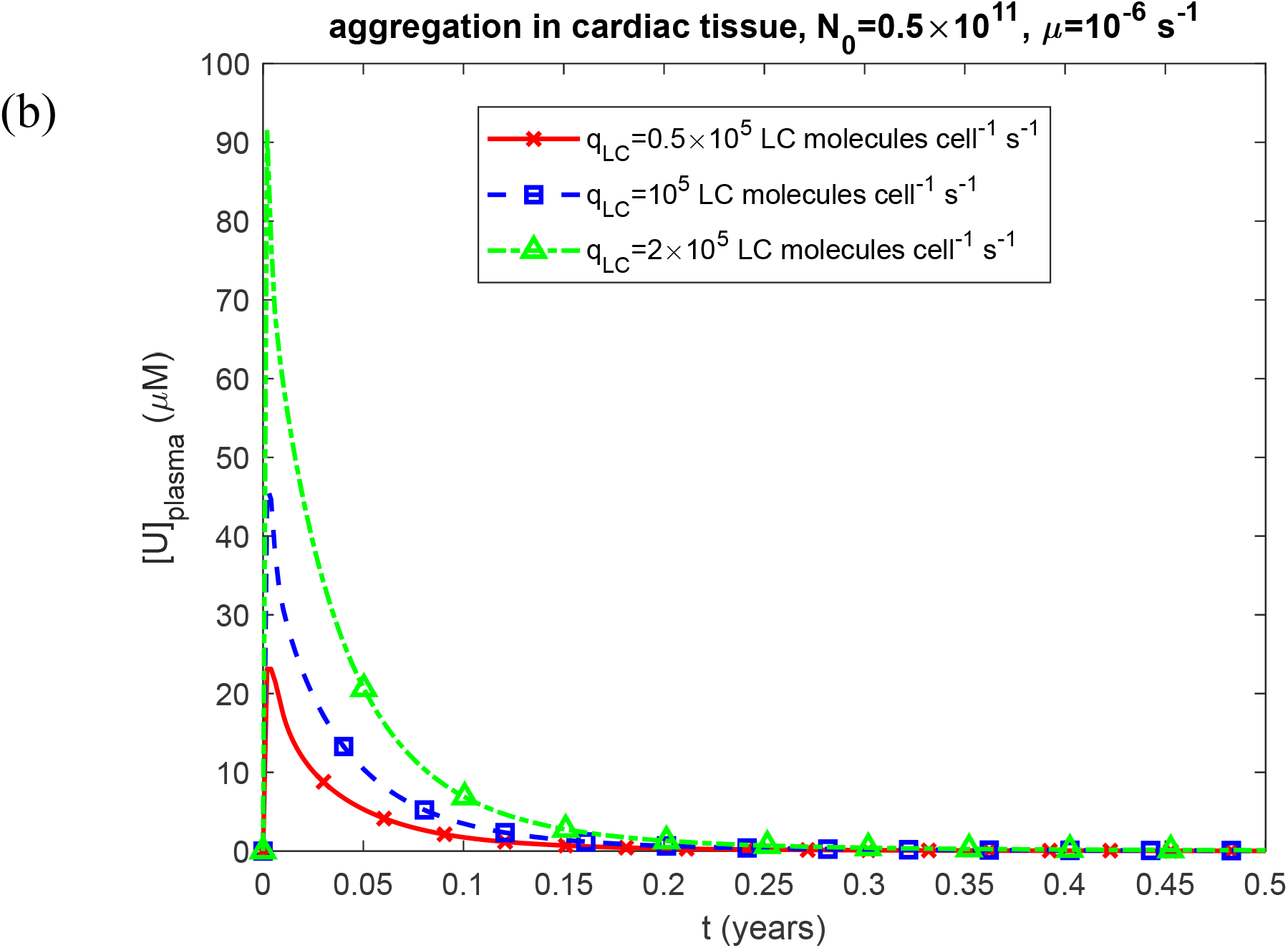
Plasma molar concentration of unfolded Ig LC monomers susceptible to aggregation, [*U*]_*plasma*_, vs time, *t*, for the scenarios of (a) aggregation occurring in the blood plasma and (b) aggregation occurring within the cardiac tissue. Results are shown for three different values of the LC secretion rate per clonal plasma cell, *q*_*LC*_ . Computations were performed for *k*_1_ =10^−6^ s^-1^, *k*_2_ = 10^−6^ *μ*M^-1^ s^-1^, *k*_*u*_ = 4 ×10^−4^ s^-1^, *θ*_1/ 2, *B*_ = 10^7^ s, and *θ*_1/ 2, *I*_ = 10^7^ s. Results correspond to simulations with anticancer therapy.

**Fig. S29.**
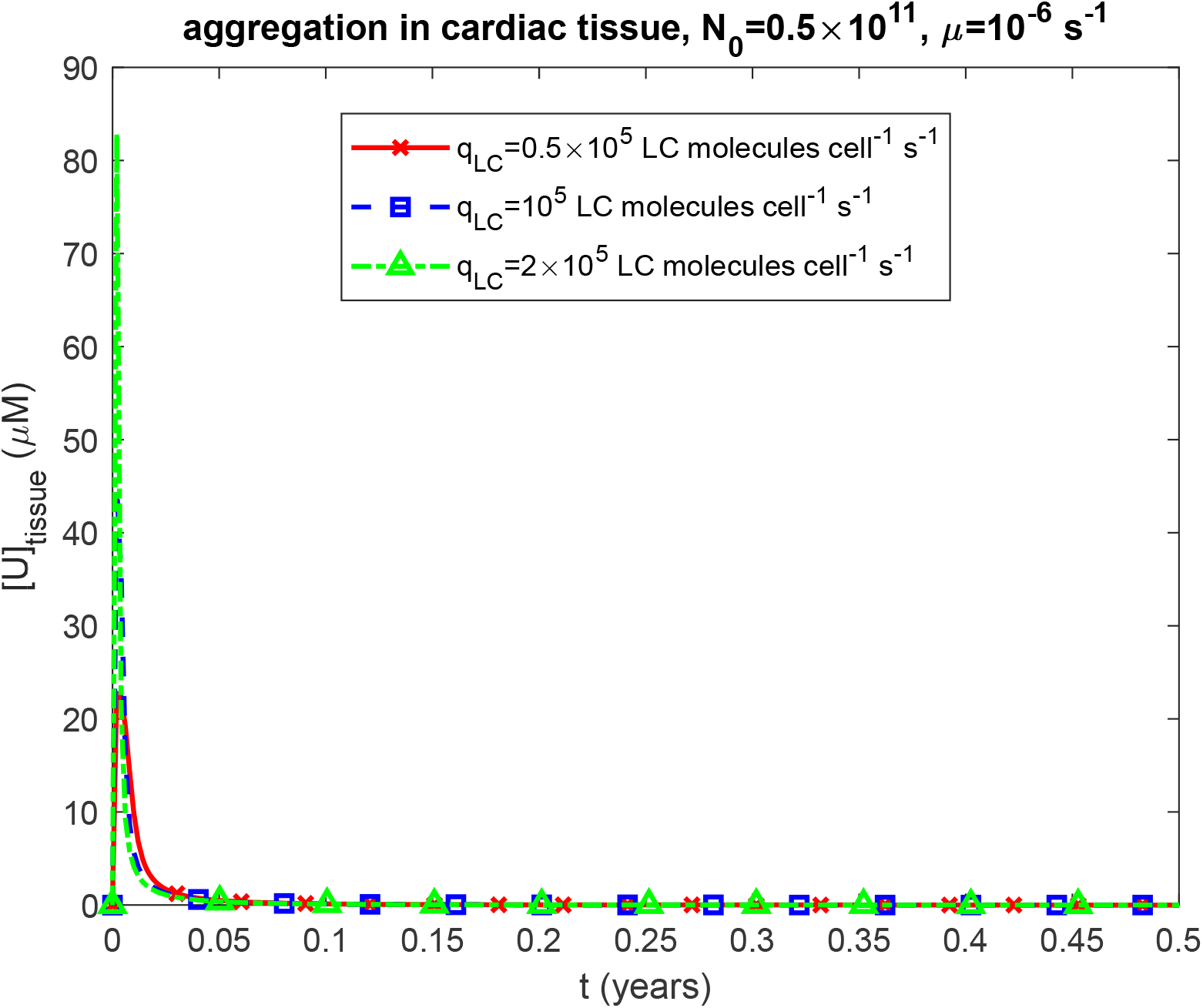
Molar concentration of unfolded Ig LC monomers in cardiac tissue, [*U*]_*tissue*_, vs time, *t*, for the scenario where aggregation occurs within the tissue. Results are shown for three different values of the LC secretion rate per clonal plasma cell, *q*_*LC*_ . Computations were performed for *k*_1_ =10^−6^ s^-1^, *k*_2_ = 10^−6^ *μ*M^-1^ s^-1^, *k*_*u*_ = 4 ×10^−4^ s^-1^, and *θ* _1/ 2, *B*_ = 10^7^ s. Results correspond to simulations with anticancer therapy.

**Fig. S30.**
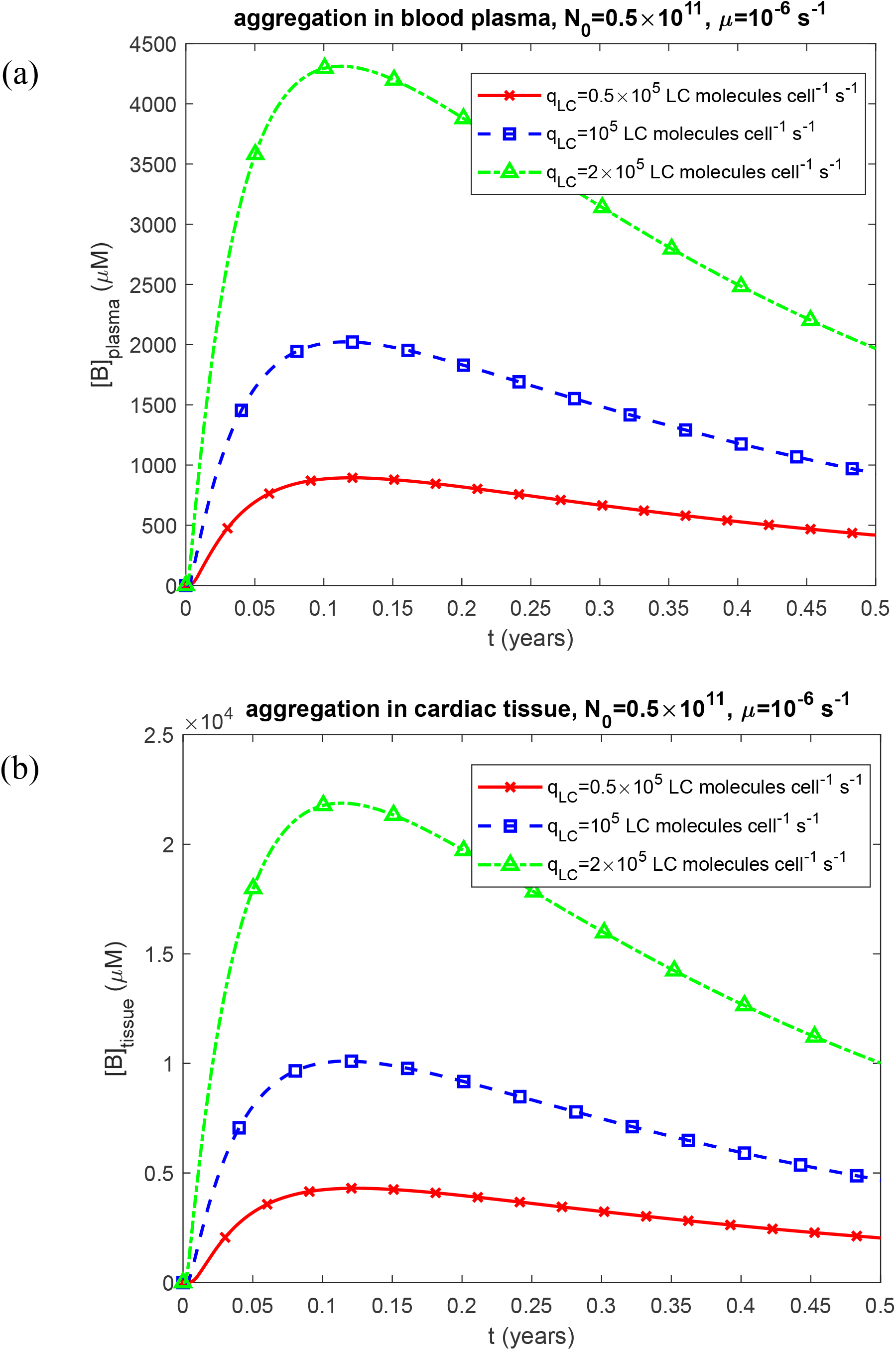
(a) Molar concentration of free LC oligomers in blood plasma, [*B*]_*plasma*_, vs time, *t*, when aggregation occurs in the plasma, and (b) molar concentration of free LC oligomers in cardiac tissue, [*B*]_*tissue*_, vs time, *t*, when aggregation occurs within the tissue. Results are shown for three different values of the LC secretion rate per clonal plasma cell, *q*_*LC*_ . Computations were performed for *k*_1_ =10^−6^ s^-1^, *K*_2_ = 10^−6^ *μ*M^-1^ s^-1^, *k*_*u*_ = 4 ×10^−4^ s^-1^, *θ* _1/ 2, *B*_ = 10^7^ s, and *θ*_1/ 2, *I*_ = 10^7^ s. Results correspond to simulations with anticancer therapy.

**Fig. S31.**
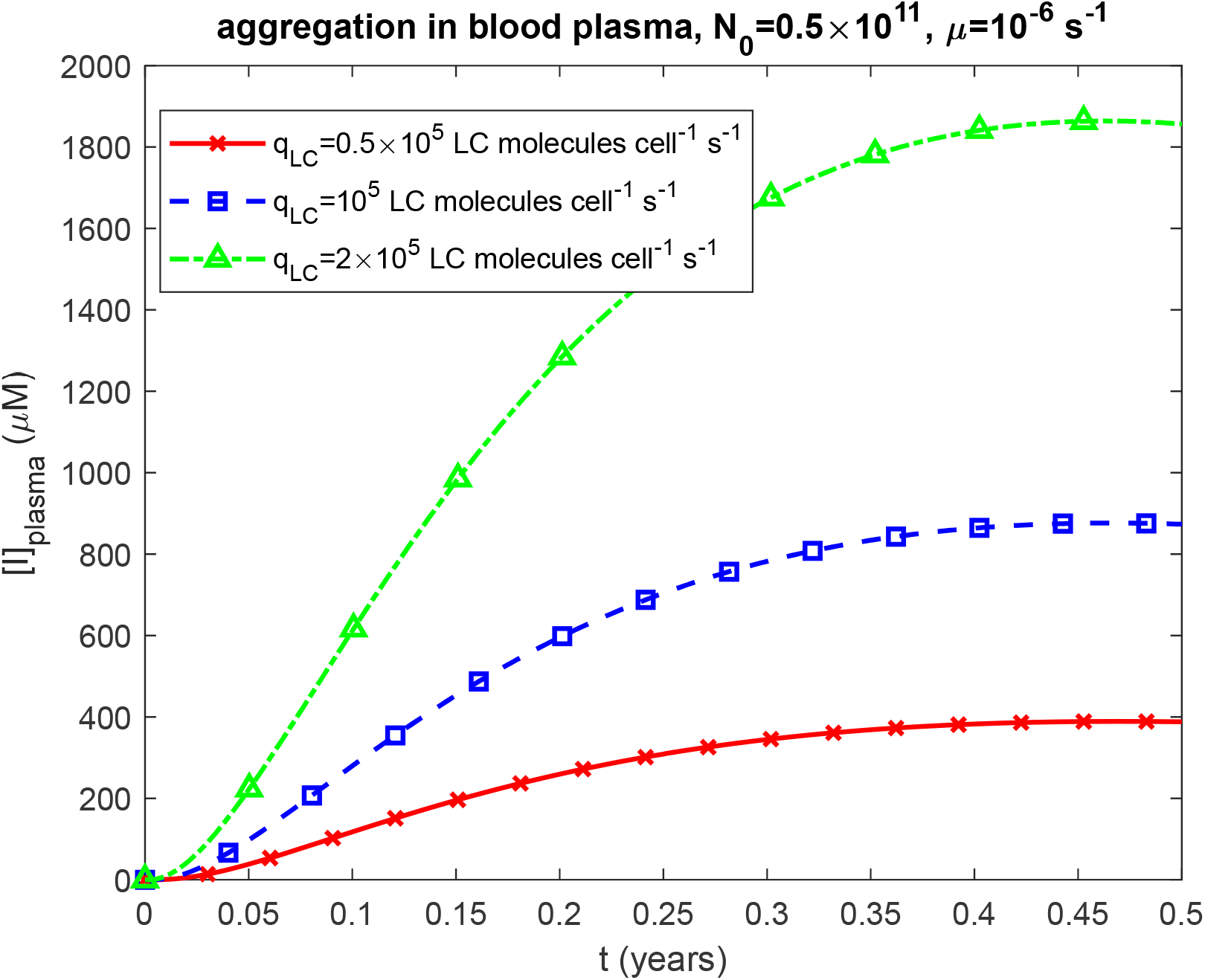
Molar concentration of LC oligomers deposited into amyloid protofibrils circulating within blood plasma, [*I*]_*plasma*_, vs time, *t*. Results are shown for three different values of the LC secretion rate per clonal plasma cell, *q*_*LC*_ . Computations were performed for *k*_1_ =10^−6^ s^-1^, *k*_2_ = 10^−6^ *μ*M^-1^ s^-1^, *k*_*u*_ = 4 ×10^−4^ s^-1^, *θ*_1/ 2, *B*_ = 10^7^ s, and *θ*_1/ 2, *I*_ = 10^7^ s. Results correspond to simulations with anticancer therapy.

**Fig. S32.**
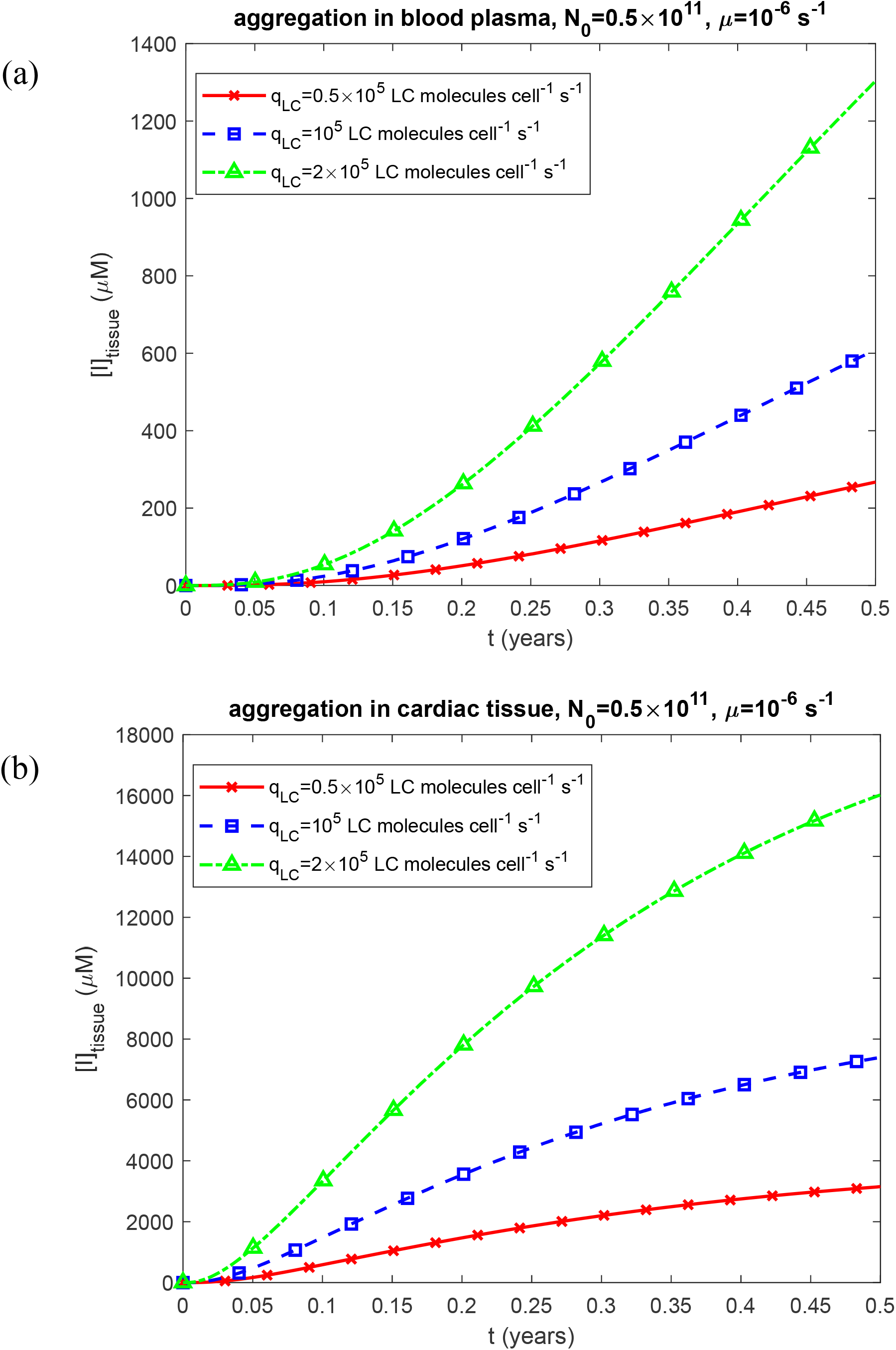
Molar concentration of LC oligomers incorporated into amyloid fibrils infiltrating the cardiac tissue, [*I*]_*tissue*_, vs time, *t*, for two scenarios: (a) aggregation occurring in the blood plasma and (b) aggregation occurring within the cardiac tissue. Results are shown for three different values of the LC secretion rate per clonal plasma celltion, *q*_*LC*_ . Computations were performed for *k*_1_ =10^−6^ s^-1^, *k*_2_ = 10^−6^ *μ*M^-1^ s^-1^, *k*_*u*_ = 4 ×10^−4^ s^-1^, *θ*_1/ 2, *B*_ = 10^7^ s, and *θ*_1/ 2, *I*_ = 10^7^ s. Results correspond to simulations with anticancer therapy.

**Fig. S33.**
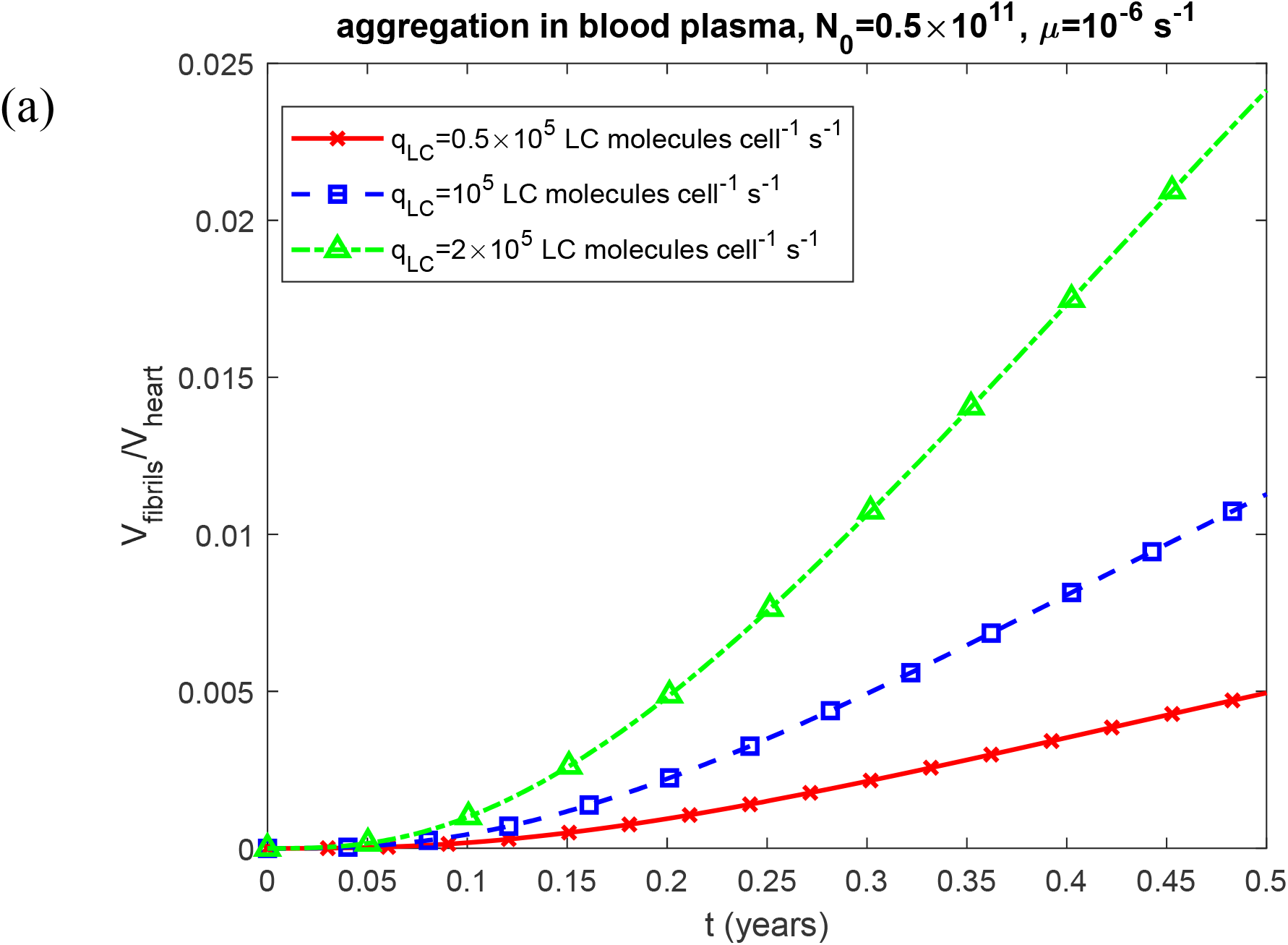
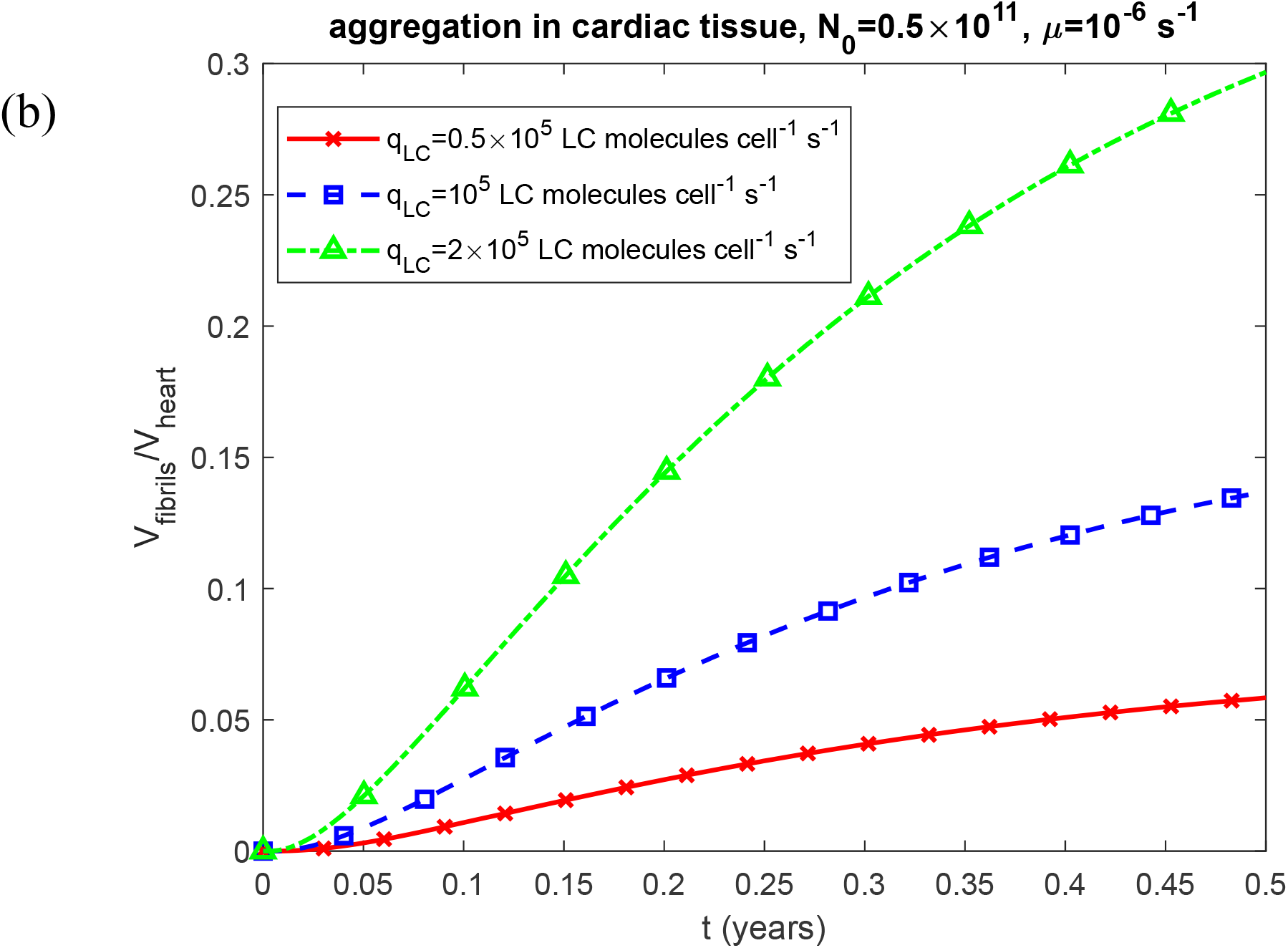
Fraction of myocardial volume occupied by LC fibrils, *V*_*fibrils*_ / *V*_*heart*_, vs time, *t*, for two scenarios: (a) aggregation occurring in the blood plasma and (b) aggregation occurring within the cardiac tissue. Results are presented for three different values of the LC secretion rate per clonal plasma cell, *q* _*LC*_ . Computations were performed for *k*_1_ =10^−6^ s^-1^, *k*_2_ = 10^−6^ *μ*M^-1^ s^-1^, *k*_*u*_ = 4 ×10^−4^ s^-1^, *θ*_1/ 2, *B*_ = 10^7^ s, and *θ*_1/ 2, *I*_ = 10^7^ s. Results correspond to simulations with anticancer therapy.

**Fig. S34.**
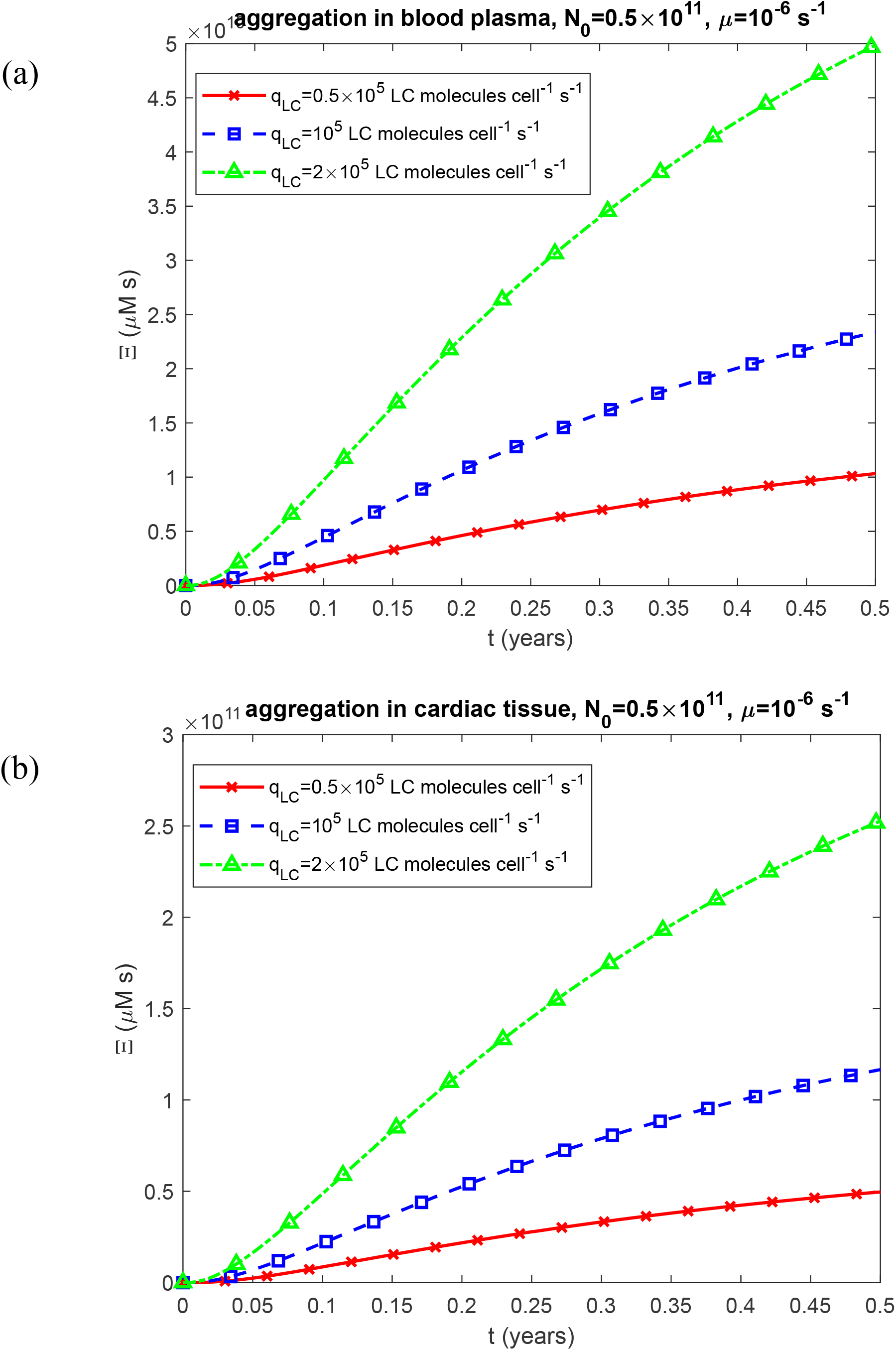
Time-dependent criterion for accumulated cardiotoxicity induced by LC oligomers, Ξ, vs time, *t*, for two scenarios: (a) aggregation occurring in the blood plasma and (b) aggregation occurring within the cardiac tissue. Results are presented for three different values of the LC secretion rate per clonal plasma cell, *q*_*LC*_ . Computations were performed for *k*_1_ =10^−6^ s^-1^, *k*_2_ = 10^−6^ *μ*M^-1^ s^-1^, *k*_*u*_ = 4 ×10^−4^ s^-1^, *θ* _1/ 2, *B*_ = 10^7^ s, and *θ*_1/ 2, *I*_ = 10^7^ s. Results correspond to simulations with anticancer therapy.

**Fig. S35.**
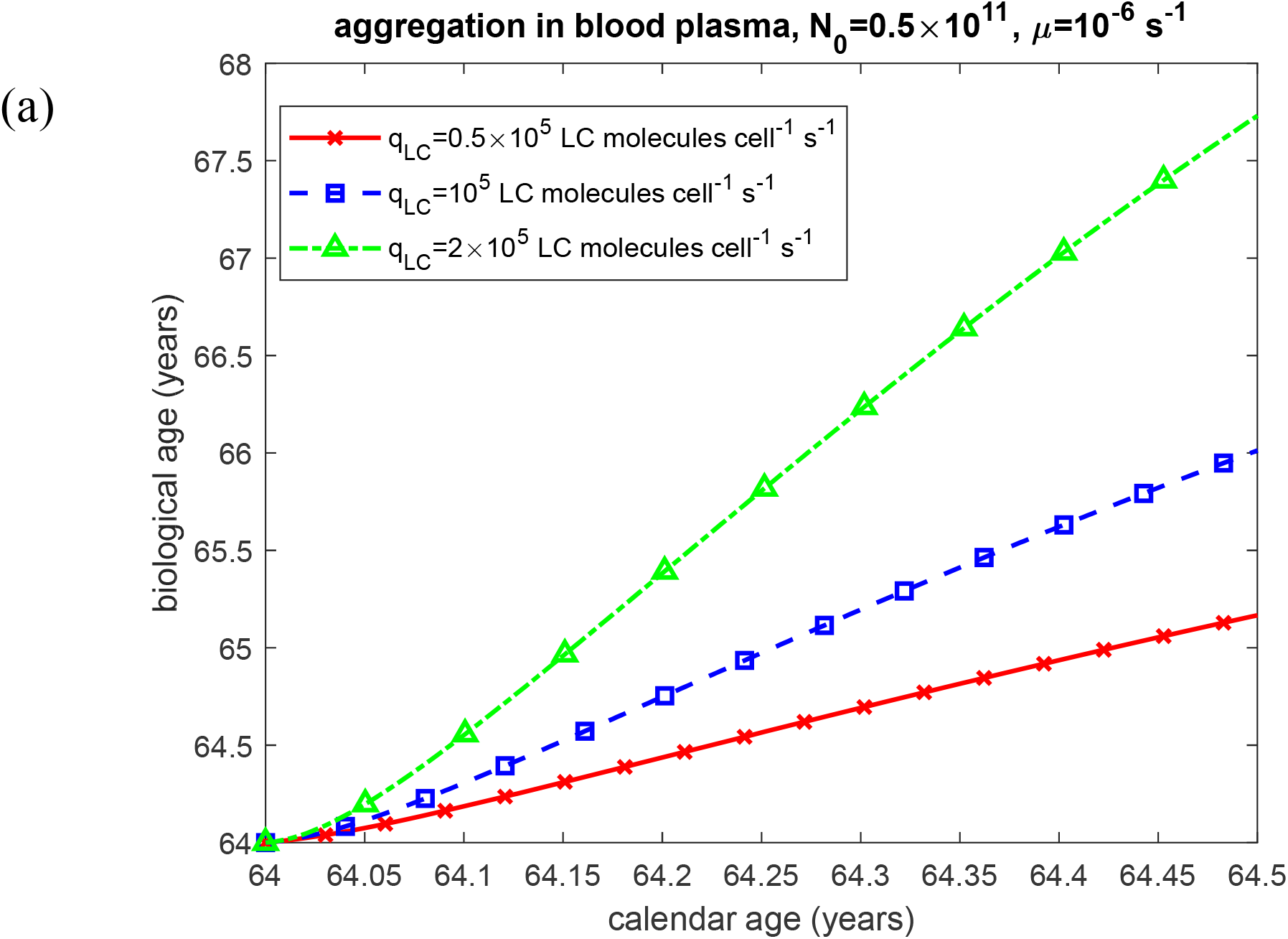
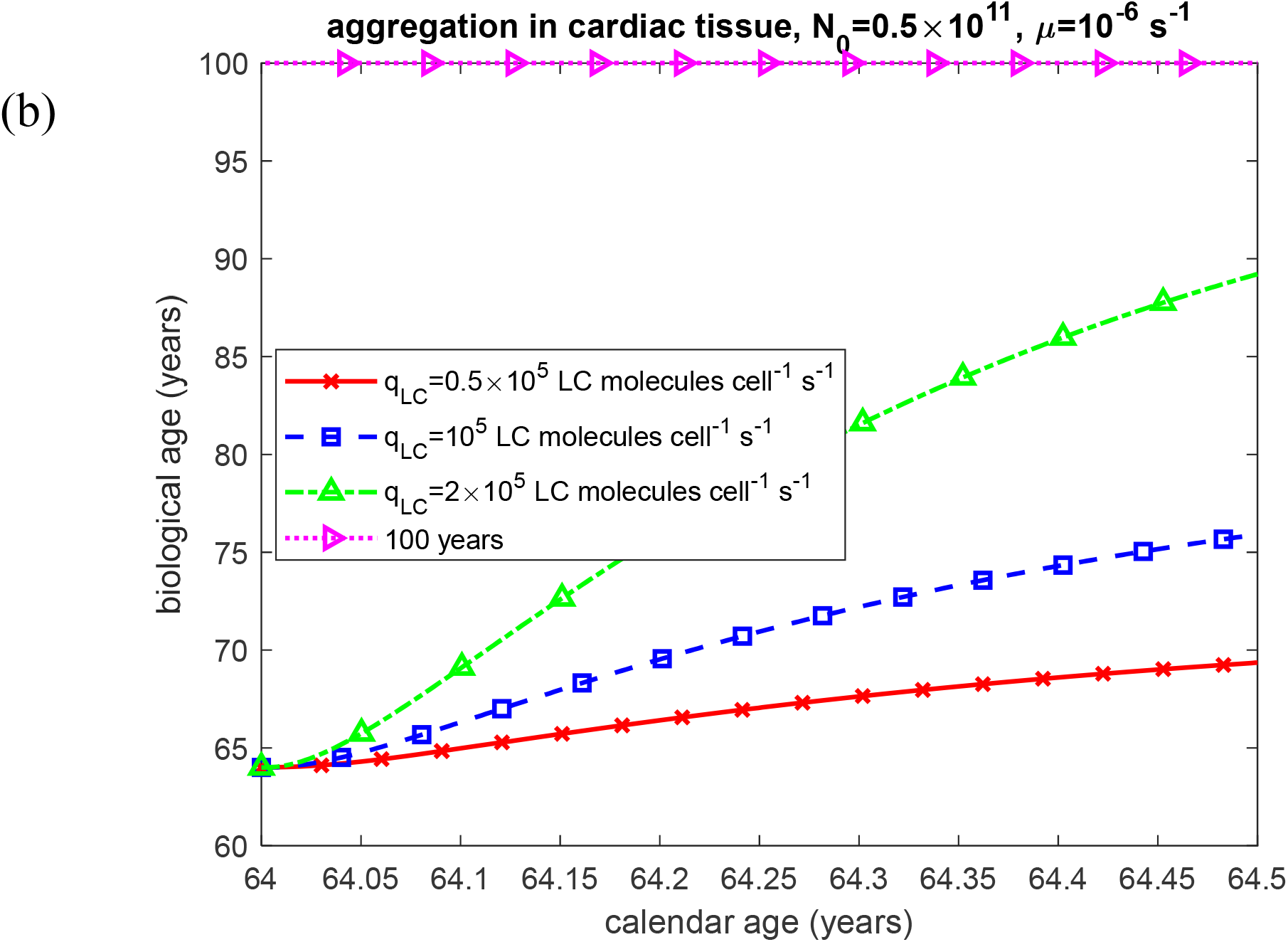
Biological age vs calendar age under two scenarios: (a) aggregation occurring in the blood plasma and (b) aggregation occurring within the cardiac tissue. Results are presented for three different values of the LC secretion rate per clonal plasma cell, *q* _*LC*_. Computations were performed for *k*_1_ =10^−6^ s^-1^, *K*_2_ = 10^−6^ *μ*M^-1^ s^-1^, *k*_*u*_ = 4 ×10^−4^ s^-1^, *θ*_1/ 2, *B*_ = 10^7^ s, and *θ* _1/ 2, *I*_ = 10^7^ s. Results correspond to simulations with anticancer therapy.

